# Mitotic chromosomes are self-entangled and disentangle through a Topoisomerase II-dependent two stage exit from mitosis

**DOI:** 10.1101/2022.10.15.511838

**Authors:** Erica M. Hildebrand, Kirill Polovnikov, Bastiaan Dekker, Yu Liu, Denis L. Lafontaine, A. Nicole Fox, Ying Li, Sergey V. Venev, Leonid Mirny, Job Dekker

**Author notes:** These authors contributed equally (Erica M. Hildebrand, Kirill Polovnikov).

## Abstract

The topological state of chromosomes determines their mechanical properties, dynamics, and function. Recent work indicated that interphase chromosomes are largely free of entanglements. Here, we use Hi-C, polymer simulations and multi-contact 3C, and propose that, in contrast, mitotic chromosomes are self-entangled. We explore how a mitotic self-entangled state is converted into an unentangled interphase state during mitotic exit. Most mitotic entanglements are removed during anaphase/telophase, with remaining ones removed during early G1, in a Topoisomerase II-dependent process. Polymer models suggest a two-stage disentanglement pathway: first, decondensation of mitotic chromosomes with remaining condensin loops produces entropic forces that bias Topoisomerase II activity towards decatenation. At the second stage, the loops are released, and formation of new entanglements is prevented by lower Topoisomerase II activity, allowing the establishment of unentangled and territorial G1 chromosomes. When mitotic entanglements are not removed, in experiment and models, a normal interphase state cannot be acquired.

## INTRODUCTION

Topoisomerase II (Topo II) controls the topological state of the genome ^1^ throughout the cell cycle by catalyzing controlled double strand breaks and allowing DNA duplexes to pass through one another ^1^. There are two forms of Topo II in the human genome, DNA Topoisomerase 2-alpha and DNA Topoisomerase 2-beta, encoded by the *TOP2A* and *TOP2B* genes, respectively ^1–9^. TOP2A has well studied roles in organizing mitotic chromosomes where it is both a structural component and is required for decatenation of sister chromatids at anaphase ^4, 5, 10–20^. TOP2B is expressed throughout the cell cycle and its activity has been detected at open chromatin sites and active chromatin including promoters and CTCF sites during interphase ^1, 21–25^

Whether and how the topological state of chromosomes changes during the cell cycle is not well understood. Hi-C has been widely used to characterize chromosome folding in mitosis and interphase ^26^. However, since Hi-C measures pairwise interactions, one aspect of chromosome folding that is not detected by this method is the entanglement or catenation state of the genome. For the purposes of this study, we define a chromosome entanglement to be a local interlink between two regions of the genome, on the same or different chromosomes. A special type of entanglement operating on rings (loops) is a catenation; catenations can turn a ring into a knotted state or can link two rings (e.g., two chromosome loops). A catenane can also knot a linear chromosome, or a pair of chromosomes, if its ends are sufficiently far away from each other as to behave like a polymer ring. Strand passage facilitated by Topo II can both remove and create entanglements, catenating or decatenating loops. Previous simulations of model polymers showed that highly entangled chains can become highly “intermingled”. The level of intermingling can be detected using multi-contact 3C (MC-3C) ^27^.

MC-3C and super-resolution chromosome tracing data are consistent with interphase chromosomes being largely free of intra-chromosomal entanglements ^27, 28^. Hi-C data also suggest folding of interphase chromosomes into an unentangled polymer state known as the crumpled (fractal) globule ^26, 29, 30^ A recent theoretical study by some of us has revealed a universal behavior of the Hi-C contact probability curve, which is only consistent with the crumpled polymer organization with loops ^31, 32^.

In contrast, the topological state of mitotic chromosomes is less understood. Self-entanglement is supported by in vitro experiments, isolated chromosomes, and some polymer models, with others supporting an unentangled state ^33–36^. In their seminal paper A. Rosa and R. Everaers (2008) propose that the unentangled and territorial interphase state is formed by decompaction from an unentangled mitotic chromosome if Topo II is inactive during exit from mitosis ^35^. Although the assumed absence of Topo II activity during interphase has been challenged by experiments, the Rosa-Everaers model highlights the importance of topological constraints in establishing and maintaining unentangled and territorial chromosomes^35^.

Here we characterized the topological states of mitotic chromosomes and how cells reorganize the topological state of chromosomes upon exit from mitosis. We found that mitotic chromosomes are highly self-entangled. We propose that cells use a two-stage process where Topo II activity eliminates these entanglements upon mitotic exit and prevent the formation of new ones, creating territorial, compartmentalized, and unentangled interphase chromosomes.

## RESULTS

### Topo II inhibition leads to incomplete compartmentalization at G1 entry

Topo II has roles in formation and maintenance of mitotic chromosome structure, as well as in unlinking of sister chromatids ^12, 13, 37^. In contrast, the relevance of Topo II for decondensation of individual chromatids after their separation and upon G1 entry is less established^38^. Here we investigated the genome-wide effect of Topo II chemical inhibition on chromosome folding and topology as cells exit mitosis and enter G1 using Hi-C, imaging, and MC-3C.

To determine whether Topo II activity is required for establishment of G1 chromosome folding, we performed Hi-C 2.0 (Hi-C) on G1 sorted cells from synchronized HeLa S3 cultures during G1 entry (Figure 1A, Figure S1H-I) ^39–41^. Cells were first arrested in prometaphase using a single thymidine block + 12 hours nocodazole arrest (*t* = 0), and then synchronously released into G1 with either DMSO, 30uM ICRF-193, or 30uM ICRF-193 + 200uM Merbarone added at two hours post nocodazole wash-out (*t* = 2 hrs), when at least 50% of cells have entered or passed anaphase (Figure 1B, S1A-F). 30uM ICRF-93 is a sufficient dose to block Topo II activity as it completely prevents sister chromatid decatenation during anaphase and stabilizes Cyclin B when added during release from a mitotic arrest (see methods,^42–44^) (Figure S1A-D). Aliquots were fixed for Hi-C using 1% formaldehyde in early G1 (*t* = 4 hrs) or late G1 (*t* = 8 hrs), and G1 cells were isolated using FACS (Figure S1A, B, E-G, I) ^40, 41^.

**Figure 1:**
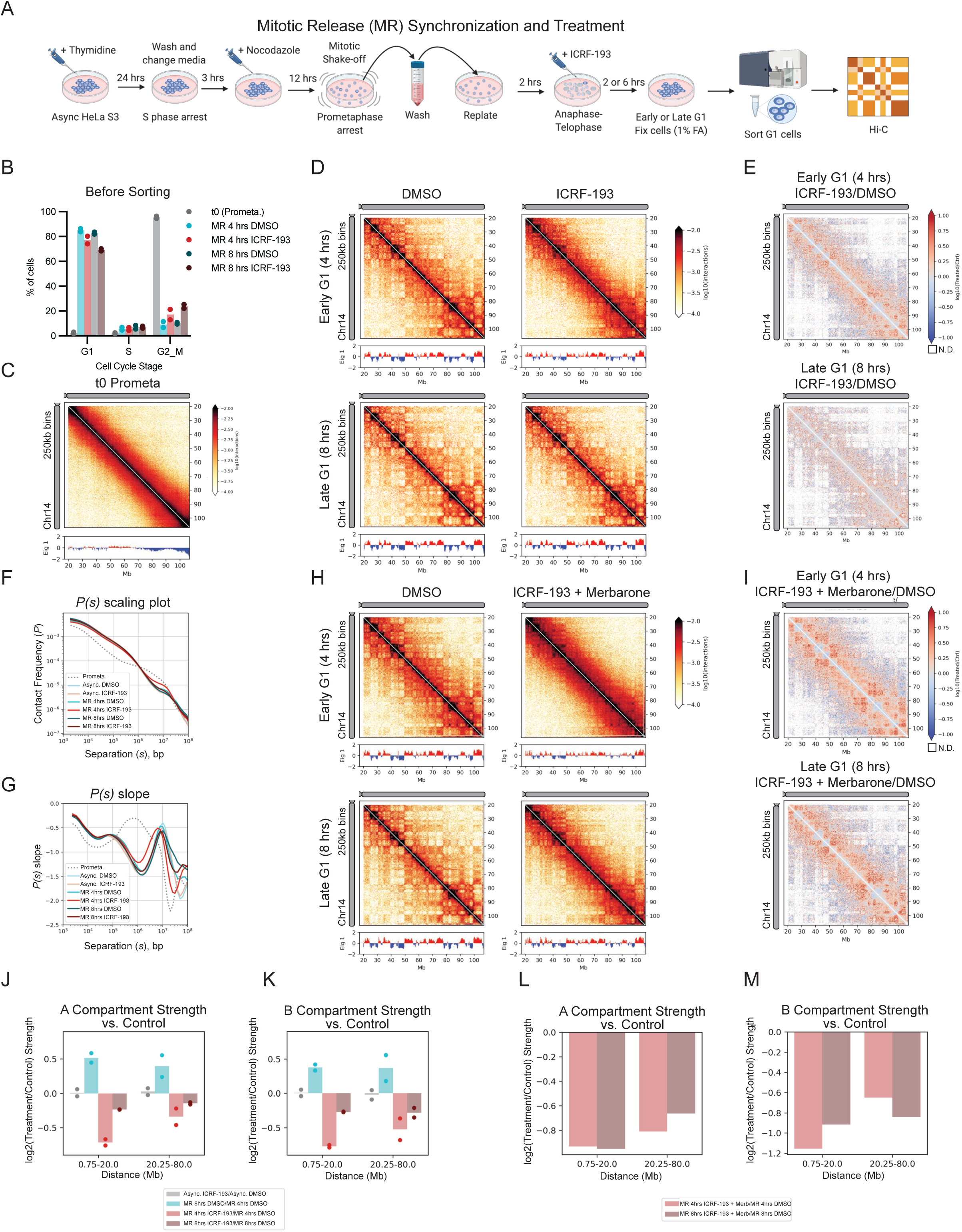
Topo II inhibition by ICRF-193 delays compartment re-establishment at G1 entry. A. Schematic of HeLa S3 mitotic synchronization and release experiment with Topo II inhibition. B. Cell cycle profiles by PI staining and flow cytometry of cells for Hi-C experiment shown in in D-G (N = 2). C. Hi-C interaction heatmap and Eigenvector 1 of unsorted HeLa S3 cells from nocodazole arrest (*t* = 0, prometaphase), from a separate experiment as an example. D. Hi-C interaction heatmaps and Eigenvector 1 of G1 sorted MR HeLa S3 cells treated with DMSO or 30uM ICRF-193, two replicates combined. E. Hi-C interaction log10 ratio heatmap comparing ICRF-193 treatment to DMSO control for each timepoint. F. *P(s)* scaling plot of G1 sorted HeLa S3 cells (Async, MR, and Prometaphase samples from separate experiments). G. First derivative (slope) of *P(s)* scaling plot shown in F. H. Hi-C interaction heatmaps and Eigenvector 1 of G1 sorted MR HeLa S3 cells treated with DMSO or 30uM ICRF-193 + 200uM Merbarone, N = 1. I. Hi-C interaction log10 ratio heatmap comparing ICRF-193 + Merbarone treatment to DMSO control for each timepoint, N = 1. J. AA compartment strength log2 ratio compared to control at each collection time by distance, for G1 sorted HeLa S3 cells treated with ICRF-193. N = 2. (AS and MR from separate experiments, as above). K. BB compartment strength log2 ratio compared to control at each collection time by distance, for G1 sorted HeLa S3 cells treated with ICRF-193, as in J. L. AA compartment strength log2 ratio compared to control at each collection time by distance, for G1 sorted HeLa S3 cells treated with ICRF-193 + Merbarone. N = 1. M. BB compartment strength log2 ratio compared to control at each collection time by distance, for G1 sorted HeLa S3 cells treated with ICRF-193 + Merbarone. N = 1.

In a representative prometaphase Hi-C contact map, a typical mitotic structure is observed, without visible TAD or compartment patterns (Figure 1C). In Hi-C data obtained from control (DMSO treated) cells, the checkerboard pattern representing compartmentalization is first apparent in early G1, and then becomes stronger in late G1, as previously reported (Figure 1D, H) ^40, 45^. In cells treated with ICRF-193 we observed two phenomena: First, at *t* = 4 hrs, we observe a very weak checkerboard and a broad diagonal of enriched interactions reminiscent of mitotic Hi-C maps (Figure 1D, Figure S2G, O) ^40, 44, 46^. Second, at *t* = 8 hrs a nearly normal compartmentalization pattern is observed, indicating that compartments can be established but with delayed kinetics (Figure 1D). Combined addition of ICRF-193 and Merbarone, a Topo II catalytic inhibitor that acts at a different step than ICRF-193, further reduces compartment strength at *t* = 8hrs compared to ICRF-193 alone, although compartments are still somewhat increased compared to the *t* = 4 hrs timepoint (Figure 1H, Figure S1T). Merbarone alone has no effect on compartment strength (Figure S1U).

The log-ratio of Hi-C interactions detected with ICRF-193 vs. DMSO treated cells at *t =* 4 hrs shows enriched interactions close to the diagonal between A and B domains (Figure 1E), similar to what is observed in mitosis (Figure 1C) ^40, 44, 46^. By *t* = 8 hrs, this difference is much smaller, but is retained in ICRF-193 + Merbarone treatment (Figure 1I). In addition, analysis of the relationship of interaction frequency (*P*) of pairs of loci as a function of the genomic distance (*s*) between them shows relatively frequent interactions in early G1 with ICRF-193 treatment for loci separated by 2-20Mb compared to interactions detected in untreated cells (Figure 1 F, Figure S1K, L). By late G1 (*t* = 8 hrs) *P(s)* curves are more similar, although ICRF-193 treatment still shows increased interactions in the 2-20Mb range, and this difference is more pronounced in the ICRF-193 + Merbarone condition (Figure S1V). These changes are readily detectable when the slope of *P(s)* is plotted as a function of genomic separation (*s*) (Figure 1G, Figure S1W). These data suggest that mitotic-enriched interactions are resolved only partially when cells exit mitosis in the presence of Topo II inhibitors. Finally, we quantified compartment strength at different distances. In ICRF-193 and ICRF-193 + Merbarone treated cells, A-A and B-B compartment strength is weaker in early G1 (*t* = 4 hrs) compared to DMSO treated cells, particularly for loci up to 20Mb apart (Figure 1H, I). Compartment strength partially recovers by late G1 in ICRF treated cells, but much less so when cells are treated with both inhibitors (*t* = 8 hrs). We observed no change in chromosome folding at the TAD or loop level upon Topo II inhibition by ICRF-193 (Figure S1M, N). In contrast to the mitotic exit results, Topo II inhibition does not affect steady state intra-chromosomal folding in an asynchronous (Async.) population of mainly interphase cells (Figure S1I-S). We conclude that the interphase conformation can be maintained in the presence of Topo II inhibition. In summary, Topo II activity is required during mitotic exit for complete dissolution of the mitotic state and full establishment of interphase compartments.

### Disruption of compartmentalization with Topo II inhibition can be observed by confocal microscopy

We next tested whether a delay in compartmentalization upon Topo II inhibition could be observed by microscopic analysis of two histone modifications: acetylation of histone H3 lysine 27 (H3K27ac), and trimethylation of H3 lysine 9 (H3K9me3) enriched in the A or B compartments, respectively ^47, 48^. We fixed HeLa S3 cells to coverslips at *t* = 4 hrs and *t* = 8 hrs after mitotic release for confocal microscopy, followed by immunostaining to label H3K9me3 and H3K27ac. We also stained cells with DAPI to mark the DNA. In the DMSO treated cells, the H3K9me3 signal is highest at the periphery, as expected (Figure 2A, Figure S2A, B) ^49–53^. H3K27ac is found in puncta in the interior of the nucleus, as expected for active chromatin (Figure 2A, Figure S2C) ^49, 51–53^. Compared to DMSO treated cells, we find that ICRF-193 treatment during G1 entry significantly increased co-localization of H3K9me3 with H3K27ac regions in early G1, corresponding to the lower compartment strength observed by Hi-C as compared to DMSO treated cells (Figure 2B). In addition, ICRF-193 treatment increases the fraction of H3K27ac signal at the nuclear periphery compared to DMSO treatment at both timepoints (Figure 2C, Figure S2C).

**Figure 2:**
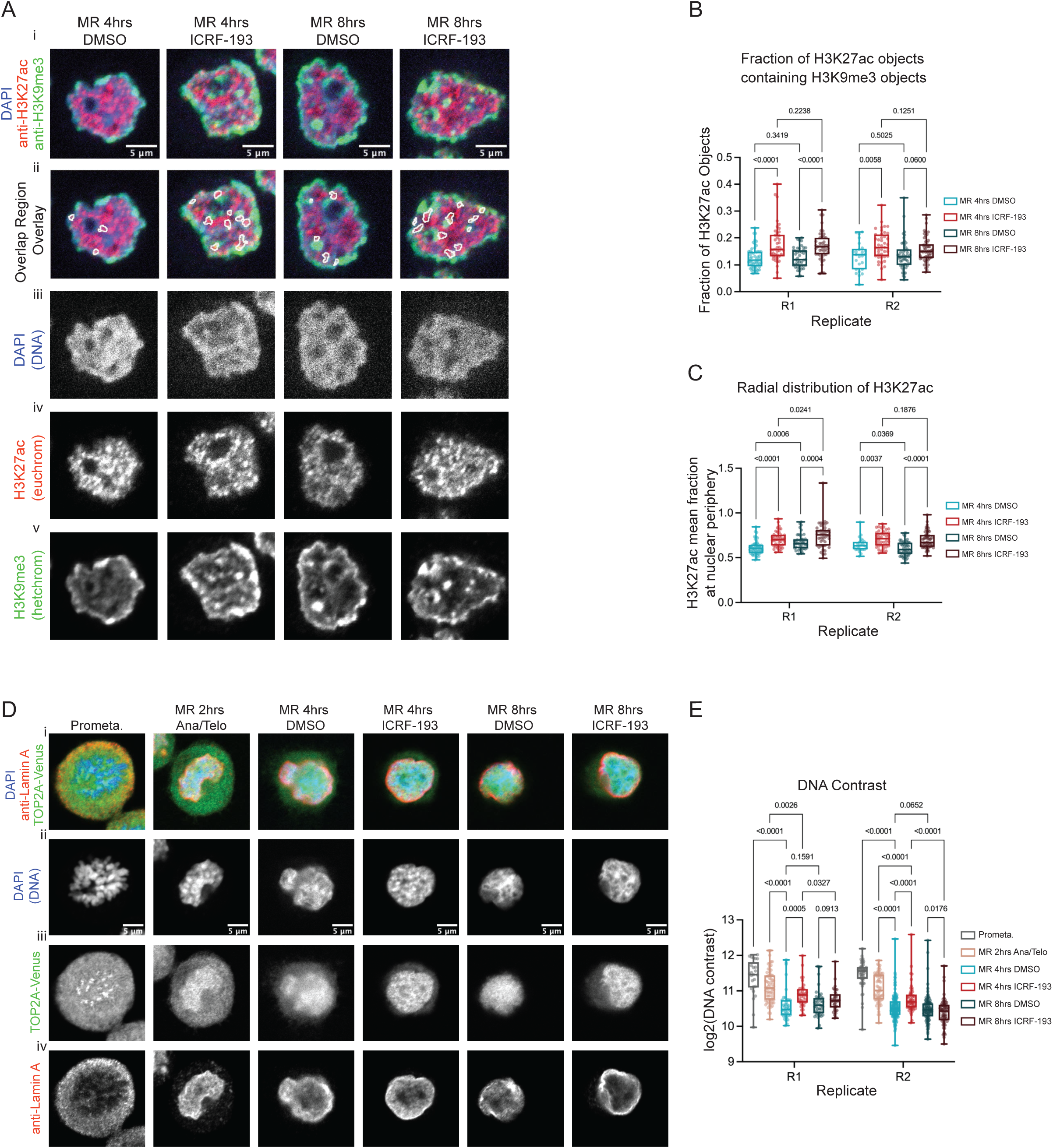
ICRF-193 treatment during G1 entry disrupts nuclear organization and chromosome morphology. A. Representative confocal microscopy images in Early (*t* = 4 hrs) and Late (*t* = 8 hrs) G1 HeLa S3 cells after mitotic release with either DMSO or 30uM ICRF-193 treatment from *t* = 2 hrs post nocodazole washout. i. merge of all channels. Ii. merged image with an overlay (white lines) of the H3K27ac segmented objects containing H3K9me3 objects, as used for quantification of overlap between the two types of chromatin in B. iii. DAPI, iv. H3K27ac, v. H3K9me3. B. Boxplot of the fraction of H3K27ac segmented regions that contain H3K9me3 segmented for each nucleus. C. Boxplot of the mean fraction of H3K27ac signal in the outermost (peripheral) radial bin, out of 10 total bins. D. Representative confocal microscopy images in a mitotic release timecourse with DMSO or ICRF-193 treatment (30uM) starting at *t* = 2 hrs post mitotic release. i. Merged images. ii. DAPI, iii. TOP2A-Venus, iv. Lamin A. E. Boxplot of DAPI signal contrast in each nucleus, at a distance of 10 pixels.

### Topo II inhibition changes DNA morphology and Topo IIA localization in early G1

Individualized mitotic chromosomes display relatively high contrast when stained with DAPI, which we quantified by calculating the contrast of DAPI signal at a 10-pixel distance in HeLa S3 cells with Topo IIA-Venus (Figure 2D, E) ^54^. As cells exit mitosis, this contrast reduces significantly as chromosomes become decondensed. In the presence of ICRF-193 treatment we observe a significantly smaller decrease in DAPI contrast between mitotic exit (starting at *t* = 2 hrs) and early G1 (*t* = 4 hrs) as compared to control cells. Additionally, we find that endogenously tagged Topo IIA-Venus signal has the highest contrast at 2 hrs and decreases as cells enter G1 (Figure S2F). This decrease is not observed in ICRF-193 treated cells. Axial TOP2A staining was observed in some early G1 ICRF-193 treated cells (Figure 2D, see methods).

### Topo II inhibition must occur early during mitotic exit to delay compartment establishment

We next tested whether the increase in compartment strength in late G1 was due to reduced ICRF-193 potency during the six-hour treatment (Figure 3A). To address this, we re-added DMSO or ICRF-193 every two hours throughout the time course and collected cells in late G1 (*t* = 8 hrs) (Figure 3B). These experiments were performed without G1 sorting, resulting in slightly larger number of cells with a G2 DNA content in populations with ICRF-193 treatment (Figure 3C, D). We observe the same phenotype in the late G1 (*t* = 8 hrs) timepoint with ICRF-193 added once or re-added every two hours (Figures 3E-J), therefore the recovery in compartment strength is not due to loss of potency of the inhibitor. Rather, ICRF-193 treatment does not inhibit all Topo II activity, and full inhibition requires use of multiple inhibitors (see above).

**Figure 3:**
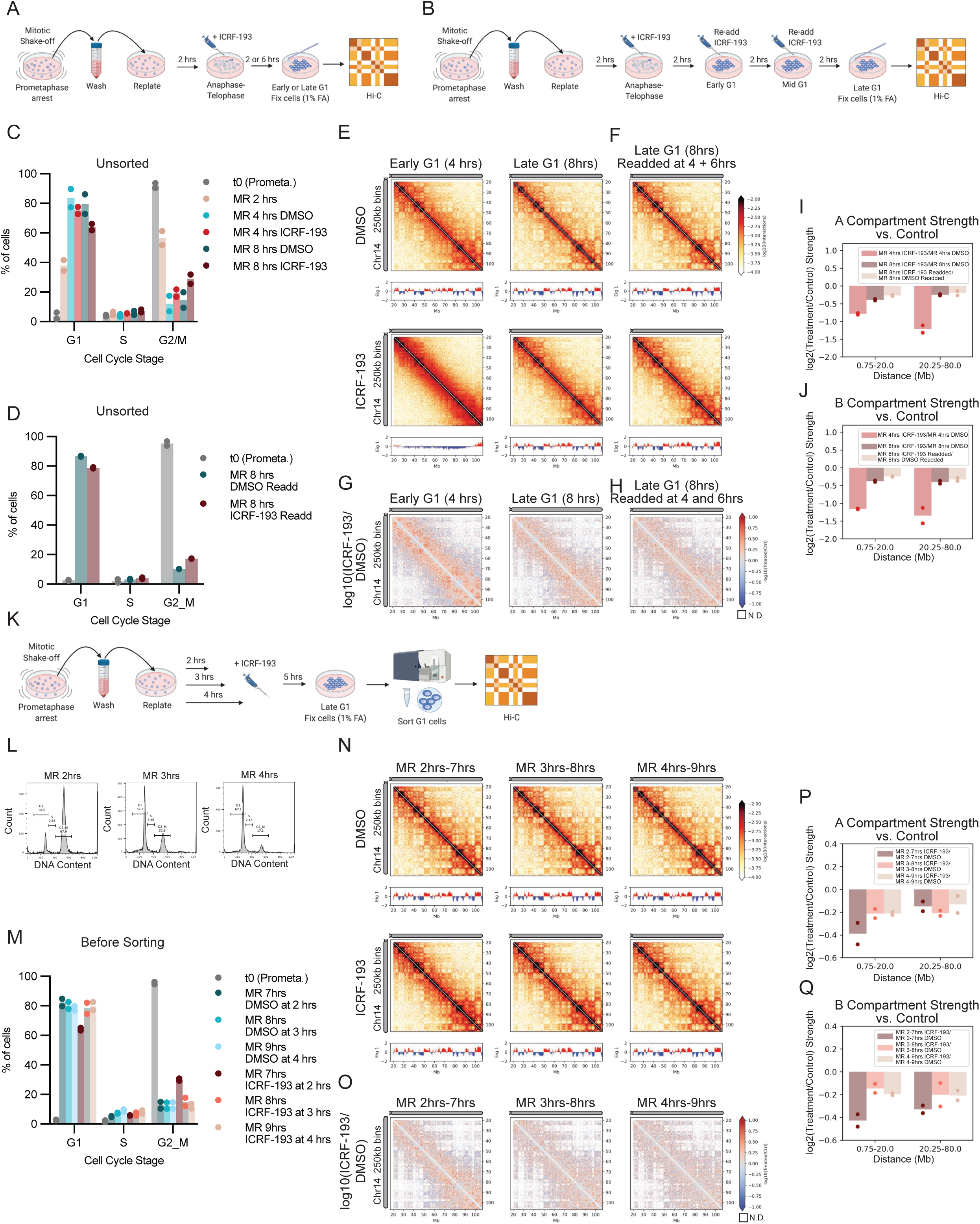
Topo II inhibition must occur during mitotic exit to delay compartment establishment. A. Schematic of HeLa S3 mitotic synchronization and release Hi-C experiment with ICRF-193 treatment starting at *t* = 2 hrs, collected at *t* = 4 hrs (early G1) or *t* = 8 hrs (late G1), without sorting. B. Schematic of HeLa S3 mitotic synchronization and release Hi-C experiment with ICRF-193 treatment starting at *t* = 2 hrs, re-added at *t* = 4 hrs and *t* = 6 hrs, collected at *t* = 8 hrs (late G1), without sorting. C. Cell cycle profiles by PI staining and flow cytometry of cells described in A before G1 sorting. (N = 2). D. Cell cycle profiles by PI staining and flow cytometry of cells described in B before G1 sorting. (N = 2). E. Hi-C interaction heatmaps and Eigenvector 1 of unsorted HeLa S3 cells treated with DMSO, ICRF-193 at *t* = 2 hrs and collected at *t* = 4 hrs or *t* = 8 hrs. Two replicates combined. F. Hi-C interaction heatmaps and Eigenvector 1 of unsorted HeLa S3 cells treated with DMSO or ICRF-193 added at *t* = 2 hrs, and readded at *t* = 4 hrs, and *t* = 6 hrs, and collected at *t* = 8 hrs. Two replicates combined. G. Hi-C interaction log10 ratio heatmap comparing ICRF-193 treatment to DMSO control for each treatment type in E. H. Hi-C interaction log10 ratio heatmap comparing ICRF-193 treatment to DMSO control for samples in F. I. AA Hi-C compartment strength log2 ratio compared to DMSO by distance, separated by compartment type, for HeLa S3 cells described in A and B. N=2. With and without re-adding samples are from separate experiments. J. BB Hi-C compartment strength log2 ratio compared to DMSO by distance, separated by compartment type, for HeLa S3 cells described in A and B. N=2. With and without re-adding samples are from separate experiments. K. Schematic of HeLa S3 mitotic synchronization and release experiment with ICRF-193 treatment starting at *t* = 2 hrs, *t* = 3 hrs, or *t* = 4 hrs post mitotic release, collected after 5 hours of treatment, with sorting for G1 DNA content. L. Flow cytometry profiles for DNA content (PI stain) of synchronized HeLa S3 cells as in I released into G1 at *t* = 2 hrs, *t* = 3 hrs, or *t* = 4 hrs, at the time of ICRF-193 addition. One representative replicate shown. M. Cell cycle profiles by PI staining and flow cytometry of cells described in K before G1 sorting. (N = 2). N. Hi-C interaction heatmaps and Eigenvector 1 of G1 sorted MR HeLa S3 cells treated with DMSO or 30uM ICRF-193 from *t* = 2 hrs to *t* = 7 hrs, *t* = 3 hrs to *t* = 8 hrs, or *t* = 4 hrs to *t* = 9 hrs after mitotic release. Two replicates combined. O. Hi-C interaction log10 ratio heatmap comparing ICRF-193 treatment to DMSO control for each treatment type. Two replicates combined. P. AA Hi-C compartment strength log2 ratio compared to DMSO by distance, separated by compartment type, for HeLa S3 cells described in K. N=2. Q. BB Hi-C compartment strength log2 ratio compared to DMSO by distance, separated by compartment type, for HeLa S3 cells described in K. N = 2.

To determine whether there is a specific transient state in early G1 that requires Topo II activity, we added ICRF-193 at different times post mitotic release (Figure 3K-M). Comparison of the compartment strength in both the A and B compartments between ICRF-193 and DMSO treated cells shows the largest difference for the earliest sample where ICRF-193 was added at *t* = 2 hrs, while ICRF-193 addition at *t* = 3 hrs and *t* = 4 hrs has a reduced effect on compartment establishment (Figure 3 N-Q, Figure S3J-O). Addition of ICRF-193 at *t* = 3hrs or *t* = 4hrs does not prevent further strengthening of compartments in late G1.

### ICRF-193 induced delay in compartment establishment is independent of transcription

Next, we examined whether the structural defects that we observe with Topo II inhibition are related to transcription induced changes in folding, thereby interfering with reformation of G1 structure. We released HeLa S3 cells from mitosis in the presence of Triptolide (TRP) and 5,6-Dichloro-1-beta-Ribo-furanosyl Benzimidazole (DRB), which inhibit transcription initiation and elongation, respectively ^55^. (Figure S3K). By Hi-C, transcription inhibition alone did not result in changes in intra-chromosomal compartment strength at any distance, and transcription inhibition did not change the ICRF-193 phenotype of decreased compartment strength in early G1 (Figure S3 M-Q, Figure S3I-N).

### Mitotic chromosomes are highly intermingled, and become swiftly unmingled during mitotic exit

Previously, we showed using Multi-contact 3C (MC-3C) data that in interphase interacting compartment domains are not extensively intermingled, which is consistent with the genome being decondensed and topologically not entangled, as also inferred from the fractal globule scaling of Hi-C data^27^. We now performed MC-3C for cells in mitosis to determine the extent of intermingling of interacting domains along mitotic chromosomes, how intermingling changes as cells exit mitosis, and how any changes depend on Topo II.

We collected synchronized cells for MC-3C at *t* = 0 (prometaphase arrest), *t* = 2 hrs after nocodazole wash-out (anaphase/telophase), and at *t* = 4 hrs/early G1 or *t* = 8 hrs/late G1 with either DMSO or 30uM ICRF-193 added at *t* = 2 hrs (Figure 4A, Figure S4A; 3 replicates). MC-3C data recapitulate the Hi-C results in terms of differences in compartmentalization and cis/trans ratio between cell cycle states and with ICRF-193 treatment (Figure 4B). Interaction distance distributions for direct pair-wise interactions derived from MC-3C data for *t* = 4 hrs and *t* = 8 hrs DMSO treated cells were similar to previously published MC-3C results in Async. cells, and Hi-C data (Figure 4C, D, Figure S4C-H) ^27^.

**Figure 4:**
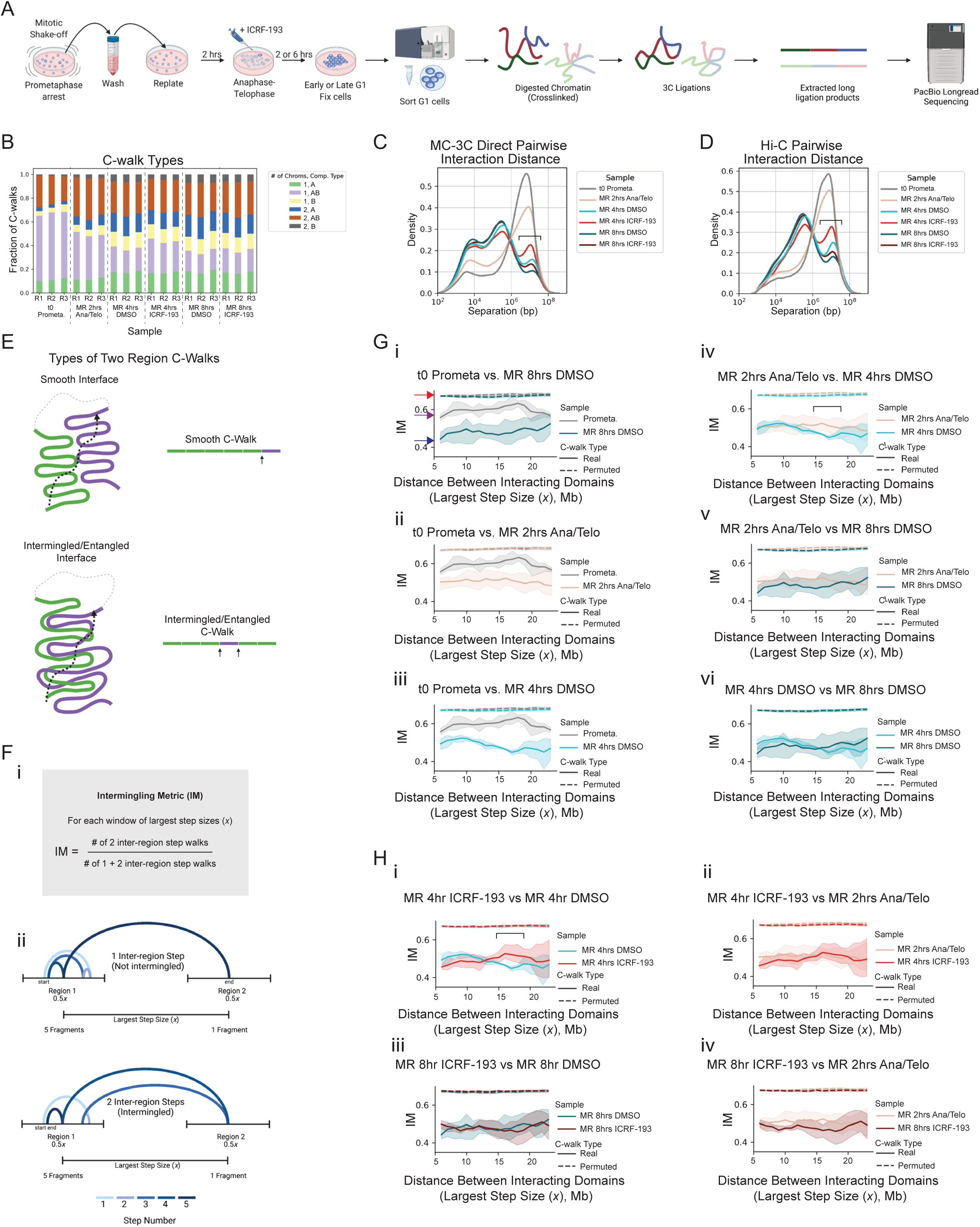
Topo II resolves mitotic entanglements during G1 establishment of interphase chromosome folding. A. Schematic of cell synchronization and MC-3C protocol. MC-3C was performed on HeLa S3 cells during prometaphase arrest (*t* = 0) or after mitotic release at *t* = 2 hrs (anaphase/telophase), *t* = 4 hrs (early G1), or *t* = 8 hrs (late G1). The early and late G1 timepoints had DMSO or 30uM ICRF-193 added at *t* = 2 hrs post mitotic release, and were G1 sorted after fixation. B. Fraction of C-walks within one chromosome or between two chromosomes, and in A, B, or both A and B compartments. Three biological replicates. C. Density plot of the direct pairwise interaction distances from MC-3C C-walks, N = 3. Bracket shows the region where ICRF-193 treatment results in retained mitotic interactions in early G1 compared to DMSO treatment (Figure 1). D. Density plot of pairwise interaction distance from sampled Hi-C libraries made from *t* = 0 prometaphase and *t* = 2 hrs anaphase/telophase unsorted cells, and G1 sorted early and late G1 cells with DMSO or 30uM ICRF-193 treatment. t = 0 and t = 2 hrs samples are from a separate experiment from t = 4 hrs and t = 8 hrs samples. Bracket shows the region where ICRF-193 treatment results in retained mitotic interactions compared in early G1 compared to DMSO treatment. E. Schematic of types of interfaces between different genomic regions on the same chromosome. A smooth interface (top) will result in a C-walk (black dashed arrow) with most steps within each region, and fewer steps between the two regions (indicated by small black solid arrow). An entangled/intermingled interface (bottom) will result in a C-walk (black dashed arrow) with more steps between the two regions (indicated by small black solid arrows). F. i. Formula of the IM calculation for determining how much intermingling/entanglement occurs between two regions. The IM is calculated as the fraction of C-walks with >1 inter-region step. F. ii. Schematic of the two possible types of two region C-walks considered for calculation of the Intermingling Metric (IM). Regions are defined as either side of the largest step (*x*) within each C-walk, with each side extending ¼ of the size of the largest step upstream and downstream of the midpoint of fragment at either end of the largest step (so each region has a maximum size of ½ the largest step size (0.5*x*)). G. Intermingling analysis of control cells during mitotic exit. Pairwise comparisons of the IM at 12Mb window size. Mean (darker line) +/- 95% CI (lighter filled areas) of three biological replicates is shown for the real C-walks. Permuted C-walks (100 permutations per sample x 3 replicates each) are also plotted, with the mean of all 300 permutations for each sample shown as dashed lines, and 95% CI shown by the surrounding filled areas. Arrows in i. indicate low intermingling (blue), medium-high intermingling (purple), and highest intermingling (red). Bracket in iv. indicates the area of significant difference between *t* = 2 hrs and *t* = 4 hrs early G1 DMSO. H. Intermingling analysis of ICRF-193 treated cells, pairwise comparisons to DMSO or t = 2hrs samples plotted as in G. Bracket indicates the area of significant difference between *t* = 4 hrs early G1 DMSO and *t* = 4 hrs early G1 ICRF-193.

MC-3C produces “C-walks”: strings of co-occurring interactions that can provide information on the extent of intermingling between chromosomal regions. Relatively high levels of intermingling can be caused by several factors, including chromatin density, chromosome geometry, but also the presence of topological entanglements^27^. Low levels of intermingling, as we found for interphase cells^27^, are consistent with the decondensed unentangled interphase state. We explored the subset of C-walks that detect interactions between two distal chromosomal domains (see Methods). C-walks at an intermingled surface will include more steps that go back and forth between the two domains as compared to C-walks at non-intermingled surfaces (Figure 4E) ^27^. The order of steps for C-walks across highly intermingled domains approaches that expected for randomized C-walks^27^.

We used MC-3C data to calculate the extent of intermingling by calculating an Intermingling Metric (IM), see Methods and Figure 4 F,G. IM is the fraction of C-walks that transit between two regions of a chromosome more than once. IM can be calculated as a function of genomic distance between the interacting domains (Methods, Figure 4F). We first measured the IM in control (DMSO treated) cells during mitotic exit. Comparing the *t* = 0 prometaphase sample to late G1, we find that prometaphase chromosomes have a higher IM, with ∼60% of C-walks containing more than one step connecting the two regions for most distances between two regions (Figure Gi; fully intermingled domains would be predicted, based on permutation, to have IM=0.67). In late G1, the IM is significantly lower. The effect size observed here is in line with what was observed in simulations of interactions between model polymers with entangled and unentangled interaction surfaces ^27^. This result shows that chromosomes transition from a relatively intermingled to a relatively unmingled state during mitotic exit.

The IM decreases quickly as cells exit mitosis (Figure 4G). At *t* = 2 hrs (consisting of mainly anaphase/telophase cells), the IM is already greatly reduced compared to prometaphase at all distances (Figure 4Gii). At the *t* = 4 hrs early G1 timepoint the IM further decreases compared to *t* = 2 hrs anaphase/telophase, particularly at distances of 12-22 Mb between interacting domains (Figure 4Giii, iv, see bracket). This is the same distance range that has the highest level of the IM in the prometaphase sample. By *t* = 8 hrs (late G1), the IM was at the lowest level (Figure 4G)

### Topo II-dependent loss of intermingling during mitotic exit suggests that mitotic chromosomes are internally catenated

We next investigated the effect of Topo II inhibition on the changes in the IM during mitotic exit. ICRF-193 treatment reduces the loss in the IM in the range of 15-20 Mb, observed from *t* = 2 hrs to *t* = 4 hrs in the DMSO treated samples (Figure 4Hi, ii). This is a similar distance range to the largest loss of compartment strength with ICRF-193 treatment by Hi-C (Figure 1J, K). However, by late G1 the IM at all distances is similar to the DMSO treated sample (Figure 4Hiii, iv), reminiscent of the compartment strength recovery by t = 8 hrs observed by Hi-C with ICRF-193 alone. The later restoration of low intermingling is probably explained by residual Topo II activity not blocked by ICRF-193, as the compartment strength does not fully recover by t = 8 hrs in the ICRF-193+Merbarone double inhibition Hi-C experiments (Fig. 1). Thus, mitotic chromosomes are internally intermingled, and during mitotic exit become decondensed and less intermingled. While high IM in mitosis can reflect, at least in part, the high level of condensation during prometaphase, the dependence of the process of unmingling during mitotic exit on Topo II suggests that decondensation involves resolution of topological entanglements.

### Polymer models of G1 entry without Topo II activity support self-entangled mitotic chromatids

We turned to polymer simulations to directly test the topological state of mitotic chromosomes based on the observed effects of Topo II inhibition by Hi-C and MC-3C. We started simulations with mitotic chromosomes with or without intra-chromosomal catenations, and simulated expansion of 90 Mb chromosomes without strand passage, as in Topo II inhibited cells (see Methods). Simulated interphase organization was then compared to Hi-C results, to determine which initial catenation state is consistent with experiments.

The mitotic chromosome was modelled by a dense array of condensin loops with the average size of 400 kb, corresponding to the size of condensin II loops, that were further confined within a cylinder to reflect chromatin condensation in the mitotic environment ^46^. We considered different topologies of the mitotic chromosome: an “unknotted state”, with loops not catenated with each other, and a “knotted state”, where loops are catenated (see Methods for details). As a global measure of catenations in a mitotic chromosome, we compute a matrix of pairwise catenations (Gaussian linking numbers) between all loop pairs (Figure 5A). While in the knotted state most loops (∼70%, Figure 5A) are catenated with at least one other loop, in the unknotted state less than 3% are. Despite different topologies, the two mitotic states – knotted and unknotted – produce equivalent *P*(*s*) curves (see Figure S5A).

**Figure 5.**
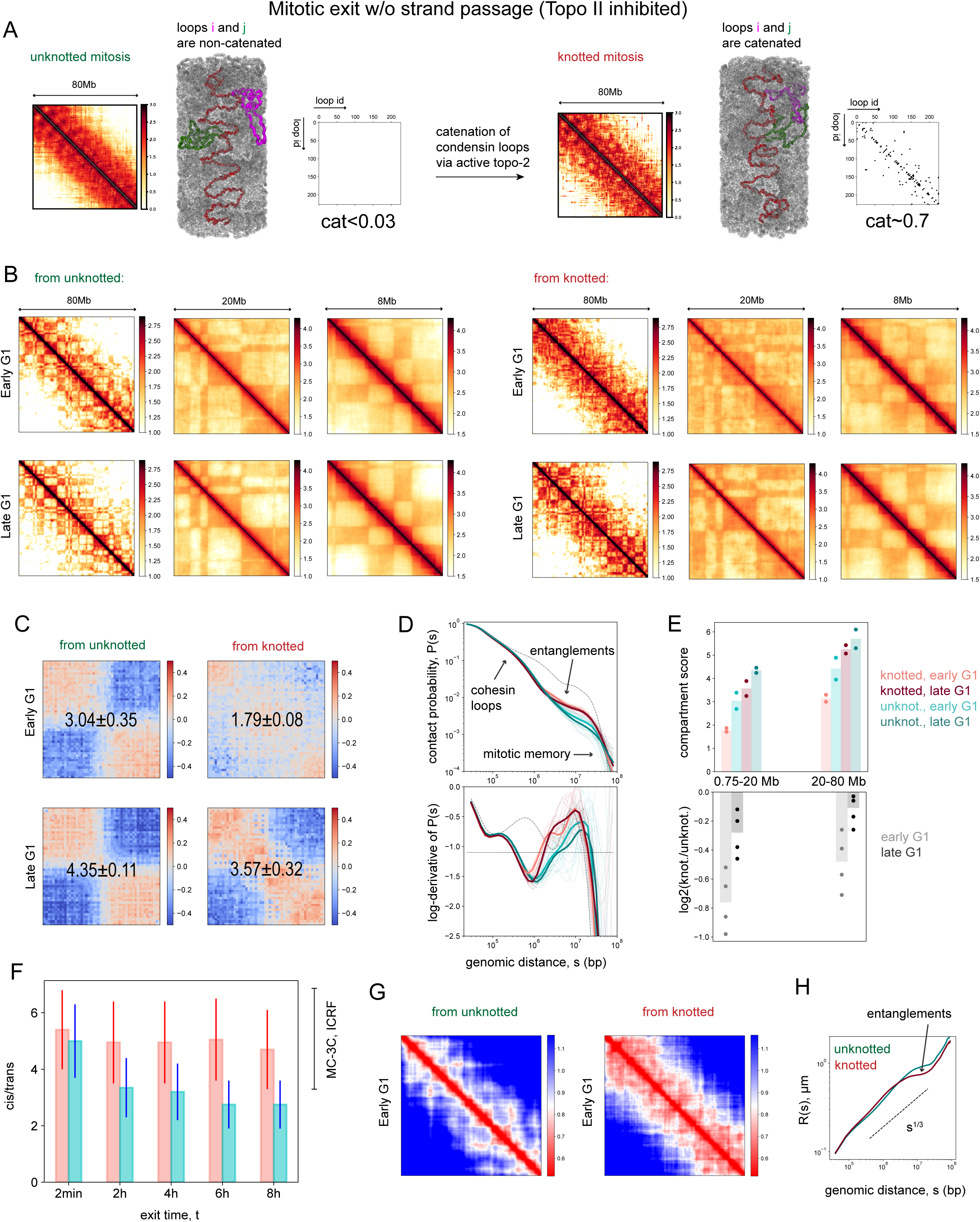
Polymer simulations reveal that hallmarks of mitotic exit with inhibited Topo II correspond to the entangled mitotic chromosome. A. Two topologies of mitotic state considered in polymer simulations. Contact maps and matrices of the Gaussian linking numbers for the loop pairs are shown for each state. The snapshots of the two states are demonstrated: condensin loops (gray) form a dense bottlebrush array; condensins in the loop bases comprise the spiraled backbone (red); two individual loops (non-catenated in the left and catenated in the right) are shown by magenta and green. B. The simulated contact matrices for unknotted (left) and knotted (right) initial states for early (1-2 hour) and late (6-8 hours) interphase. C. The compartmental saddle plots and the corresponding compartmental scores for the contact matrices from panel B at scales 0.75-20Mb are demonstrated. See STAR Methods for more details. D. The contact probability curves and the log-derivatives for the two initial states at different timepoints in G1 as indicated in the legend on the panel E. The gray dashed curve corresponds to mitosis. E. Interphase compartmental scores for the exits from the two mitotic states (top) and the corresponding log2-ratios (bottom) computed for two timepoints in G1, see the legend. Each dot represents a value for the interphase contact map averaged over 16 mitotic replicates and the bars represent the mean value. F. Kinetics of the chromosomal territoriality for the two initial states (knotted – red, unknotted – cyan), measured as cis/trans ratio in simulations. G. Distance maps for the early G1 timepoint out of the two mitotic states. H. The plots of the mean squared end-to-end distances !^!^(#) for the interphase segments of length # obtained from the two mitotic states.

We simulated expansion of mitotic chromosomes by releasing the cylindrical constraints and condensin loops, followed by activation of compartmental interactions and cohesin-mediated loop extrusion (see Methods). We observed very different final states when starting from knotted or unknotted mitotic states. When initiated from an unknotted state, TADs and compartments can form in G1, even without strand passage (Figure 5B, left). In contrast, when started from the knotted state where condensin loops are catenated, mitotic exit without strand passage results in a retained mitotic band of interactions close to the diagonal (Figure 5B, right) and weaker compartmental interactions, while TADs formed similarly to those in the unknotted chromosomes (Figure 5B, C). The retained mitotic band, visible on the simulated interphase contact map formed upon exit from the knotted but not from the unknotted mitotic state, closely resembles Hi-C patterns seen in Topo II inhibition experiments (Figures 5B, S5A-B and Figure 1H). This mitotic band can be also seen as a broad shoulder on the corresponding *P*(*s*) curves (Figure 5D).

Presence of this mitotic band visible during interphase, seen in experiments and in simulations from the knotted mitotic state, reflects retention of mitotic entanglements in the interphase chromosomes. The same is seen in simulations without compartments, indicating that the band is not caused by compartmental interactions (Figure S5B-D). Interestingly, interphase chromosomal conformation emerging from knotted and unknotted mitotic states also have drastically different distance maps (Figure 5G-H), with loci 0.8-2 Mb apart being about 1.5-times closer in space when exiting from the knotted mitotic space. More compact chromosomes are also observed in microscopy when cells exit mitosis in the presence of ICRF-193 (Figure 2E). In the absence of Topo II activity, initial mitotic entanglements cannot be resolved and prevent full opening of chromosomes (Videos S1-S2). This further prevents establishment of long-range interactions between homotypic compartments both at short and large distances (Figure 5С,E), consistent with experiments. Thus, simulations suggest that retention of mitotic-like morphology and weaker compartmentalization upon Topo II inhibition during mitotic exit indicate a highly self-entangled mitotic state.

Further evidence of entanglement in the mitotic state comes from comparison of territoriality of interphase chromosomes between experiments and simulations, quantified by the cis/trans ratio in simulations and MC-3C data (see Methods). Simulations that start from knotted and unknotted mitotic chromosomes, with inhibited strand passage, yield very different cis/trans ratios in the subsequent G1 phase. We calculated cis/trans ratios as a function of time, for the two initial states (Figure 5F). After several hours, the cis/trans ratios of chromosomes expanded from an unknotted mitotic state fall below the range observed experimentally upon ICRF-193 treatment (cis/trans ratio inferred from MC-3C data), see Fig. S6I. At the same time, territoriality of chromosomes expanded from a knotted mitotic state quickly saturates at values close to those observed in the Topo II inhibition experiments. This agreement between experiments and simulations additionally indicates that mitotic chromosomes are knotted.

### Two-stage exit allows chromosomes to transition from an entangled mitotic to an unentangled interphase state

How can chromosomes transition from an entangled mitotic state into a largely unentangled G1 state, while establishing proper interphase organization? Our goal is to reproduce two global features of interphase chromosome organization, starting from entangled mitotic chromosomes: (i) the fractal (crumpled) globule intra-chromosomal organization measured on the *P(s)* curve; and (ii) chromosome territoriality (measured by the cis/trans ratio). The fractal globule, a compact and unknotted polymer state, is evident from the −1… −1.2 slope of the *P*(*s*) curve^26, 56–60^, and is best seen when cohesin-mediated loops do not obscure this scaling, e.g., upon cohesin depletion^61, 62^.

First, we explored a model where entangled mitotic chromosomes simply expand in the presence of high Topo II activity (modeled by having a low barrier to strand passage). In this “one-stage model”, condensin loops are released, which occurs by late telophase^40, 63^, with simultaneous release of cylindrical confinement of the chromosome (Figure 6A-C). Simulations show that while the knotted mitotic state rapidly expands, chromosomes extensively mix both in cis and in trans. Specifically, the fractal globule organization cannot be established as evident from the slope of the *P*(*s*) curve (Figure 6B). To better check for the fractal scaling, we remove cohesin-mediated loops in the late G1 timepoint ^31, 32^. We observe that the slope of the *P*(*s*) curve approaching −1.5, far below the expected −1… −1.2 for the fractal globule. This behavior is expected as high Topo II activity turns the chain into a topologically unconstrained and highly entangled nearly ideal chain (Figure 6B) ^64^. Furthermore, we see that chromosomal territoriality falls below the levels observed in MC-3C for WT interphase cells (Figure 6C and Figure S6I). Together, these results indicate that while Topo II activity allows entangled mitotic chromosomes to expand, it precludes establishment of chromosome territories and formation of the fractal globule state, as generally expected for topologically unconstrained polymers^35, 64^.

**Figure 6.**
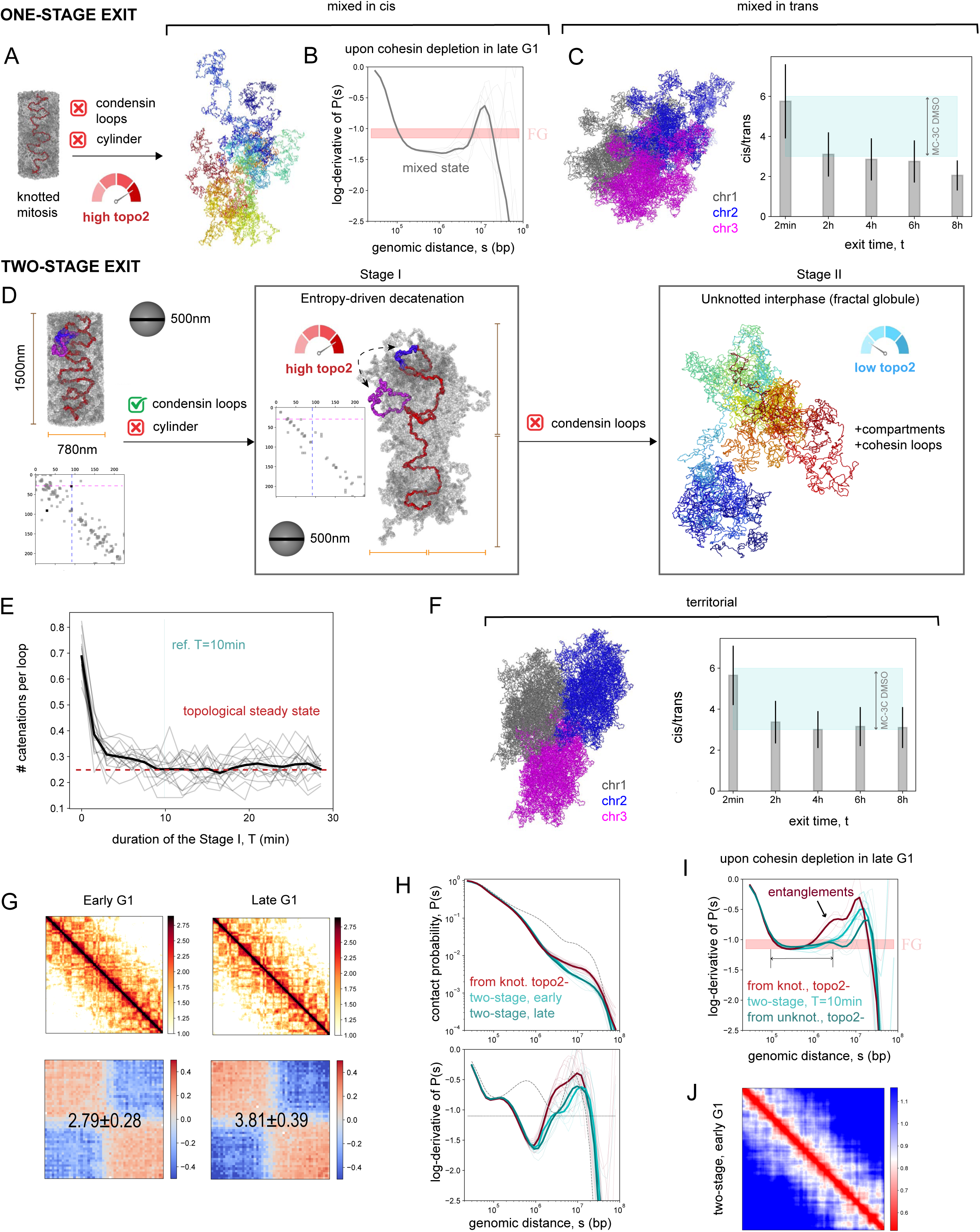
The two-stage model of mitotic exit allows for directed Topo II and removal of most of the mitotic catenations. (A-C): one-stage exit. (D-J): two-stage exit. A. Schematic for the one-stage exit.. Such an exit results in internal mixing of the chains, as shown by the disordered organization of the colored interphase chain on the right. B. The log-derivative of the average contact probability *P*(*s*) computed after removal of cohesin loops at the late G1 timepoint (bold gray curve; different replicates are shown by thin gray lines). The experimental range of the *P*(*s*) slopes between −1.15 and −1, corresponding to the fractal globule (FG) state, is shown by pink. C. Snapshots of three overlapping chromosomes (PBC images) from one-stage simulations (left). The cis/trans ratio as the function of the exit time (right). D. Schematic for the two-stage exit. The matrices of the linking number for the condensin loop pairs are shown in the knotted mitotic state and by the end of Stage I; magenta and blue dashed lines correspond to the two loops that are catenated in the mitosis, but decatenate during Stage I. E. Mean number of catenations per loop as a function of duration of the first stage (bold black curve; 16 replicates are shown by thin gray lines). The red dashed line denotes the steady-state number of residual catenations ∼0.25. This steady-state is achieved in *T* ≈ 10 minutes, which is used in further analysis of the two-stage exit. F. Snapshots of three weakly overlapping chromosomes (PBC images) from two-stage simulations (left). The cis/trans ratio as the function of the exit time (right). G. Interphase contact maps for early and late timepoints, obtained via the two-stage exit (top). The corresponding compartmental saddle plots with the compartmental scores are shown in the bottom. H. The interphase *P*(*s*) and its log-derivative for the two-stage exit at two timepoints (cyan) and for the exit with inhibited strand passage from the knotted initial state (red). The dashed black curves correspond to the mitotic state. I. The log-derivative of *P*(*s*) computed after removal of cohesin loops at the late G1 timepoint. The curves corresponding to the two-stage exit is shown by cyan, while the curves for the exit with inhibited strand passage from knotted and unknotted states are shown by red and dark cyan, correspondingly. The experimental range of the *P*(*s*) slopes between −1.15 and −1, corresponding to the fractal globule (FG) state, is shown by pink. J. Distance map for the early G1 timepoint obtained via the two-stage process.

One-stage exit from a knotted mitotic state with Topo II activity does not reproduce the features of an unentangled interphase. In fact, while Topo II activity can allow expansion of knotted chains, its continued activity leads to mixing and does not lead to formation of the unentangled interphase. Thus, we seek a mechanism that can efficiently disentangle the mitotic state and then maintain this unentangled state later through interphase. Our key idea is that keeping mitotic loops while allowing chromosomes to decondense could entropically bias Topo II towards decatenation of the loops. This could lead to formation of the sought unentangled state, that then needs to be maintained through the rest of expansion.

On the basis of this idea, we developed a two-stage expansion process (Figure 6D). During Stage I, Topo II is active and the cylindrical constraints on the mitotic chromosome are released, while mitotic loops are still present, i.e., the nuclear environment/chromatin changes to their interphase state, but condensin loops remain. Simulations show that the first stage results in directed decatenation of condensin loops (Figure 6D-E); this is quantified using the matrices of linking numbers between the loops. A simulated chromosome is reminiscent of a swollen bottlebrush, which gradually lengthens, as more and more loops become decatenated from each other. In 2 minutes of the first stage around 50% of mitotic catenations are removed, while at T=10 minutes after mitosis the average amount of catenations per one loop reduces almost 3-fold and reaches the topological steady-state, in which the number of newly formed catenations matches the number of removed catenations (Figure 6E). Temporal activation of Topo II activity (or delayed inhibition) for 10-20% of duration of Stage I results in rapid relaxation of catenations, with associated increase of G1 compartmental strength and decrease of territoriality (Figure S5H-J). Importantly, inducing active Topo II inside the mitotic chromosome before it starts expanding (i.e., while in the cylinder) is not able to decatenate the loops, indicating that disentanglement requires Topo II activity during expansion with intact loops (Figure S6A). Thus, directed expansion of the chains during the first stage drives repulsion between the loops, and Topo II activity mediates loop-loop decatenation.

At Stage II, the mitotic loops are released, and Topo II activity is significantly decreased. It starts with already unentangled chromosomes and maintains this state during their further expansion. Active cohesin-mediated loop extrusion and compartmentalization are also introduced at Stage II (Figure 6D). Simulations show that the remaining level of catenations from Stage I is negligible, and chromosomes form the fractal globule with a characteristic slope of *P*(*s*), clearly seen upon depletion of cohesin (Figure 6I, 7F) ^32, 56, 57, 59–62, 65^. The fractal globule is also evident from the visual comparison of snapshots of chromosomes colored along the chain (Figure 6D and 6A). The fractal globule is known to produce clear “intra-chromosomal territoriality” of genomic segments within a chromosome, as seen for the two-stage exit, in contrast to mixing of segments in the case of the one-stage exit ^26^. Simulations also show the fractal globule state can be formed even in the presence of some low-level of Topo II activity in Stage II (occasional strand passage) (Figure 7F). Furthermore, we find that the territoriality of simulated chromosomes after the two-stage exit agrees with that of experimental interphase MC-3C values, i.e., overlaps with the cis/trans range for chromosomes in DMSO treated cells (Figure 6F). We note that territoriality is difficult to achieve in the one stage model unless the Topo II timing is precisely fine-tuned (Figure S6G).

**Figure 7:**
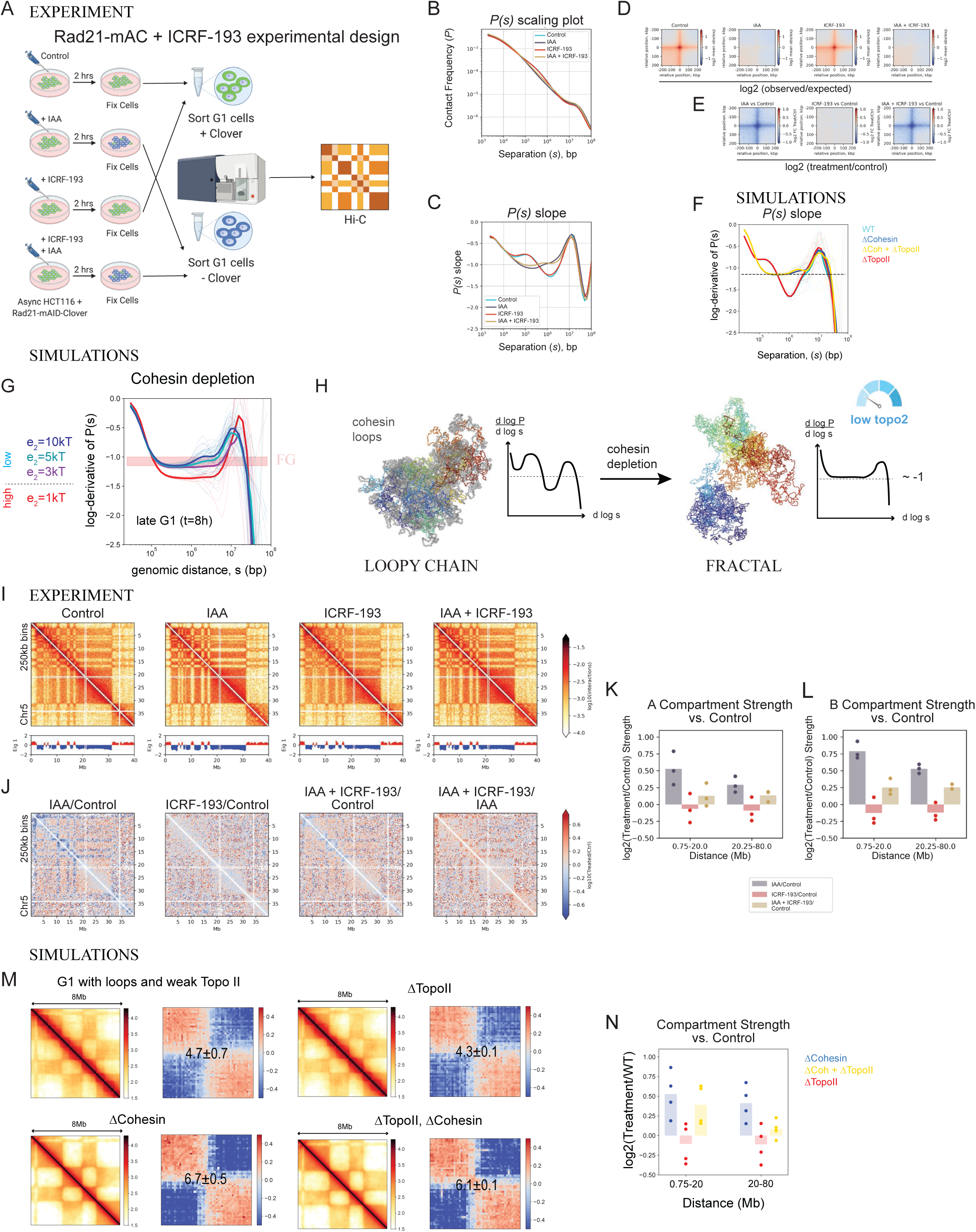
Weak Topo II is required for increased compartment strength due to cohesin degradation. A. Schematic of RAD21 degradation and Topo II inhibition by ICRF-193 in AS HCT116 + RAD21-AID-mClover (HCT116 + RAD21-mAC) cells for Hi-C. Cells were treated for 2 hours with IAA and/or ICRF-193 before fixation for Hi-C, and sorting for G1 DNA content with (-IAA samples) or without (+ IAA samples) mClover. B. P(*s*) scaling plot of G1 sorted HCT116 + RAD21-mAC cells described in A. N = 3. C. First derivative (slope) of P(*s*) scaling plot shown in B. N = 3. D. Aggregate loop pileup (APA) of experiment shown in A at dots called in published high resolution Hi-C data from HCT116 + RAD21-mAC (Untreated) (4DNFIFLDVASC). Average log2(observed/expected). N = 3. E. Log2 fold change of APA for each treatment vs the control F. The *P*(*s*) log-derivatives computed for simulations of four states: unperturbed interphase obtained via the two-stage process with low activity of Topo II (*e*_*tr*_ = 5*kT*), interphase with further inhibited Topo II (ΔTopo II; *e*_*tr*_ = 10*kT*), interphase with further depleted cohesin loops but remained low activity of Topo II (ΔCohesin; *e*_*tr*_ = 5*kT*), interphase with depleted cohesin loops and inhibited Topo II (ΔTopo II, ΔCohesin; *e*_*tr*_ = 10*kT*). G. Simulations of various levels of the strand passage activity during Stage II of the two-stage exit, as modelled by varying the excluded volume barrier. The graph shows the log-derivatives of the contact probability P(s) in interphase after depletion of cohesin loops (late G1). The red strip shows the range of experimental log-derivatives between –1.15 and –1 (panel C). H. A model: Depletion of cohesin loops in the interphase state allows to assess the topological state of chromosomes from the log-derivative of P(s). The slopes around −1 correspond to the fractal organization, which is only consistent with sufficiently weak activity of Topo II in the interphase. I. Experimental Hi-C interaction heatmaps and Eigenvector 1 of HCT116 + RAD21-mAC cells described in A, three replicates combined. J. Hi-C interaction log2 ratio comparing each treatment to the Control or to the IAA treatment, as indicated. Three replicates combined. K. AA compartment strength log2 ratio compared to Control by distance for HCT116 + RAD21-mAC cells, N=3. L. BB compartment strength log2 ratio compared to Control by distance for HCT116 + RAD21-mAC cells, N=3. M. Contact maps and the corresponding compartmental saddle plots from simulations for four states described in F. N. The log2 ratios of the compartment score in perturbed and unperturbed interphase simulations at short and large genomic scales. Each dot represents a ratio computed for a pair of perturbed and unperturbed contact maps, both averaged over 16 replicates. The bars represent the corresponding mean values.

In the course of the second stage of our model the chromosomes become more compartmentalized, reaching the values of compartmental scores observed in DMSO experiments (Figure 6G). Similar to experiments, we also see dissolution of the mitotic band as chromosomes decompact, which is evident both in the contact maps and *P*(*s*) curves (Figure 6G,H). Consistently, the mitotic band dissolves in the distance map of a chromosome (Figure 6J), in sharp contrast with the distance map obtained as a result of expansion without strand passage (Figure 5G, right).

We note that the two-stage expansion out of the hypothetical unknotted mitotic configuration would produce a qualitatively different change of compartmental strength than observed in Topo II inhibition experiments (Figure S5E-F). Indeed, the Topo II activity in Stage I would increase the number of loop-loop catenations from negligible to the level of the two-stage expansion from the knotted mitotic state (Figure S5G), yielding a less compartmentalized state than in the situation of inhibited strand passage without catenations. This result highlights that the unknotted mitotic state is inconsistent with experimental findings on Topo II inhibition.

Taken together, the proposed two-stage mechanism of mitotic exit and decondensation facilitates large-scale chromosome disentanglement, and then maintains this unentangled state allowing the establishment of interphase organization with these hallmarks: the fractal globule, chromosome territoriality, and strong compartmentalization.

### Topo II activity is required for increased compartment strength upon loss of cohesin

To test whether Topo II activity is also required for other forms of chromosome reorganization, we used the previously described HCT116 cell line carrying an auxin inducible degron (AID) and mClover fusion of RAD21, a subunit of the cohesin complex (HCT116 + RAD21-mAC cell line) ^61, 66–68^. In this system, depletion of cohesin results in weaker TADs and CTCF-CTCF loops, and stronger compartmentalization ^61, 68^. Using this system, we measured how chromosome folding is affected by the combined loss of Topo II activity and cohesin by Hi-C. We treated Async. HCT116 + RAD21-mAC cells, which are mainly in the G1 phase of the cell cycle, with 30uM of ICRF-193 to inhibit Topo II or with DMSO, and/or 500uM Auxin (Indole-3-acetic acid, IAA) to degrade RAD21 for two hours (Figure 7A, Figure S7A). Following fixation, the cell populations were sorted for G1 DNA content, and +/-mClover expression (Figure S7B,C).

As previously observed, short-range Hi-C interactions are decreased, while long-range compartment-specific interactions are increased with RAD21 degradation (Figure 7B, C) ^61^. The loss of cohesin-mediated loop extrusion can be observed by a loss of the peak in the derivative of *P(s)* at ∼100 kb, and loss of TADs on 50kb binned heatmaps (Figure 7B, C, Figure S7D-F,H) ^32, 69, 70^. RAD21 depletion + ICRF-193 did not significantly changes the *P(s)* curves compared to RAD21 depletion alone. RAD21 depletion alone reduces loop strength, as previously published ^61^, while ICRF-193 treatment alone has no effect on looping interactions, nor on the effect of cohesin depletion [IAA + ICRF-193] (Figure 7D).

Simulations of cohesin depletion in late interphase, obtained through the two-stage process with some weak Topo II activity in Stage II (see Methods), recapitulate the subtle effects of Topo II inhibition observed in *P*(*s*) curves (Figure 7F); importantly, the model suggests that some Topo II activity in Stage II can still reproduce the fractal scaling of *P*(*s*) observed experimentally (Figure 7G,H), as long as this activity is sufficiently weak.

Compartment strength is increased with RAD21 degradation (Figure 7I-L, FigureS7G) ^61, 69, 70^. However, the compartment strength increase observed with IAA treatment is partially blocked by the addition of ICRF-193, particularly in the B compartment (Figure 7L). Treatment with ICRF-193 alone has only minimal effects on compartment strength, as also observed in the Async HeLa S3 cell line (see Figure S1K,L). We observe similar results in the simulations: while complete inhibition of strand passage alone marginally affects compartmentalization of the interphase chains, in combination with cohesin depletion it yields lower compartmental scores than in the situation of cohesin depletion only (Figure 7M,N). Therefore, weak Topo II activity contributes to the transition in chromosome folding at the compartment scale upon the loss of cohesin.

Importantly, these results reveal two features of the interphase genome: first, loss of cohesin-mediated loops reveal a crumpled chromatin state consistent with an unentangled conformation. Second, the fact that the increase in compartmentalization upon loss of cohesin is partly dependent on Topo II activity suggests Topo II activity during interphase. The two features are consistent with each other as long as the interphase Topo II activity is sufficiently weak, see Figure 7H.

## DISCUSSION

### The mitotic chromosome is internally entangled

Whether intra-chromosomal entanglements occur in mitotic chromosomes has long been an open question in the field, due to an inability to directly measure entanglements in endogenous chromosomes. An unentangled mitotic state has long been the dominant view, as it would naturally expand into an unentangled interphase, which is observed experimentally, assuming strand passage is not active ^34, 35^. However, our results demonstrate that artificial inhibition of Topo II during mitotic exit results in dramatic changes in the subsequent interphase structure. Our polymer simulations show that these experimentally observed changes can be reconciled with an entangled mitotic chromosome that requires Topo II activity to expand into interphase.

Entangled mitotic chromosomes have been previously predicted to form a stiffer structure than unentangled chromosomes, which may be important for ensuring proper chromosome segregation^33^. A recent study using in vitro mitotic chromosome reconstitution in Xenopus egg extracts showed that Topo II activity is required to increase chromosome thickness when condensin is present but results in inter-chromosome entanglements when condensin is depleted^71^. This model of mitotic intra-chromosomal entanglements mediated by Topo II directed by condensin to form mitotic chromosomes is consistent with our entanglement measurements during mitosis^71^.

### Two-stage exit from mitosis

Formation of an unentangled interphase organization from an entangled mitotic chromosome poses a serious challenge: While Topo II activity is required for expansion and compartmentalization, at the same time it prevents the establishment of hallmarks of interphase organization such as chromosome territories and the unentangled fractal globule state. Polymer simulations show that this paradox can be resolved by a two-stage mitotic exit where chromosomes first become unentangled and then are maintained at this state.

In the first stage, decompaction of mitotic chromosomes with mitotic (condensin) loops still present produces a swollen bottlebrush conformation. Loops in this state entropically repel each other, biasing Topo II towards decatenation. As we find in time-calibrated simulations, Topo II needs to be active for ∼10-20 min to largely disentangle condensin loops; this time is close to the condensin residence time ^72, 73^. In the second stage, the disentangled chromosomes expand upon the loss of condensin loops, yet with reduced Topo II activity, allowing to maintain the unentangled state, and thus enabling formation of territorial and fractal globule chromosomes. In accord with our cohesin depletion experiments in G1, polymer simulations further demonstrate that some weak Topo II activity in the second stage is consistent with a largely unentangled interphase organization (Figure 7G).

In a recent synchronized Hi-C study on mitotic exit, the existence of a loop-free, and possibly unentangled state was demonstrated during telophase, when most of the condensins have dissociated and cohesin has not re-associated with chromatin ^40^. This importantly suggests that the disentanglement of mitotic chromosomes takes place during the stage after the metaphase-to-anaphase transition and before the condensins are released from chromosomes. Our current experiments further highlight an important role of Topo II in these early stages of mitotic exit, as its early inhibition results in retention of mitotic-like organization in the following interphase (Figures 1-3). Consistent with these experimental observations, the two-stage model of mitotic exit demonstrates that most of the mitotic entanglements can be removed via decompaction of mitotic chromosomes with condensin loops under high Topo II activity (Stage I). Thus, a swollen bottlebrush state is likely present until telophase onset and linked to decatenation of mitotic loops.

Taken together, our work shows that cells control the entanglement state of the genome during mitotic exit, with important roles for chromosome loops, chromatin decondensation, and regulation of Topo II activity levels.

Our study makes several predictions that future experiments can test. First, the two-stage mechanism suggests the presence of a transient state with associated condensins, expanded mitotic-like state, and high Topo II activity. Detection and characterization of this state, with its unique morphology, chromatin associated condensins and Topo II is a challenge for live-cell microscopy. Second, our models suggest that Topo II inhibition can keep chromosomes in a similar partially expanded mitotic-like state even after condensin loops are gone. We predict how distance maps accessible by high-resolution microscopy^74^ would appear, and how the scaling of the spatial distance with genomic separation would be affected by Topo II inhibition.

### Limitations of the study

First, our study uses chemical inhibition of Topo II activity. ICRF-193 leads to immobilization of Topo II on chromatin, and this may affect chromosome conformation in unknown ways. Alternative methods include the use of degron-based removal of Topo II, but such methods lack the temporal control required for study of chromosome folding dynamics during mitotic exit, when topoisomerases are also required to separate sister chromatids. Second, our proposal that mitotic chromosomes are self-entangled is based on the combined integration of polymer modelling and the analysis of experimental effects of Topo II inhibition (Hi-C, MC-3C data, imaging data). We do not have direct experimental data showing that mitotic loops are catenated. For instance, while MC-3C data show high levels of intermingling of chromatin within condensed prometaphase chromosomes, and a dependence on Topo II activity for unmingling during decondensation, the chromatin interaction data by itself is not providing direct evidence for topological entanglements. Experimental evidence for catenation of mitotic loops will await development of imaging-based methods with sufficient resolution and scale to trace individual loops at nm resolution in 3D. Third, although the proposed two-stage mechanism of mitotic exit seems to be a natural way to disentangle the chromosomes via involvement of regulated topoisomerase activity, it is only tested by polymer modeling.

## ACKNOWLEDGEMENTS

We are grateful to Masato Kanemaki for kindly providing the HCT-116-Rad21-mAID-mClover cell line. We thank several UMass Chan Medical School Facilities for their assistance, including Christina Baer at the Sanderson Center for Optical Experimentation, the Deep Sequencing Core, the PacBio Core Enterprise, the Molecular Biology Core Labs, and the Flow Cytometry Core. We thank Caryn Navarro, Caroline Austin, and members of J.D.’s and L.M.’s groups for helpful discussions and comments on the manuscript. Diagrams included in figures were created with BioRender.com. Experiments in this study used a HeLa cell line. Henrietta Lacks, and the HeLa cell line that was established from her tumor cells without her knowledge or consent in 1951, have made significant contributions to scientific progress and advances in human health. We are grateful to Henrietta Lacks, now deceased, and to her surviving family members for their contributions to biomedical research. We also thank the anonymous patient who contributed the HCT 116 cell line used in this study. We acknowledge support from the National Institutes of Health Common Fund 4D Nucleome Program (DK107980, HG011536), the National Human Genome Research Institute (HG003143) and the National Institute of General Medical Sciences (R01GM114190.). J.D. is an investigator of the Howard Hughes Medical Institute. E.H. was supported by an F32 fellowship from the NIH (F32-CA224689). The work of K.P. is supported by the Russian Science Foundation (Grant No. 21-73-00176). K.P. acknowledges support of the Programme d’investissements d’avenir (LabEx DEEP) at Institut Curie, Paris, France, and the hospitality of LPTMS laboratory (University of Paris-Saclay), where part of this work was done.

## PUBLISHED DATASETS USED

4DNFIFLDVASC, 4DNFIBM9QCFG

## AUTHOR CONTRIBUTIONS

Conceptualization, EH, JD, KP, and LM. Methodology, EH, KP, NF, YLiu. Software, EH, KP, BD, SV. Validation, EH, YLiu. Formal Analysis, EH, KP. Investigation, EH, YLiu, BD, NF, DL, YLing. Data Curation, EH. Writing – Original Draft, EH. Writing – Review and Editing, EH, JD, KP, LM. Visualization, EH, KP. Supervision, JD, LM. Funding acquisition, JD, LM, EH, KP.

## STAR METHODS

### Resource availability

#### Lead contact

Further information and requests for resources and reagents should be directed to and will be fulfilled by the lead contacts, Job Dekker

#### Materials availability

- Plasmids generated in this study have been deposited to Addgene, and will be available upon publication.
- There are restrictions to the availability of the HeLa S3 TOP2A-Venus clone B1 cell line, due to the requirement of a material transfer agreement from ATCC for the use of the original HeLa S3 cell line.

#### Data and code availability

- Hi-C short-read sequencing data have been deposited at GEO and are publicly available as of the date of publication. Accession numbers are listed in the key resources table. MC-3C long-read sequencing data have been deposited at GEO and are publicly available as of the date of publication. Accession numbers are listed in the key resources table. Microscopy data have been deposited at BioStudies and are publicly available as of the date of publication. Accession numbers are listed in the key resources table. Original western blot images have been deposited at Mendeley Data and are publicly available upon the date of publication. The DOI is listed in the key resources table. This paper analyzes existing, publicly available data. These accession numbers for the datasets are listed in the key resources table.
- All original code is available on Zenodo, and will be publicly available as of the date of publication. Zenodo URLs are listed in the key resources table.
- Any additional information required to reanalyze the data reported in this paper is available from the lead contact upon request.

### EXPERIMENTAL MODEL AND STUDY PARTICIPANT DETAILS

- HeLa S3 CCL-2.2 cells (ATCC, CCL-2.2) were cultured in DMEM, high glucose, GlutaMAX^TM^ supplement with pyruvate (Gibco, 10569010) supplemented with 10% fetal bovine serum (Gibco, 16000044) and 1% penicillin-streptomycin

(Gibco, 15140) at 37 degrees C in 5% CO_2_.

- HeLa S3 CCL-2.2 cells (ATCC, CCL-2.2) with TOP2A-Venus were cultured in DMEM, high glucose, GlutaMAX^TM^ supplement with pyruvate (Gibco, 10569010) supplemented with 10% fetal bovine serum (Gibco, 16000044) and 1% penicillin-streptomycin (Gibco, 15140) at 37 degrees C in 5% CO_2_.
- HCT116 + RAD21-mAC cells ^67^ were cultured in McCoy’s 5A (Modified) Medium, GlutaMAX^TM^ supplement (Gibco, 36600021) supplemented with 10% fetal bovine serum (Gibco, 16000044) and 1% penicillin-streptomycin (Gibco, 15140) at 37 degrees C in 5% CO_2_.

### METHOD DETAILS

#### Creation of a stable HeLa S3 TOP2A-Venus cell line

The cell line was constructed as described for generation of a stable HeLa S3 NCAPH-dTomato cell line in Abramo et al, 2019^40^. Briefly, pSpCas9(BB)-2A-Puro (BX459) v.2.0 (a gift from F. Zhang (Addgene plasmid, 62988; RRID, Addgene_62988)) was used to construct CRISPR/Cas vectors according to the protocol of Ran et al ^75^. gRNA sequences are provided in Key Resources Table. To construct the donor plasmid for C-terminal integration of Venus, pUC19 (Thermofisher, Cat SD0061) was used as the backbone and was constructed using synthesized DNA and homology arms generated by PCR (primers provided in Key Resources Table). Genomic DNA from HeLa S3 cells was used as template, and was amplified using Q5 High-Fidelity DNA Polymerase (New England Biolabs Cat M0491S) to generate 800 bp *TOP2A* homology arms. A gBlock containing 5X Glycine linker, Venus, T2A and Hygromycin resistance was synthesized by Integrated DNA Technologies (IDT) (sequence is provided in Key Resources Table). Homology arms and gBlocks were cloned into pUC19 by Gibson assembly using NEBuilder HiFi DNA Assembly Master Mix (NEB E2611S). To generate stable cell lines, 5×10^6 cells were electroporated with gRNA plasmids and homology arm donor plasmid. Then, 24h after electroporation, 1ug/ml puromycin selection was added, and, 2d later, 150 ug/mL hygromycin B Gold (Invivogen # ANTHG1) was added for TOP2A-Venus selection. After 4 d, colonies were picked for further selection in a 96 well plate.

#### Cell Culture

HeLa S3 CCL-2.2 cells (ATCC, CCL-2.2) with or without TOP2A-Venus were cultured in DMEM, high glucose, GlutaMAX^TM^ supplement with pyruvate (Gibco, 10569010) supplemented with 10% fetal bovine serum (Gibco, 16000044) and 1% penicillin-streptomycin (Gibco, 15140) at 37 degrees C in 5% CO_2_. HCT116 + RAD21-mAC cells^67^ were cultured in McCoy’s 5A (Modified) Medium, GlutaMAX^TM^ supplement (Gibco, 36600021) supplemented with 10% fetal bovine serum (Gibco, 16000044) and 1% penicillin-streptomycin (Gibco, 15140) at 37 degrees C in 5% CO_2_. Topo II inhibition was performed by adding ICRF-193 (Sigma-Aldrich, I4659-1MG) dissolved in DMSO (4mg/ml stock solution) and used at 30uM final concentration in cell culture media, or Merbarone (Santa Cruz Biotechnologies, sc-500526) dissolved in DMSO, at a 200uM final concentration. A 30uM dose of ICRF-193 is sufficient to inhibit Topo II, as this dose blocks chromosome segregation and delays mitotic exit when added at t0 (nocodazole wash-out), as measured by cyclin B degradation in HeLa S3 cells and analysis of DNA content by flow cytometry (Figure S1A-G)^42–44^. In addition, it has been shown that a five-hour treatment with 35uM ICRF-193 results in an overall loss of DNA supercoiling in RPE1 cells, and blocks changes in supercoiling due to transcription inhibition^44^. ICRF-193 potency was tested for each batch by adding 30uM ICRF-193 or an equal volume of DMSO at *t* = 0 to a sample of washed prometaphase arrested HeLa S3 cells re-cultured in standard media, and collecting both the floating and adherent cells for cell cycle analysis by PI staining and flow cytometry after four hours. Experiments were only continued if the ICRF-193 at *t* = 0 treatment arrested cells with G2/M DNA content, by inhibiting decatenation of sister chromatids, but the DMSO control allowed the majority of cells to enter G1. Chromosome copy number of G1 cells is not significantly changed by ICRF-193 treatment (data not shown). Transcription inhibition was performed by adding 25uM Triptolide (TRP) (Millipore, 645900-5MG, stock was 55mM in DMSO) and 200uM 5,6-Dichlorobenzimidazole 1-β-D-ribofuranoside (DRB) (Sigma, D1916-50MG, stock was 70mM in DMSO) and incubating for the indicated times at 37 degrees C in 5% CO_2_. Auxin induced degradation of RAD21-AID-mClover in the HCT116 + Rad21-mAC cells was performed by adding 500uM 3-Indoleacetic acid (IAA, auxin) (Sigma, 45533-250MG) dissolved in 100% Ethanol, and incubating for the indicated times at 37 degrees C in 5% CO_2_.

#### Mitotic Synchronization and Release Time Course

Prometaphase synchronization and release of HeLa S3 CCL-2.2 cells for Hi-C, MC-3C, and IF was performed as previously described^40, 41^. This protocol routinely produces cell cultures where >90% of cells display high levels of p-H3 (data not shown). For all experiments except DAPI and TOP2A-Venus contrast measurements by microscopy, a single thymidine block was used to first arrest cells in S phase: cells were plated at 4×10^6 cells per 15 cm plate in medium containing 2mM Thymidine (Sigma, T1895), and were grown for 24 hours. Next, cells were washed with 1x DPBS (Gibco, 14190144) and standard medium was added to the plates for 3 hours to allow release from S phase and recovery of cells. For all synchronization experiments, 100ng/ml nocodazole (Sigma, M1404) was added for 12 hours to synchronize cells in prometaphase, by depolymerizing the spindle microtubules. Floating mitotic cells were collected by mitotic shake-off and washed in 1x DPBS (Gibco, 14190144). Mitotic samples were collected directly from washed prometaphase cells and immediately prepared for downstream purposes. The remaining cells were re-cultured in pre-warmed standard medium for synchronous release into G1 and were collected at indicated timepoints post-mitotic release, depending on the experiment. For the *t* = 2 hrs timepoint, both the floating and adherent cells were collected. For timepoints from *t* = 4hrs to *t* = 9hrs, only adherent cells were collected, unless otherwise indicated. For Hi-C, MC-3C, and IF, 30uM ICRF-193 (Sigma-Aldrich, I4659-1MG) in DMSO, or an equal volume of DMSO as a vehicle control, was added at *t* = 2 hours.

#### Flow Cytometry

For cell cycle analysis, approximately 0.25-1×10^6 cells at each timepoint were collected in 15 ml conical tubes. For samples where both floating and adherent cells were collected, the media was first transferred to the conical, then the plate was washed 1x with 1x DPBS (Gibco, 14190144), and then the adherent cells were released using StemPro Accutase (ThermoFisher, A11105-01) or TrypLE Express (ThermoFisher, 12605036). For adherent only samples, the media was instead discarded. Released cells were transferred to the 15 ml conical, and were then washed 1x with 1x DPBS (Gibco, 14190144), and resuspended in 100ul of 1x DPBS. Cells were fixed with 1 ml of cold 100% Ethanol added drop-wise while vortexing on the lowest setting, to reduce clumping. Samples were then incubated at −20 degrees C for at least 30 minutes, or stored at this temperature until staining was continued. For propidium iodide (PI) staining, fixed cells were washed in 1x PBS (Gibco, 70013-073), and then resuspended in 1x PBS containing 0.1% NP-40 (MP Biomedicals, 0219859680), 0.05mg/ml RNAse A (Roche, 10109169001) and 5 or 50ug/ml propidium iodide (Thermo, P1304MP). The samples were incubated at room temperature (20 degrees C) for 30 minutes, and were then analyzed using a BD LSR II, MACSQuant VYB, or BD Fortessa flow cytometer. For the LSR II, PI was excited by either a yellow laser (561nm), with the PI fluorescence collected with either a 600LP dichroic mirror and a 610/20 filter or a 570LP dichroic mirror and a 580/20 filter, or by a blue laser (488nm) with a 685LP dichroic mirror and a 695/40 filter. For the MACSQuant VYB, PI was excited by either a yellow/green laser (561nm), and the PI fluorescence was collected with either a 586/15 filter or a 615/20 filter, or was excited by a blue laser (488nm) and PI fluorescence was collected with a 614/50 filter. For the BD Fortessa, PI was excited by 561 nm laser, and the PI fluorescence was collected with a 586/15 filter. No compensation was required. For analysis of RAD21-mAC degradation, cells were fixed and PI stained as for Hi-C with 1% Formaldehyde (Fisher, BP531-25), to prevent loss of free RAD21-mAC due to ethanol induced cell permeabilization (see below for full method). Cells were run on the MACSQaunt VYB, PI was excited by a yellow/green laser (561nm), and the PI fluorescence was collected with either a 586/15 filter or a 615/20 filter, and Clover was excited by the blue laser (488nm), with Clover fluorescence collected with a 525/50 filter. No compensation was required.

For analysis of apoptotic cells using Annexin-5 staining, the FITC AnnexinV/Dead Cell Apoptosis Kit (Invitrogen, V13242) was used. Cells were run on the MACSQuant VYB: PI was excited by a yellow/green laser (561nm), and the PI fluorescence was collected with either a 586/15 filter or a 615/20 filter, and FITC was excited by the blue laser (488nm), with FITC fluorescence collected with a 525/50 filter. No compensation was required.

#### Hi-C

Hi-C was performed using the Hi-C 2.0 protocol^39^. For unsorted samples, approximately 5×10^6 cells were fixed in 1% formaldehyde (Fisher, BP531-25) diluted in serum-free media, as previously described. For G1 sorted samples, 10-20×10^6 adherent cells were first removed from the plate using StemPro Accutase (ThermoFisher, A11105-01) or TrypLE Express (ThermoFisher, 12605036), were washed 1x in HBSS (Thermofisher, 14025134), and then were fixed in 1% formaldehyde (Fisher, BP531-25) in HBSS (Thermofisher, 14025134), to reduce clumping during FACS. Formaldehyde fixation was quenched with 0.125M Glycine for 5 minutes at room temp, and 15 minutes on ice. Cells were washed after fixation with DBPS, and the cell pellets were flash frozen in liquid N_2_ and stored at −80 degrees C. For G1 sorting, flash frozen cross-linked cell pellets were thawed on ice and were partially permeabilized using 0.1% Saponin in 1x PBS. (Sigma-Aldrich, 47036-50G-F). Cells were then treated with 2mM MgCl2, 100ug/ml RNAse A (conc, Roche, 10109169001) and 50ug/ml conc propidium iodide (Thermo, P1304MP) for 30 minutes at room temperature with gentle mixing. Cells were spun down and resuspended in 1x PBS (Gibco, 70013-073) + 1-3% BSA (Sigma Aldrich, A7906-50G) + 50ul/ml propidium iodide, and were sorted for G1 DNA content for the HeLa S3 experiments, or for G1 DNA content plus or minus mClover (GFP channel) signal for the HCT116 + Rad21-mAC experiments, into PBS + 1-3% BSA using a BD FACS Melody with the following channels: FSC, SSC, PI: 561nm laser, 605LP dichroic mirror, 613/18 filter, Clover (FITC channel): 488nm laser, 507LP dichroic mirror, 427/32 filter or a BD FACS Aria IIu with the following channels: FSC, SSC, PI: 561nm laser, 600LP dichroic mirror, 610/20 filter or 488nm laser, 655LP dichroic mirror, 695/40 filter, Clover (FITC channel): 488nm laser, 502 or 505LP dichroic mirror, 530/30 or 525/50 filter. After sorting, cells were pelleted by centrifugation, flash frozen in liquid N_2_, and stored at −80 degrees C. Flash-frozen cross-linked cells either with or without sorting were thawed on ice for Hi-C, and were then lysed and digested with DpnII (NEB, R0543M) at 37 degrees C overnight, following the Hi-C 2.0 protocol^39^. The overhanging DNA ends were filled in using biotin-14-dATP (LifeTech, 19524016) at 23 degrees C for four hours, and ligated with T4 DNA ligase (Life Technologies, 15224090) at 16 degrees C for four hours. Chromatin was then treated with proteinase K (ThermoFisher, 25530031) at 65 degrees C overnight to remove all cross-linked proteins. Ligation products were purified by phenol:chloroform extraction with ethanol precipitation, fragmented by sonication, and size-selected using SPRI beads to retain fragments of 100-350bp. Next, we performed end repair and then selectively purified biotin-tagged DNA using streptavidin coated beads (DYNAL™ MyOne™ Dynabeads™ Streptavidin C1, Invitrogen 65001). A-tailing and Illumina TruSeq adapter ligation (Illumina, 20015964) were performed on the bead-bound ligation products, and samples were amplified using the TruSeq Nano DNA Sample Prep kit (Illumina, 20015964). PCR primers were removed using SPRI beads (1.1x ratio) before sequencing the final Hi-C libraries using PE50 bases on an Illumina HiSeq 4000 or NextSeq 2000.

#### MC-3C

MC-3C was performed as previously described, with some changes in the crosslinking and ligation protocols^27^. Cells were collected and fixed following the Hi-C 2.0/MC-3C protocol ^27, 39^, for adherent cells dissociated for G1 sorting as described above (G1 HeLa S3 samples, and adherent population of *t* = 2 hrs HeLa S3 samples), or from suspension cells (*t* = 0 prometaphase and floating population of *t* = 2 HeLa S3 samples), with the following changes: After quenching of formaldehyde crosslinking, cells were washed 2x with DPBS (Gibco, 14190144), and were additionally fixed with 3mM Disuccinimidyl Glutarate (DSG) (ThermoFisher, 20593) diluted in DPBS from a 300mM stock in made with DMSO and incubated at room temperature for 45 minutes. DSG crosslinking was quenched by addition of 0.75M Tris pH 7.5^76–78^. Cells were pelleted (1000xg) and then washed 2x after fixation with DBPS. Cell pellets were flash frozen in liquid N_2_ and stored at −80 degrees C until either G1 sorting or starting MC-3C. For MC-3C, flash-frozen G1 sorted or unsorted cross-linked samples were thawed on ice, and followed the Hi-C 2.0/MC-3C protocol through DpnII digestion ^27, 39^. Importantly, throughout the remainder of the protocol care was taken to reduce pipetting of samples to avoid shearing and damaging DNA at all steps, and mixing was performed by invert mixing and light vortexing only. After DpnII digestion and inactivation, the overhanging DNA ends were ligated using 50uL T4 DNA ligase (Life Technologies, 15224090) at 16 degrees C for 4 hours in a ligation buffer containing 1x Invitrogen Ligation Buffer, 0.1% Triton-X-100 (Sigma, 93443-500mL), and 0.01mg/mL BSA (Sigma Aldrich, A7906-50G). Ligated chromatin was then treated with 2x 50ug Proteinase K (ThermoFisher, 25530031) at 65 degrees C overnight to remove all cross-linked proteins, as in the published Hi-C 2.0/MC-3C protocols ^27, 39^. Ligation products were purified by phenol:chloroform extraction with ethanol precipitation, as in Hi-C 2.0/MC-3C ^27, 39^, with a final elution in 20ul dH_2_O.

#### PacBio Library Preparation

Barcoded PacBio SMRTbell libraries for MC-3C replicate 1 of the mitotic release experiment were constructed by the UMass Chan Medical School PacBio Core Enterprise. All other MC-3C PacBio libraries were constructed using the following protocol. Barcoded PacBio SMRTBell adapters from the PacBio 8A barcoded adapter kit (Barcoded Overhang Adapter Kit - 8A, PacBio PN: 101-628-400) were added to ligation products following an adapted version of the PacBio Microbial Multiplexing PacBio protocol (Part Number 101-696-100 Version 07 (July 2020)) using the PacBio SMRTbell Express Template Prep Kit 2.0 (PacBio, PN: 100-938-900) as follows: the ligated DNA was quantified using a Qubit fluorometer and Qubit HS DNA reagents, as recommended by the manufacturer (ThermoFisher, Q32851). DNA quantification was adjusted to 33ng/ul with 1x PacBio Elution Buffer (PacBio, PN: 101-708-100). Then 14.6ul of each ligated DNA sample was aliquoted into a 1.5ml DNA lo-bind tube (Eppendorf, Z666548). All reagents and master mixes were kept on ice throughout this protocol. To remove single-strand overhangs, DNA prep additive was first diluted 1:5 in enzyme dilution buffer. Next, the DNA prep master mix was added. For an 8-plex reaction, volumes including 25% overage for this mix were: 23.3ul DNA prep buffer, 3.3ul NAD, 3.3ul Diluted DNA Prep Additive, 3.3ul DNA Prep Enzyme. For each sample, 3.3ul of the DNA prep master mix were added to 14.6ul ligated DNA. Tubes were mixed by finger-tapping and briefly spun down, then incubated at 37 degrees C for 15 minutes. Reactions were then returned to 4 degrees C. Next, the DNA Damage Repair master mix was prepared. For an 8-plex reaction, 6.7ul of DNA Damage Repair Mix v2 + 3.3ul Enzyme Dilution Buffer were added. Then 1.0ul of DNA Damage MM was added to each 17.9ul single strand digested sample. Tubes were mixed by finger-tapping and briefly spun down, then incubated at 37 degrees C for 30 minutes and returned to ice. End-Repair/A-tailing was performed in one step by adding 1ul of End Prep Mix to each 18.9ul damage-repaired sample. Tubes were finger-tapped to mix, briefly spun down, and incubated at 20 degrees C for 10 minutes, followed by 65 degrees C for 30 minutes, then returned to 4 degrees C. For ligation of barcoded overhang SMRTBell adapters, 2.0ul of adapters were added to each A-tailed sample first before adding ligase. Tubes were finger-tapped to mix and briefly spun down, then were placed on ice while the Ligation Master Mix was prepared. For an 8-plex reaction, the Ligation Master Mix contained 88ul Ligation Mix, 2.9ul Ligation Additive, and 2.9ul Ligation Enhancer. 10.7ul of ligation master mix was added to each A-tailed/Barcoded Adapter mix, followed by finger-tapping to mix, and a brief spin. Samples were incubated at 20 degrees C for 60 minutes, and then the ligase was heat killed by incubating at 65 degrees C for 10 minutes. SMRTBell libraries were then purified using 0.45X AMPure PB Beads, either 1 or 2 times. For 0.45X AMPure PB purification, AMPure PB beads were first brought to room temperature. Next, the volume of each library was adjusted to 100ul with PacBio Elution Buffer, and 45ul of the AMPure PB beads (0.45X volume) was added. Bead/DNA solution was mixed by gently tapping the tube, followed by a quick spin down to collect the beads. Mix was incubated at room temp for 5 minutes, quickly spun down, and placed on a magnetic bead rack for 5 minutes to collect the beads to the side of the tube. Once supernatant was cleared, it was slowly pipetted off and discarded. Library DNA was now bound to beads. Beads were washed 2x with 80% ethanol without removing tubes from magnet by slowly pipetting 1ml into each tube, incubating for 30 seconds, and then slowly pipetting out the ethanol. Residual ethanol was removed by quickly spinning the tubes and returning to the magnetic tube rack. Any residual 80% ethanol droplets were pipetted off of the beads. For the first AMPure purification, DNA was eluted in 100ul 1x PacBio Elution buffer, while for the second purification DNA was eluted in 20ul Elution Buffer. To elute DNA, elution buffer was added to the tube and tube was mixed by finger-tapping to resuspend beads. Mix was incubated at 37 degrees for 15 minutes, then quick spun and placed on the magnetic bead rack until supernatant was clear. Supernatant containing DNA library was transferred to a new 1.5ml DNA Lo-Bind tube using a wide-bore lo-bind tip for the 100ul eluate, and a 20ul regular bore tip for the 20ul eluate, and beads were discarded. DNA quantity was measured using a Qubit fluorometer and the Qubit HS DNA kit. Libraries were analyzed by fragment analyzer to determine average size, and were sequenced on a PacBio Sequel II, using v8.0.0 (replicates 1 and 2) or v10.1.0 (replicate 3) instrument control software.

#### Immunofluorescence Microscopy Antibody Staining

IF samples were prepared using standard methods. HeLa S3 or HeLa S3 + TOP2A-Venus cell lines were either grown or spun onto 22×22mm No 1.5 coverslips in 6-well plates (VWR Cat No 48366-277). (cell cycle analysis experiments, euchromatin and heterochromatin localization experiments) or were spun onto 25×75×1.0mm Superfrost Plus Precleaned Microscope Slides with 13mm Single Rings (Fisher Cat No. 22-037-241) using a Cytospin 4 centrifuge (DNA and Topo IIα-Venus contrast experiments) (800rpm, 5min). For experiments with cells on coverslips, for adherent cells, the media was removed, cells were washed with DPBS (Gibco, 14190144), and were fixed in 4% Paraformaldehyde (Fisher, 15710) in 1x PBS (Gibco, 70013-073) for 10 minutes at room temp onto 22×22mm coverslips in a 6-well plate, and were then washed 3x with 1x PBS before proceeding to staining. For suspension cells, an equal volume of the cell suspension in media and 8% paraformaldehyde in 1x PBS was mixed together and added to a 6-well plate containing a 22×22mm coverslip. The plate was then centrifuged for 15 min at 800xg, during which time the cells were fixed and became adhered to the coverslip, and were then washed 3x with 1x PBS before proceeding to staining. For experiments using the Cytospin 4, for adherent cells the media was first collected, and cells were washed 1x with DPBS before cells were disassociated with TrypLE Express (ThermoFisher, 12605036), and combined with the collected media. For suspension cells, the cell suspension was collected directly. Cell suspensions were spun to pellet cells, and were washed 1x with 1x PBS, and then fixed for 10 minutes in 4% Paraformaldehyde at room temp. After fixation, cells were washed 1x with PBS, then resuspended in 1ml PBS and stored at 4 degrees C in dark. 0.5-1.5×10^5 fixed cells were spun onto slides using the Cytospin 4, using Shandon™ EZ Single Cytofunnel™ with White Filter Cards and Caps (Thermo Fisher, A78710020). After removal of the Cytofunnel, cells were allowed to dry for a few minutes, before proceeding to staining. To stain cells, slides or coverslips were first incubated in block buffer (3% BSA (Sigma Aldrich, A7906-50G), 0.1% Triton-X-100 (Sigma, 93443-500mL), 1x PBS) in a humidified chamber for 30 minutes to 2 hours at room temp, or overnight at 4 degrees C. To prevent evaporation, droplets of buffer on cells were covered with squares of parafilm during incubation. After blocking, primary antibody in block buffer was added to each slide or coverslip, which was incubated for 2 hours at room temp or overnight at 4 degrees C. Slides or coverslips were washed 3x 5 min in 1x PBS + 0.1% Triton-X-100 after primary antibody incubation in a, either in Coplin jars (slides), or in 6 well plate (cover slips). Secondary antibody in block buffer was added next and incubated for 30 minutes at room temp, in a dark humidified chamber. Slides or coverslips were then washed 3×5min in 1x PBS + 0.1% Triton-X-100, and then 2x in PBS without detergent. For DAPI stained coverslips or slides, this was followed either by a 10min incubation at room temp with 3uM DAPI in PBS (Invitrogen, D1306), or by using ProLong Diamond mounting media containing DAPI (Invitrogen, P36962). Before mounting, slides or coverslips were washed 5min in distilled water and allowed to air dry, they were then mounted using ProLong Diamond mounting media either with DAPI (Invitrogen, P36962) or without DAPI (Invitrogen, P36961), and allowed to cure for at least 24 hours in the dark before imaging. Primary antibodies used: rabbit anti-Lamin A (Abcam ab26300, 1:1000 dilution), rabbit anti-H3K9me3 (Abcam ab8898, 1:500 dilution), mouse anti-H3K27ac (Active Motif 39085, 1:500 dilution), mouse anti-Alpha Tubulin (Sigma T6199, 1:5000 dilution). Secondary antibodies used: goat anti-mouse IgG H+L Alexa Fluor 568 (1:1,000, Abcam ab175473), goat anti-rabbit IgG H&L Alexa Fluor 488 (1:1,000, Abcam ab15007)

#### Widefield Fluorescence Microscopy

For widefield fluorescent image acquisition, we used a Nikon Eclipse Ti microscope. Imaging was performed using an Apo TIRF, N.A. 1.49, 60x oil immersion objective (Nikon), and a Zyla sCMOS camera (Andor). Images were acquired using Nikon Elements software (Version 4.4). Data are available in the BioStudies database (https://www.ebi.ac.uk/biostudies/) ^79^.

#### Confocal Fluorescence Microscopy

For image acquisition, we used a Nikon A1 point-scanning confocal microscope with 405 nm, 488 nm and 561 nm lasers. Imaging was performed using an Apo TIRF, N.A. 1.49, 60x oil immersion objective (Nikon) with GaAsP detectors (488 and 561 lasers) or a high sensitivity MultiAlkali PMT (405 laser) at 0.1um/pixel resolution, resulting in 512/512 pixel images, in Galvano imaging mode. Z-steps were 1um for images taken as Z-series. Images were acquired using Nikon Elements software (Version 4.4). Data are available in the BioStudies database (https://www.ebi.ac.uk/biostudies/) ^79^.

#### Western Blots

Protein immunoblots were performed using standard methods. Protein lysates were made by harvesting the same number of cells (generally 0.5×10^6 each) for each sample in an experiment and lysing in 2x Laemmli Buffer (.07M Tris pH6.8, 10% sucrose, 3% SDS, 50mM DTT) by boiling for 10 minutes. Western blots were performed by running protein lysates on 4-12% NuPage Bis-Tris gels using 1x MES running buffer for 45 minutes at 175V using an Invitrogen XCell Sure-Lock minigel and blotting system. Gels were transferred to 0.2um nitrocellulose membrane in Pierce™ 10X Western Blot Transfer Buffer, Methanol-free (Thermo Fisher Scientific, 35040), by running for 1 hour at 30V at 4 degrees C. The membranes were blocked with 4% milk in PBS-T (1x PBS and 0.1% Tween-20) for 30 minutes at room temperature. The membranes were then incubated with the specified antibodies diluted in 4% milk/PBS-T either for 2 hours at room temp or overnight at 4 degrees C, and washed three times with PBS-T for 10 minutes at room temperature. Membranes were then incubated with secondary antibodies (anti-rabbit IgG or anti-mouse IgG HRP linked) diluted 1:4000 in 4% milk/PBS-T for 30 minutes to 1 hr at room temperature, then washed three times with PBS-T for 10 minutes at room temperature. The membranes were then developed and imaged using SuperSignal West Dura Extended Duration Substrate (Thermo, 34076) and a Bio-Rad ChemiDoc. Primary antibodies used: mouse anti-Topoisomerase IIα (1:500, Santa Cruz sc-166934), rabbit anti-DNA Topoisomerase IIα and DNA Topoisomerase IIβ (1:10,000, Abcam ab109524), rabbit anti-GFP (1:5,000, abcam ab290), mouse anti-β-actin (1:2,000, Cell Signaling 8H10D10), mouse anti-cyclin B1 (1:500, Cell Signaling 4135), rabbit anti GAPDH (1:1000, Cell Signaling 14C10). Secondary antibodies used: goat anti-rabbit IgG-HRP (1:4,000, Cell Signaling 7074), goat anti-mouse IgG-HRP (1:4,000, Cell Signaling 7076).

#### Details of polymer modeling

##### Simulations setup

Polymer simulations were performed using polychrom (available at https://github.com/open2c/polychrom), a wrapper around the open source GPU-assisted molecular dynamics package OpenMM ^80^. We simulated a bead-spring chain of length *N* = 22500 in the periodic boundary conditions (PBC), which were effectively emulating interactions of different chromosomes with each other. Each bead represented 4kb of chromatin (*N* = 90 Mb in total). With chromatin linear density 80bp/nm the contour length of the 4kb chromatin segment was equal to *l* = 50 nm. The interphase volume density of such a chain was 35%, which corresponds to the recent estimate from the electron microscopy tomographic analysis (Ou et al, Science 2018).

Rigidity of the bead-spring chain was adjusted to reproduce the value of the Kuhn length *l_k_* = 100 nm ^81^. Since it corresponded to 2 beads of the contour length *l* the persistence length was *l_p_* = 4 kb in the framework of the worm-like chain model ^82^. To further estimate the spatial size σ of one 4kb bead, we computed the corresponding gyration radius for a straight segment of length *l*; then the size of the bead σ is twice the gyration radius, i.e. 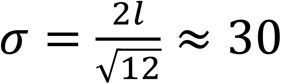 nm.

The excluded volume potential *U_e_*_*v*_ between a pair of beads *i* and *j* was introduced through the auxiliary Weeks-Chandler-Anderson (WCA) potential **U*_*WC*A_ (r_i_, r_j_)* = *U_WCA_*(*r_ij_* = |*r_i_*-*r_j_*|) 1kt which is a lifted repulsive branch of the Lennard-Jones potential

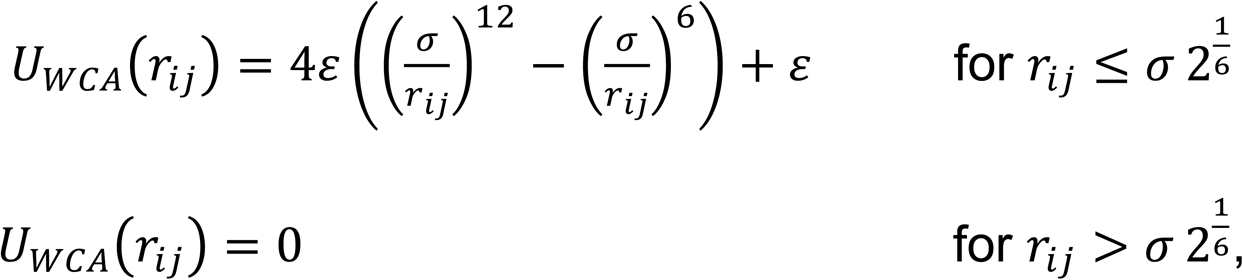

where *σ* is the characteristic length scale of excluded volume interactions, corresponding to the spatial size of one bead; at **r*_ij_* = σ the potential **U*_*WC*A_*(*σ*) = ε = 1*kT*. In order to avoid singularities of the excluded volume potential (infinite forces), the WCA potential was further smoothly truncated at the prescribed truncation value *ε_*tr*_* as follows

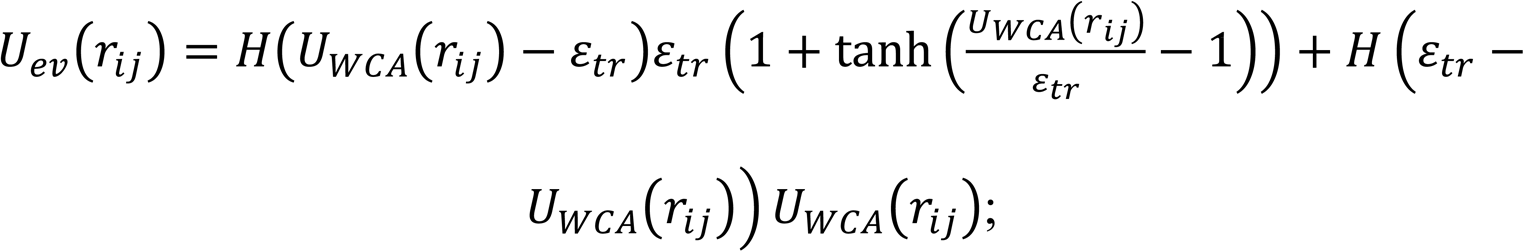

*H*(*x*) is the Heaviside step function.

The energy of harmonic bonds (springs) between the beads reads as follows

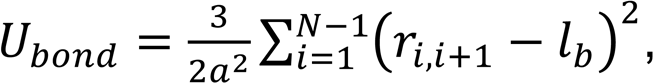

where *a* is the standard deviation of the bead-to-bead spatial distance *r_i,i_*_+1_ = |*r*_i_ − *r*_i+1_| from the equilibrium length of the bond *l*_*b*_. We preset *a* ≈ 0.06*σ*. In order to repress occasional crossing of two closely located polymer bonds in space, we chose the value of the equilibrium bond length *l*_*b*_ = 0.8*σ* and further switched off the excluded volume for neighboring beads. As we previously found ^31^, such a 20% reduction of the bond length compared to the excluded volume scale together with a high excluded volume barrier between non-neighboring beads ensures conservation of the global chain topology in simulations. To check for that, we computed the matrix of pairwise catenations (Gaussian linking number) between condensin loops and made sure that both in mitotic and bottle-brush phases (Stage I, see below) the loops remained mutually non-catenated. Notably, for *l*_*b*_ = *σ* some occasional strand passage occurred, despite of the high excluded volume barrier. Similarly, a reduction of the equilibrium bond length was previously implemented in the Kremer-Grest model ^83^, however, with the FENE bond potential, which logarithmically diverges at some critical distance. In contrast, in this work we used soft harmonic bonds, which is a more convenient choice for modeling of dynamic extrusion of loops. The resulting persistence length *l*_*p*_ in the model was estimated by fitting the end-to-end squared distance *R*^2^(*s*) of a segment of length *s* to the theoretical expression for the worm-like chain model ^82^. The polymer framework with conserved topology allows for a tunable activity of Topo II in simulations. Completely prohibited strand passage was modeled with a high excluded volume barrier, which was parametrized by the truncation threshold ε_*tr*_ = 10*kT*. Weak activity of Topo II was modeled by smaller values of the barrier, such as ε_*tr*_ = 5*kT*, 2*kT*. High activity of Topo II corresponded to the barrier of ε_*tr*_ = 1*kT*.

##### Initial mitotic state

For each of the replicates the simulations of mitotic exit were initiated with a chromosome folded into a sequence of 225 random loops, exponentially distributed in size with the mean *λ* = 200*kb* (except for the last one, for which the size was chosen such that the lengths of all the loops sum up to *N*). The size of mitotic loops in our simulations corresponded to the size of condensin II loops, found previously^46^; also, it can be independently estimated ^31^ in our data from the peak position on the log-derivative of the average contact probability P(s) (Fig. 1G).

A chromosome with loops was further constrained within a cylindrical volume with the longitudinal length *L* = 50*σ* and the radius *R* = 13*σ*, such that the volume density of the mitotic chromosome was equal to ∼0.85. Similarly to Gibcus et al^46^, in order to obtain spiralization of the backbone, we equidistantly tethered each 20^th^ loop at the axis of the cylinder. Also, the ends of the chain were tethered to the caps of the cylinder at the axis.

In order to prepare an initial state with mutually non-catenated mitotic loops we performed slow extrusion of the loops with sizes sampled from the exponential distribution, under inhibited strand passage conditions (high excluded volume barrier, ε_*tr*_ = 10*kT*), followed by the equilibration of the mitotic structure. As the matrix of linking numbers between the loops shows (Figure 5A), this indeed resulted in mutually non-catenated loops on the chain. An initial state with catenated loops was then obtained from the non-catenated mitotic state by decreasing the barrier of excluded volume to ε_*tr*_ = 1*kT* (high activity of Topo II) and equilibrating the chain again. Eventually, the mean number of catenations per one loop in the knotted mitotic state was around ∼0.7 (see Figure 5A), which corresponded to the steady state at conditions of fluctuating topology. The mean genomic distance between the catenated loops in this state was equal to ∼9Mb (∼22 loops), which is close to the size of the mitotic layer.

##### Compartments and TADs

In the course of the mitotic exit we added compartmental interactions, giving rise to compartments, and active loop extrusion, yielding topologically-associated domains on the simulated Hi-C maps. In our main model, the two features were added to the simulation at the same time, namely when mitotic loops were removed after T=10min of the Stage I (in both, inhibited strand passage and wild type simulations).

In order to properly introduce compartmental interactions, we used the experimental annotation into the compartments. For that, we computed the first eigenvector of the centralized observed over expected Hi-C map (Late G1 DMSO combined R1 + R2 sample, see Hi-C Compartment Analysis methods section for details) for chromosome 14 in the 250kb resolution. The first 20Mb segment of the chromosome consists of NaN values and, thus, it was eliminated from the analysis. The remaining part of the eigenvector was used to assign the compartmental types: positive components corresponded to the type A, negative and zero components corresponded to the type B (each component of the eigenvector generated a batch of 63 4kb-monomers of the same type). To comply with the smoothed and truncated excluded volume potential, used for the interactions of the beads of different type, we introduced the following attractive potential between the beads of the same type

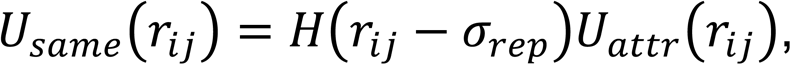

where 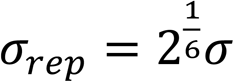 is the scale, above which the repulsion between the beads is absent; attraction of the same-type beads is turned on according to the potential

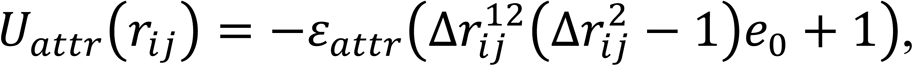

where *e*_0_ = 823543.0/46656.0 is some constant and Δ*r*_i/_ is the scaled distance defined as follows

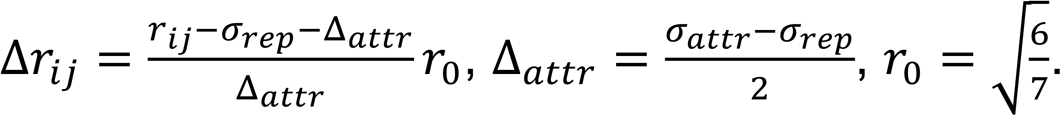

The special choice of the constants *e*_C_ and *r*_C_ ensured that the potential smoothly went to zero at *σ*_*r*e*p*_ and *σ*_*attr*_ and has an attractive valley in between these two length scales with the magnitude ε_*attr*_. For all simulations we used the parameters ε_*attr*_ = 0.11*kT* and *σ*_*attr*_ = 1.5*σ*, which resulted in a relatively weak micro-phase separation corresponding to experimental Hi-C compartmental scores observed here and previously^84^.

Active loop extrusion was added in two steps: (i) one-dimensional extrusion process (stochastic binding and active extrusion) followed by (ii) incorporation of the loops into the three-dimensional polymer simulation. For the one-dimensional loop extrusion, we preset the locations of the TAD boundaries at the positions inferred by calling insulation minima on the Late G1 DMSO R1 + R2 combined experimental Hi-C matrix for chromosome 14 (see Hi-C Insulation Profiles methods section for details). Then the extrusion between the boundaries was performed with some small permeability of the TAD boundary (probability for an extruder to bypass the boundary) *p* ≈ 0.1, which emulated stochasticity of CTCF binding to chromatin. The resulting average loop size was *λ* ≈ 140*kb* and the average gap size between the loops was *g* ≈ 84*kb*, corresponding to ∼62.5% coverage of the chain by loops. This ratio of 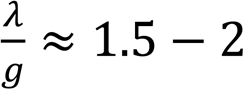 matches the previously inferred density of loops by an analytical model from various mammalian cell types ^31^. Cohesin motors were simulated as stochastically exchanging with the chain; loop sizes were linearly growing with time while the cohesins were bound, unless blocked by the boundaries. In the three-dimensional simulation the sampled configurations of loops for each replicate were modeled as additional harmonic constraints with the same parameters as the polymer bonds. The configuration of loops in the three-dimensional polymer simulation was renewed every 4 seconds, resulting in extrusion rate of 1kb/s, typical for cohesin *in vitro* ^85–87^. The chosen parameters resulted in generation of TADs on the averaged (over 32 replicates and 1 hour of time) contact maps at the positions preset by the experimental boundaries, and were similar to TADs observed on experimental Hi-C maps.

##### Calibration of timescales in simulations

The timescales in simulations were calibrated to match the experimental timescales. For that, we computed the mean-squared displacements (MSD) of the beads ⟨*R*^2^(*t*)⟩, averaged over different beads in the chain, as a function of time (Figure S6H). The MSDs were measured in separate short-time simulations starting from the canonical interphase state with compartments and cohesin loops (late interphase, i.e. ∼8 hours of the two-stage mitotic exit from the knotted state with duration of the first stage T=10min). As expected, at times smaller than the diffusion time between the neighboring Kuhn segments, *t* < *τ*_0_, the dynamics is diffusive, while at larger times, *t* > *τ*_0_, the MSD follows the Rouse law, ⟨*R*^2^(*t*)⟩ = Γ*t*^0.5^. While experimental values of the Rouse diffusion coefficient Γ vary between the organisms and experiments, they all are of the order of 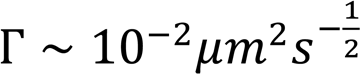. Thus, the conversion of timescales between simulations and experiments was performed by matching the values of the Rouse diffusion coefficient.

Our analysis of times in simulations and experiments further allows to estimate the value of the microscopic Rouse time for chromatin *τ*_0_ ≈ 0.2 − 0.3*s*, which is inaccessible in experiments. This is done by matching the displacements of the beads to the spatial size of the Kuhn segment 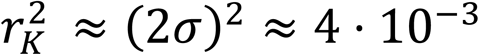 *μm*^2^ at short times. At large times we observe a crossover from Rouse MSD to ⟨*R*^2^(*t*)⟩ ∼ *t*^*z*^ with *z* ≈ 0.3, which is the result of the topologically-constrained “crumpled” chains. Several theoretical predictions of the exponent *z* for crumpled chains have been previously suggested by theories: the generalized Gaussian model for the compact chain with the fractal dimension *d*_*f*_ = 3 suggested *z* = 0.4 ^88, 89^ the fractal loopy globule model had *z* ≈ 0.29 ^90^ and the annealed lattice animal model predicted *z* ≈ 0.26 ^91^. The crossover time from Rouse to crumpled dynamics in our simulations is at *τ_e_* ≈ 10 − 20*s* and the corresponding spatial size *r_e_* ≈ 200*nm* or ≈ 10 randomly packed Kuhn segments. This gives an estimate for the dynamic entanglement length *N*_*e*_ ≈ 80*kb*, which agrees with the previous estimates ^59^. In particular, it further suggests that the cohesin loops are unentangled (*N*_*e*_ < *λ*), giving rise to the effect of dilution of entanglements upon folding of chromosomes into loops ^31^.

As an independent analysis of timescales, we turn to a *characteristic time of the mitotic exit* as the time needed to reach the topological steady state during the Stage I of the two-stage model (Figure 6D). Namely, we observe that in *T* = 10 − 20 minutes a mitotic chain becomes swollen under high activity of Topo II to the steady state with largely decatenated loops (Figure 6E). This time corresponds to the biological time of transition from anaphase to telophase ^40^. Furthermore, we note that the time of this transition matches the residence time of condensin II on chromosomes ^72, 73, 92^. Therefore, setting up the time scales in simulations according to the Rouse diffusion coefficient Γ ≈ 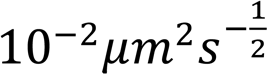 results in the decatenation kinetics with the characteristic time *T* similar to the residence time of condensin II. This suggests that in the real cell the unbinding kinetics of condensin during anaphase-telophase transition ensures the condensin loops have sufficient time to decatenate, additionally justifying the two-stage model of the mitotic exit.

### QUANTIFICATION AND STATISTICAL ANALYSIS

#### Flow Cytometry Analysis

Cell cycle data were analyzed using FlowJo V10.8.1. Viability gates using forward and side scatter area were set on the Async. sample in each experiment, and were applied to all of the samples within the set. Doublet discrimination was then performed using forward scatter area vs forward scatter height, or forward scatter area vs forward scatter width. DNA content was plotted as a histogram of the PI channel area, the exact channel settings varied depending on the instrument used. No compensation was required. G1, S, and G2/M gates were set manually on the Async. sample and applied to all of the samples within the set to obtain the percentage of cells in each state throughout the experiment. Barplots of cell cycle state show the % of single cells with G1, S, or G2/M DNA content as the mean of all replicates, with values from individual replicates as overlaid points.

#### Hi-C Analysis

##### Read Mapping

Hi-C data processing was performed as previously described ^39^. Hi-C PE50 FASTQ sequencing reads were mapped to the hg38 human reference genome using the nextflow based distiller-nf pipeline, followed by filtering to retain only reads with mapping quality (MapQ) of 30 or higher, and removal of duplicate reads. The resulting filtered reads are referred to as valid pairs ^93^. The number of valid pairs was normalized within each experiment using cooltools (v0.5.1) random-sample with the –exact argument on 1kb resolution .cool files ^94^. MultiQC was used for quality control of the Hi-C libraries ^95^. Read normalized valid pairs were binned into .mcool formatted contact matrices at 1kb, 5kb, 10kb, 25kb, 50kb, 100kb, 250kb, 500kb, and 1Mb using the python package cooler (v0.8.11), and were iteratively balanced (ICEed), while masking the first two bins at the diagonal, which can contain artifactual ligation junctions ^96, 97^. Hi-C analysis was performed using custom python based Jupyter notebooks utilizing Hi-C specific python packages such as cooler (v0.8.11), cooltools (v0.5.1), bioframe (v0.3.3) and pairtools (v0.3.0) ^96–100^. Other python packages used included pandas (v1.4.2), numpy (v1.22.3), scipy (v1.8.0), scikit-image (v0.19.2), seaborn (v0.11.2) and matplotlib (v3.5.2) ^101–108^. For HeLa experiments, heatmaps are the q arm of Chr14 binned at 250kb with iterative correction, with Eigenvector 1 plotted below each heatmap, and read normalized between samples within each experiment. For Rad21 degradation experiment heatmap is Chr5:0-40Mb, binned at 250kb with iterative correction and read normalized between samples. Replicates (where available) are combined in heatmap visualizations. Examples of custom python scripts used for Hi-C analysis and visualization is available at https://github.com/dekkerlab/Topo-II-Inhibition-Manuscript/HiC.

##### Distance Decay

Contact frequency (*P*) as a function of genomic distance (*s*) and the derivative (slope) of this *P(s)* curve were calculated using cooltools compute-expected followed by cooltools logbinned-expected using only intrachromosomal reads from the read normalized cooler files, binned at 1Kb ^94^. For Hi-C from HeLa S3 cells, only chromosomes without major translocations were used (chr4, chr14, chr1, chr18, chr20, chr21) ^41^.

##### Insulation Profiles

Diamond insulation score was calculated on 10kb binned .cool files using the cooltools API function calculate_insulation_score, with a 250kb diamond sliding window ^94^. Domain boundaries were found by locating minima in each profile, and thresholding using skimage threshold_otsu ^107^. Boundaries were then filtered to remove boundaries that overlapped with changes in compartment identity, to analyze TAD only boundary strength. Aggregate insulation plots were made by plotting the average insulation in 500kb windows around all called TAD boundaries in each Hi-C library ^105^.

##### Compartment Analysis

Active and inactive compartment regions were identified using eigenvector decomposition on 100kb or 250kb binned Hi-C .cool files using cooltools, as previously described ^39, 94, 109^. The eigenvector was phased such that positive values corresponded to the more gene-dense regions of the genome (the ‘A’, or active compartment), while negative values corresponded to the gene-poor regions of the genome (the ‘B’ or inactive compartment). To measure the strength of compartmentalization, we used cooltools saddleplot analysis on observed/expected Hi-C data at a range of distances, as previously described, where the expected matrix corresponds to the average distance decay ^39, 94, 109^. Observed/expected matrix bins within each distance band were sorted and aggregated into 50 bins according to their eigenvalue. Strength of compartmentalization for each distance was calculated as the ratio of AA/AB or BB/AB interactions, where AA is the average of the corner 10 bins with positive, positive eigenvalues, BB is the average of the 10 corner bins with negative, negative eigenvalues, and AB is the average of the 10 bins in the corner with positive, negative eigenvalues. Bargraphs show the mean of biological replicates, and scatterplot overlay shows values from individual replicates.

##### Loop pileups

We called loops on deep HeLa S3 (4DNFIBM9QCFG) or HCT116 + Rad21-mAC Untreated (4DNFIFLDVASC) Hi-C data from the 4DN data portal ^61^, using cooltools call-dots for the loop pileup analyses, with cooltools v0.4.0 ^110^. For the HeLa S3 data, we obtained 13385 total dots, 2359 on the structurally intact HeLa S3 chromosomes. For the HCT116 + RAD21-mAC data we obtained total 3030 dots on all chromosomes. To plot the aggregate looping interactions, we used cooltools snipping to aggregate 10kb cooler files at the intersection of the two loop anchors, and plotted the average observed/expected ratio for each sample ^94^. Replicates were combined for visualization. Statistical details of each experiment can be found in the figure legends.

#### MC-3C Analysis

##### CCS calling, demultiplexing, fastq extraction

For replicates 1 and 2, PacBio SMRTTools version 8.0.0 was used for consensus consensus sequence (CCS) calling, demultiplexing, and fastq extraction. For replicate 3, PacBio SMRTTools version 10.1.0 was used for these analysis steps.

##### Mapping and Annotation

Reads in fastq format were mapped to the hg38 genome assembly downloaded from ENCODE (GRCh38_no_alt_analysis_set_GCA_000001405.15.fasta.gz) using minimap2 (v2.17) with the option *--secondary=no* ^111, 112^. Mapped C-walks were then annotated both at the fragment and the C-walk level for further filtering and analysis, including number of fragments per C-walk, number of chromosomes and compartment types visited, proximity of each fragment to the largest step in each C-walk, the size of the largest step for each C-walk, the distance of each direct pairwise interaction, the total span of each C-walk, and other features (see examples of custom python scripts used for mapping and secondary analysis on Github (https://github.com/dekkerlab/Topo-II-Inhibition-Manuscript/tree/master/MC3C) for further details). Fastq files containing CCS reads and tables of annotated C-walks are included on GEO.

##### QC

Mapped and annotated reads were filtered to include only reads where all fragments had a mapping quality >59, visited <3 chromosomes, were in regions of the genome called as A or B compartments by Hi-C, and >80% of the entire read was aligned to the genome. For the HeLa S3 libraries, only the chromosomes that have been determined to be minimally translocated were analyzed, as for the Hi-C experiments (chr4, chr14, chr1, chr18, chr20, chr21) ^41^. In addition, except when analyzing raw read length and fragment number for each library, all analyses were performed on the first 6 fragments of all C-walks >= 6 fragments.

##### Descriptive plots

Density plots of C-walk span, direct pairwise interaction distance, and read length were made using the kdeplot function from the python package seaborn (v0.11.2) ^108^. Step style density plot of fragment number per read was made using python package seaborn (v0.11.2) ^108^. Stacked barplots of C-walk type by chromosome number and compartment identity were made using DataFrame.plot function from the python package pandas (v1.4.2) ^101^.

##### Intermingling Metric Analysis

The Intermingling Metric (IM) was calculated using custom python scripts on either real or permuted C-walks. Permuted C-walks were made by shuffling the order of the first 6 fragments of C-walks >= 6 fragments 100 times for each library. To calculate intermingling, a sliding window algorithm was used on C-walks which interacted with only two regions on a single chromosome, as defined by all fragments being within ½ of the size of the largest step from either side of the largest step in the C-walk. C-walks were further filtered to only include C-walks where 5 fragments were close to side 1 of the largest step, and 1 fragment was close to side 2 of the largest step, allowing for either 1 or 2 steps between the two sides. The IM was calculated by binning C-walks with similar sizes of largest steps together depending on the window size (4Mb or 12Mb), and calculating the fraction of C-walks within each window with 2 steps between the two sides of the largest step out of the total number of this type of C-walk. The window was then slid by 1Mb steps to 30Mb separation total. We used seaborn relplot to display the IM within selected window sizes (4Mb and 12Mb) as line plots for each pair of treatments showing the average IM (solid line) and the 95% CI of 3 biological replicates (surrounding shaded area), along with the average IM from permuted walks (3 replicates x 100 permutations each) (dashed line) with the 95% CI from all 300 permutations (surrounding shared area) ^108^. To ensure that the large steps between the interacting domains result are true contacts and not random ligations, we analyzed *P(s)* plots for either all pair-wise interactions from these two region C-walks, or the largest step from each two region C-walk, and found that the *P(s)* plots for both types of interactions were similar, with both displaying distance dependent decay in frequency, indicating that the largest steps follow the expected contact frequency, and therefore are not random ligations (Figure S4L). Statistical details of each experiment can be found in the figure legends.

#### Microscopy Analysis

##### Cell Cycle Staging

Cell cycle stage of wide-field immunofluorescence images was analyzed by blinding the images and manually classifying each cell into a cell cycle category based on morphological features using Nikon Elements software.

##### Euchromatin and Heterochromatin Localization

Confocal images of H3K27ac (red, euchromatin) and H3K9me3 (green, heterochromatin) localization were analyzed using CellProfiler v4.1.3. For each Z-slice, images were first segmented using the DAPI signal to identify nuclei, with object tracking to link nuclei between Z-slices for each image based on location. Next, H3K9me3 and H3K27ac objects were identified within each Z-slice for each nucleus. To avoid out of focus images, and to obtain one representative cross-section of each nucleus, one Z-slice per nucleus was considered for further analysis, which was selected by having the largest nuclear area, H3K9me3 and H3K27ac object counts between the 20% and 90% quantiles within each experiment, nuclear compactness < 90% quantile, and area solidity > 10% quantile. Within this representative Z-slice, the % of H3K27ac objects containing H3K9me3 objects, the radial distribution of each signal in 10 radial bins within each nucleus, and the number and average area of H3K9me3 objects per nucleus were calculated. One representative Z-slice near the center of each nucleus is shown in the figure, autoscaled within each channel of each image to the maximum dynamic range. Boxplot quantification of the confocal microscopy experiments in Figure 2 show min, 25%, 50%, 75%, max. Scatterplot overlay shows individual values for each nucleus. q-values shown on graph from 2-way ANOVA analysis with multiple comparison correction using false discovery rate (FDR = 0.05) using the method of two-stage linear step-up procedure of Benjamini, Krieger and Yekutieli with one family per experimental replicate. Statistical details of each experiment can be found in the figure legends.

##### Texture Analysis

Single Z-slice confocal images from approximately the center of each nucleus were analyzed for DNA and TOP2A-Venus signal contrast using CellProfiler (v4.1.3) to segment nuclei using DAPI signal, and save all channels for each nucleus separately ^113^. Representative images in figure were individually scaled to the full dynamic range within each channel. Segmented images were then used to calculate Haralick features using the python library mahotas (v1.4.11) ^54^. Contrast at a distance of 10 pixels (1um) averaged across all 4 directions for each nucleus was used to compare the textures of DNA (DAPI) and TOP2A-Venus between samples. Statistical analysis was performed using 2-way ANOVA in Graphpad Prism. q-values shown on graph from 2-way ANOVA analysis with multiple comparison correction using false discovery rate (FDR = 0.05) using the method of two-stage linear step-up procedure of Benjamini, Krieger and Yekutieli with one family per experimental replicate. Boxplot shows min, 25%, 50%, 75%, max. Scatterplot overlay shows individual values for each nucleus. Statistical details of each experiment can be found in the figure legends. CellProfiler Pipelines and python code used for image analysis are available in the BioStudies database (https://www.ebi.ac.uk/biostudies/) ^79^.

#### Simulation Quantification

##### Cis/Trans Ratio

The chromosomal territoriality is computed as the ratio of the total number of cis contacts (of the replicate with itself) and the total number of trans contacts (number of contacts of the replicate with all other replicates). The contacts are registered for the two values of the contact radius: 3 beads (3*σ* ≈ 90*nm*) and 5 beads (5*σ* ≈ 150*nm*). The error bars in the corresponding figures reflect the uncertainty of the experimental contact radius. The interval of MC-3C cis/trans data for all chromosomes (from the ICRF-193 or DMSO experiment) is shown for each cis/trans plot (data from Fig. S6I).

For the computation of cis and trans contacts, we construct the respective binned contact maps: of a replicate with itself (symmetric) and of a replicate with some other replicate (non-symmetric). Each contract map is binned with the bin size equal to 100 beads (400kb). For computation of the cis contact map, we used the built-in function from polychrom.contactmaps. For the computation of the trans contact map, we used the custom contact finder function, that takes into account all possible contacts of a given bin with the corresponding bins from all other replicates.

##### Scaling plots

The contact probability curves are computed for the logarithmically spaced bins with 10 bins per the order of magnitude. The code is used from polychrom.polymer_analyses. The cutoff radius for the computation is 5*σ* ≈ 150*nm* for the curves with cohesin and condensin. For the chains without loops (i.e. cohesin depletion simulations) the contact radius is increased to 6*σ* ≈ 180*nm* to compensate for the decreased volume density.

To estimate the error for each condition the scaling curves for 8 replicates (independent runs from the corresponding mitotic state) are plotted along with the corresponding mean. The absolute values of the contact probabilities were normalized 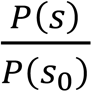 by the initial value *P*(*s*) at *s* = 24*kb*. The slopes were computed as log-derivatives, i.e. 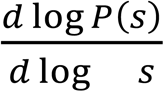, which were further smoothed using the gaussian kernel with the kernel size 1.5.

##### Compartment Saddleplots

For each contact map averaged over 16 replicates we compute the compartmental score and then take the mean and standard deviation with another set of 16 replicates. The corresponding numbers appear on top of the compartment saddleplots in the figures.

To compute the compartment score first we computed the compartment saddleplot as follows. (1) we binned the contact map to the resolution of 400kb; (2) we replaced all zeros in the binned map (no contacts registered) with the minimal possible number of contacts (=1); (3) we computed the observed over expected map using the corresponding tool from mirnylib.numutils; (4) we computed the saddleplot for the resulting map in the specified range of genomic distances; (5) due to computational error the step (4) should be realized several times, and then the compartmental score defined as (AA+BB)/(2*AB) is computed. Here AA is the number of A-A contacts, BB is the number of B-B contacts and AB is the number of A-B contacts, according to the first non-trivial eigenvector of the observed over expected matrix. The number of contacts of the corresponding type is take from the 20% corner of the saddle plot matrix.

##### # catenations per loop

Catenations between the condensin loops are quantified using the Gaussian linking number. For a conformation with a set of loops the matrix of pairwise linking numbers between the loops can be computed using the corresponding function from polychrom.polymer_analyses. All non-zero elements in this symmetric matrix reflect a catenation between the pair of loops; in the figures these matrices are presented using spy function from matplotlib, i.e. all non-zero elements are shown as dots.

The mean number of catenations per loop is computed as the average of number of non-zero elements in the rows of the matrix; then, this number is averaged over the mitotic replicates. For the knotted mitotic state, the mean number of catenations equals ∼0.7, while for the unknotted configurations vast majority of loops are non-concatenated with the average of ∼0.028 catenations per loop.

##### Pairwise distance

The distance matrices are computed for the corresponding conformations from simulations. First, the polymer trajectory is binned to the resolution of 40kb. Second, the covariance matrix between the coordinates of the beads is computed. Third, the squared Euclidean distances are computed based on the elements of the covariance matrix. After the averaging over the trajectories and 32 mitotic replicates, the square root is taken from the average matrix for the presentation in Figures 5 and 6. Thus, the colorbar quantifies the average pairwise distances in microns. The gyration squared size of the segments as a function of the genomic length in Figure 5H is computed from the squared distance matrices.

## SUPPLEMENTAL MATERIALS

**Figure S1, related to Figure 1:**
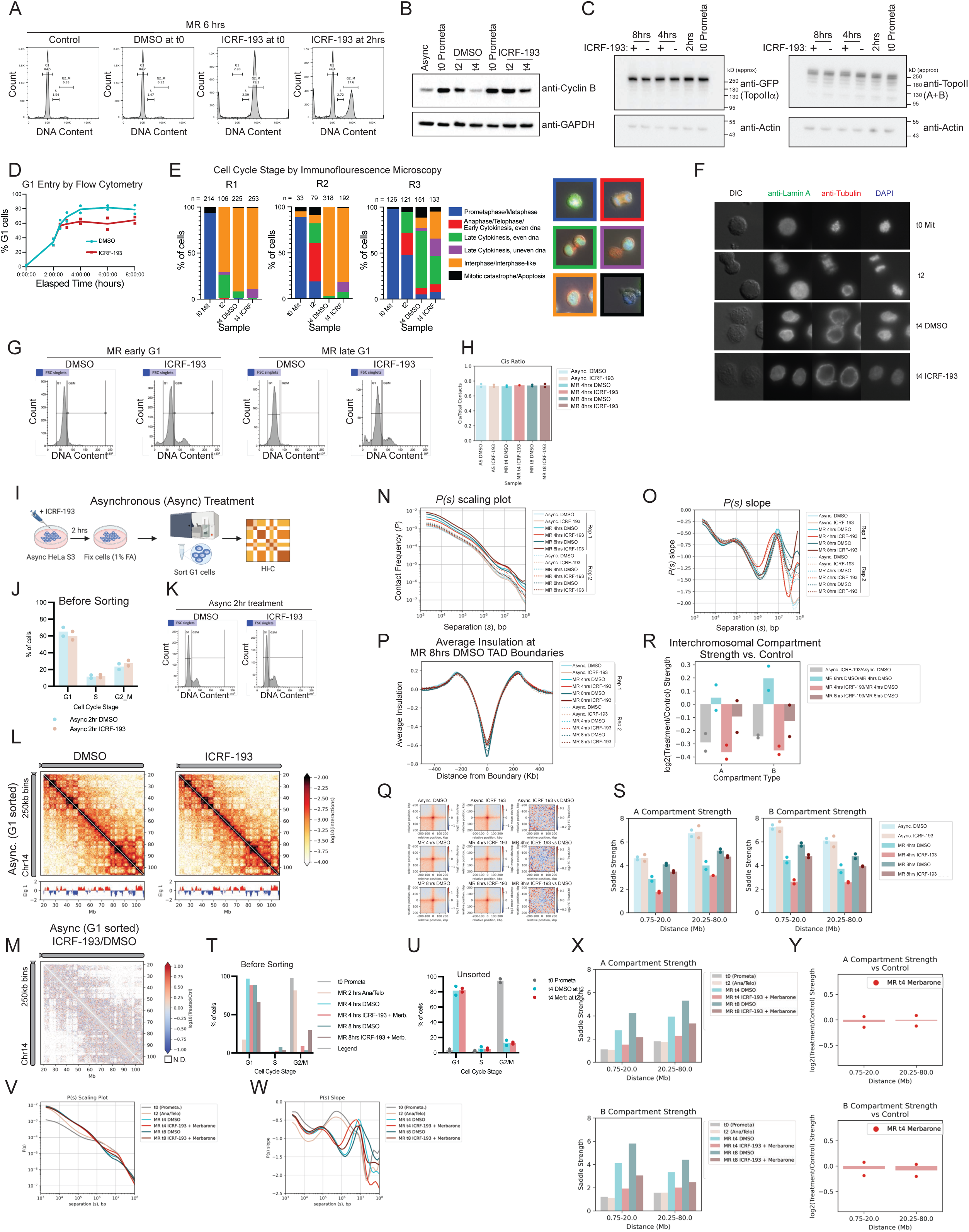
A. PI stained cells were analyzed by flow cytometry. 30uM ICRF-193 blocks G1 entry when added at *t* = 0, but allows G1 entry when added at *t* = 2 hrs. Cells were collected at *t* = 6 hrs. B. Western blot of cyclin B, with GAPDH as a loading control, in asynchronous (Async), prometaphase (t0 Prometa), and mitotic release t = 2hr and t = 4hr HeLa S3 cells. DMSO or 30uM ICRF-193 were added at *t* = 0 (mitotic release), and cells were collected at *t* = 2 or *t* = 4 hrs for western blot analysis. Cyclin B is stabilized with ICRF-193 treatment from *t* = 0 (mitotic release). C. Western blot of Topo II in prometaphase arrest (*t* = 0), two hours after mitotic release (*t* = 2 hrs), and during mitotic release with DMSO or ICRF-193 treatment starting from *t* = 2 hrs after mitotic release and collected at *t* = 4 hrs or *t* = 8 hrs, in the HeLa S3 + TOP2A-Venus cell line. TOP2A levels are analyzed using an anti-GFP antibody, which can recognize the Venus tag. TOP2A and TOP2B levels were also detected using an antibody that recognizes both proteins. Anti-actin is shown as a loading control. Topo II levels are not reduced with ICRF-193 treatment at *t* = 2 hrs post mitotic release. D. Cell cycle progression as measured by DNA content of the total population (adherent + floating cells) when cells are released from prometaphase arrest with or without 30uM ICRF-193 added at *t* = 2 hrs. Analyzed by flow cytometry of PI-stained cells. Samples: DMSO (blue), ICRF-193 (red). Mean of two replicates is shown in line plot, values for individual replicates are shown in scatterplot overlay. E. Stacked bar plots of the distribution of different cell cycle stages and chromosome segregation defects upon ICRF-193 treatment, analyzed by immunofluorescence microscopy (Green channel: Lamin A, Red channel: Tubulin, Blue channel: DAPI). Categories: Prometaphase/Metaphase (blue), Anaphase/Telophase/Early Cytokinesis, with even DNA segregation (red), Late Cytokinesis, with even DNA segregation (green), Late Cytokinesis, with uneven DNA segregation (purple), Interphase/Interphase-like (orange), Mitotic catastrophe/Apoptosis (black). Example images are shown to the right of the plot for each category. Floating and adherent cells were analyzed at *t* = 0 and *t* = 2 hrs, at *t* = 4 hrs only adherent cells were used. F. Representative time-course images from E, replicate 3. G. Example G1 gating for FACS for HeLa S3 cells treated with DMSO or 30uM ICRF-193 from *t* = 2 hrs for MR G1 Hi-C. H. Fraction of interactions in cis (intrachromosomal) for each sample. Samples: Async. DMSO (pale blue), Async. ICRF-193 (pink), MR 4hrs DMSO (bright blue), MR 4hrs ICRF-193 (bright red), MR 8hrs DMSO (dark blue), MR 8hrs ICRF-193 (dark red). Mean of two replicates is shown in the bar graph, with the individual replicate synchronizations shown as scatterplot overlay. I. Schematic of Topo II inhibition by ICRF-193 treatment in asynchronous (Async.) HeLa S3 cells with G1 sorting. Cells were treated for two hours with DMSO or 30uM ICRF-193, and then fixed for Hi-C with 1% formaldehyde (FA). G1 cells were sorted using propidium iodide (PI) staining for DNA content. G1 sorted cells were analyzed by Hi-C 2.0 with DpnII digestion. J. Cell cycle state from flow cytometry profiles of DNA content (PI stain) of Async. HeLa S3 cells with two-hour DMSO or 30uM ICRF-193 treatment before sorting K. Example of G1 gating for FACS to collect G1 cell population from PI-stained cells for Async. G1 sorted Hi-C. L. Hi-C interaction heatmaps of Async. G1 sorted HeLa S3 cells treated with DMSO or 30uM ICRF-193, for the q arm of Chr14. Binned at 250kb with iterative correction and read normalized between samples, two replicates combined. Eigenvector 1 is plotted below each heatmap. M. Hi-C interaction log10 ratio heatmap comparing ICRF-193 treatment to DMSO control Async. samples. Binned at 250kb with iterative correction and read normalized between samples, two replicates combined. N. *P(s)* (scaling) plots of separate replicates of Async. and MR G1 sorted Hi-C of HeLa S3 cells with DMSO or 30uM ICRF-193 treatment. Samples: Async. DMSO (pale blue), Async. ICRF-193 (pink), MR 4hrs DMSO (bright blue), MR 4hrs ICRF-193 (bright red), MR 8hrs DMSO (dark blue), MR 8hrs ICRF-193 (dark red). Replicate 1: solid lines, Replicate 2: dotted lines. O. *P(s)* plot slope of separate replicates of Async. and MR G1 sorted Hi-C of HeLa S3 cells with DMSO or 30uM ICRF-193 treatment. Samples: Async. DMSO (pale blue), Async. ICRF-193 (pink), MR 4hrs DMSO (bright blue), MR 4hrs ICRF-193 (bright red), MR 8hrs DMSO (dark blue), MR 8hrs ICRF-193 (dark red). Replicate 1: solid lines, Replicate 2: dotted lines. P. Average insulation pileup at TAD boundaries, Async. and MR G1 sorted HeLa S3 cells with DMSO or 30uM ICRF-193 treatment Hi-C samples. Two replicates are shown for each sample. Samples: Async. DMSO (pale blue), Async. ICRF-193 (pink), MR 4hrs DMSO (bright blue), MR 4hrs ICRF-193 (bright red), MR 8hrs DMSO (dark blue), MR 8hrs ICRF-193 (dark red). Replicate 1: solid lines, Replicate 2: dotted lines. Q. Aggregate loop analysis, log2 of the mean observed/expected Hi-C interactions in Async. and MR G1 sorted HeLa S3 cells with DMSO or 30uM ICRF-193 treatment samples at 4DN high resolution HeLa S3 Hi-C (4DNFIBM9QCFG) called loops is shown in the left and center columns, respectively, and log2 fold-change of ICRF-193 vs DMSO for each cell cycle condition is shown in the right column. R. Log2 ratio of interchromosomal compartment strength vs control, as indicated. Async. and MR G1 sorted HeLa S3 cells with DMSO or 30uM ICRF-193 treatment Hi-C samples. Mean of two replicates log2 fold change of each indicated comparison is shown in the bar graph, with the log2 fold change from individual replicate synchronizations shown as scatterplot overlay. Comparisons are: Async. ICRF- 193/Async. DMSO (grey), MR 8hrs DMSO/MR 4hrs DMSO (blue), MR 4hrs ICRF-193/MR 4hrs DMSO (bright red), MR 8hrs ICRF-193/MR 8hrs DMSO (dark red). S. Intrachromosomal compartment strength for A or B compartment regions, by distance (0.75-20Mb vs 20.25-80Mb). Samples: Async. DMSO (pale blue), Async. ICRF-193 (pink), MR 4hrs DMSO (bright blue), MR 4hrs ICRF-193 (bright red), MR 8hrs DMSO (dark blue), MR 8hrs ICRF-193 (dark red). Mean of two replicates is shown in the bar graph, with the individual replicate synchronizations shown as scatterplot overlay. T. Cell cycle state from flow cytometry profiles of DNA content (PI stain) of MR HeLa S3 cells with two-hour or four-hour DMSO or 30uM ICRF-193 + 200uM Merbarone treatment from t = 2 hrs MR, before sorting. N = 1. U. Cell cycle state from flow cytometry profiles of DNA content (PI stain) of MR HeLa S3 cells with two-hour 200uM Merbarone treatment from t = 2 hrs MR, unsorted. N = 2. V. *P(s)* (scaling) plots of MR G1 sorted Hi-C of HeLa S3 cells with DMSO or 30uM ICRF-193 + 200uM Merbarone treatment. Samples: t0 Prometa (grey), t 2hrs Ana/Telo (pink), MR 4hrs DMSO (bright blue), MR 4hrs ICRF-193 + Merbarone (bright red), MR 8hrs DMSO (dark blue), MR 8hrs ICRF-193 + Merbarone (dark red). N = 1. W. *P(s)* plot slope of MR G1 sorted Hi-C of HeLa S3 cells with DMSO or 30uM ICRF-193 + 200uM Merbarone treatment. Samples: t0 Prometa (grey), t 2hrs Ana/Telo (pink), MR 4hrs DMSO (bright blue), MR 4hrs ICRF-193 + Merbarone (bright red), MR 8hrs DMSO (dark blue), MR 8hrs ICRF-193 + Merbarone (dark red). N = 1. X. Intrachromosomal compartment strength for A or B compartment regions, by distance (0.75-20Mb vs 20.25-80Mb). Samples: t0 Prometa (grey), t = 2hrs (pink), MR 4hrs DMSO (bright blue), MR 4hrs ICRF-193 + Merbarone (bright red), MR 8hrs DMSO (dark blue), MR 8hrs ICRF-193 + Merbarone (dark red). N = 1. Y. Intrachromosomal AA (top) and BB (bottom) compartment strength log2 ratio compared to control for MR 4hrs by distance, for unsorted HeLa S3 cells treated with 200uM Merbarone from t = 2hrs post mitotic release. N = 2.

**Figure S2, related to Figure 2:**
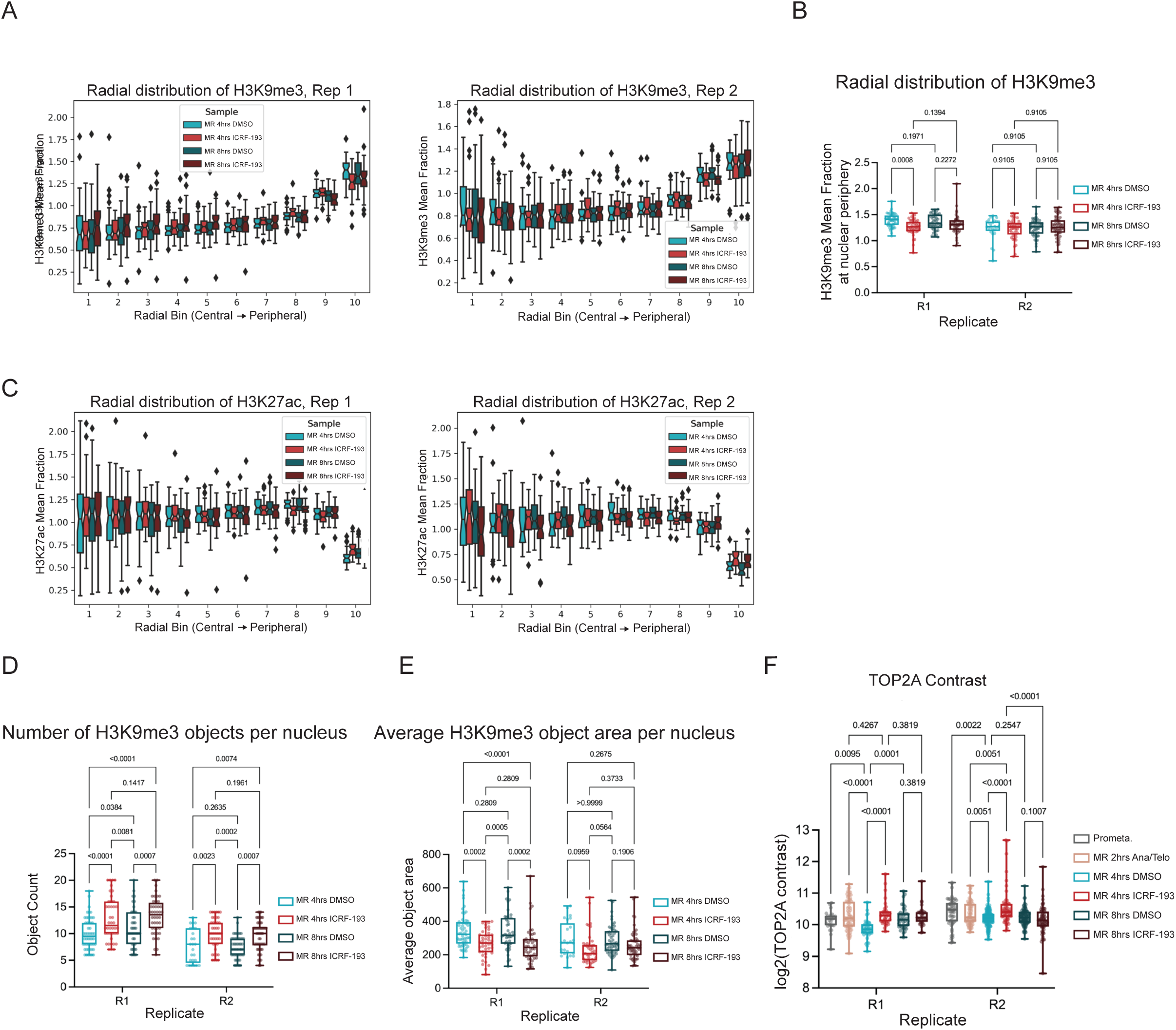
A. Boxplot of radial distribution (10 bins) of Heterochromatin (H3K9me3) (Left: replicate 1, Right: replicate 2) from confocal microscopy experiment. Samples: MR 4hrs DMSO (bright blue), MR 4hrs ICRF-193 (bright red), MR 8hrs DMSO (dark blue), MR 8hrs ICRF-193 (dark red). Boxplot shows 25%, 50%, and 75% quartiles, with whiskers extending to 1.5x the IQR, and outliers beyond that shown as points. Notches indicate 95% CI. B. Boxplot of the mean fraction of H3K9me3 signal in bin 10 (peripheral bin) in both replicate 1 and 2 from confocal microscopy. q-values shown on graph are from 2-way ANOVA analysis with multiple comparison correction using false discovery rate (FDR = 0.05) using the method of two-stage linear step-up procedure of Benjamini, Krieger and Yekutieli with one family per experimental replicate. Boxplot shows min, 25%, 50%, 75%, max. Scatterplot overlay shows individual values for each nucleus. Samples: MR 4hrs DMSO (bright blue), MR 4hrs ICRF-193 (bright red), MR 8hrs DMSO (dark blue), MR 8hrs ICRF-193 (dark red). C. Boxplot of radial distribution (10 bins) of euchromatin (anti-H3K27ac) (Left: replicate 1, Right: replicate 2) from confocal microscopy experiment. Samples: MR 4hrs DMSO (bright blue), MR 4hrs ICRF-193 (bright red), MR 8hrs DMSO (dark blue), MR 8hrs ICRF-193 (dark red). Boxplot shows 25%, 50%, and 75% quartiles, with whiskers extending to 1.5x the IQR, and outliers beyond that shown as points. Notches indicate 95% CI. D. Boxplot of the number of H3K9me3 objects per nucleus in both replicate 1 and 2 from confocal microscopy. q-values shown on graph are from 2-way ANOVA analysis with multiple comparison correction using false discovery rate (FDR = 0.05) using the method of two-stage linear step-up procedure of Benjamini, Krieger and Yekutieli with one family per experimental replicate. Boxplot shows min, 25%, 50%, 75%, max. Scatterplot overlay shows individual values for each nucleus. Samples: MR 4hrs DMSO (bright blue), MR 4hrs ICRF-193 (bright red), MR 8hrs DMSO (dark blue), MR 8hrs ICRF-193 (dark red). E. Boxplot of the average area of H3K9me3 objects per nucleus in both replicate 1 and 2 from confocal microscopy. q-values shown on graph are from 2-way ANOVA analysis with multiple comparison correction using false discovery rate (FDR = 0.05) using the method of two-stage linear step-up procedure of Benjamini, Krieger and Yekutieli with one family per experimental replicate. Boxplot shows min, 25%, 50%, 75%, max. Scatterplot overlay shows individual values for each nucleus. Samples: MR 4hrs DMSO (bright blue), MR 4hrs ICRF-193 (bright red), MR 8hrs DMSO (dark blue), MR 8hrs ICRF-193 (dark red). F. Quantification of the confocal microscopy experiment shown in Figure 2D. Boxplot of TOP2A--Venus signal contrast in each nucleus, at a distance of 10 pixels. q-values shown on graph from 2-way ANOVA analysis with multiple comparison correction using false discovery rate (FDR = 0.05) using the method of two-stage linear step-up procedure of Benjamini, Krieger and Yekutieli with one family per experimental replicate. Boxplot shows min, 25%, 50%, 75%, max. Scatterplot overlay shows individual values for each nucleus. Samples: MR 4hrs early G1 DMSO (bright blue), MR 4hrs early G1 ICRF-193 (bright red), MR 8hrs late G1 DMSO (dark blue), MR 8hrs late G1 ICRF-193 (dark red)

**Figure S3, related to Figure 3:**
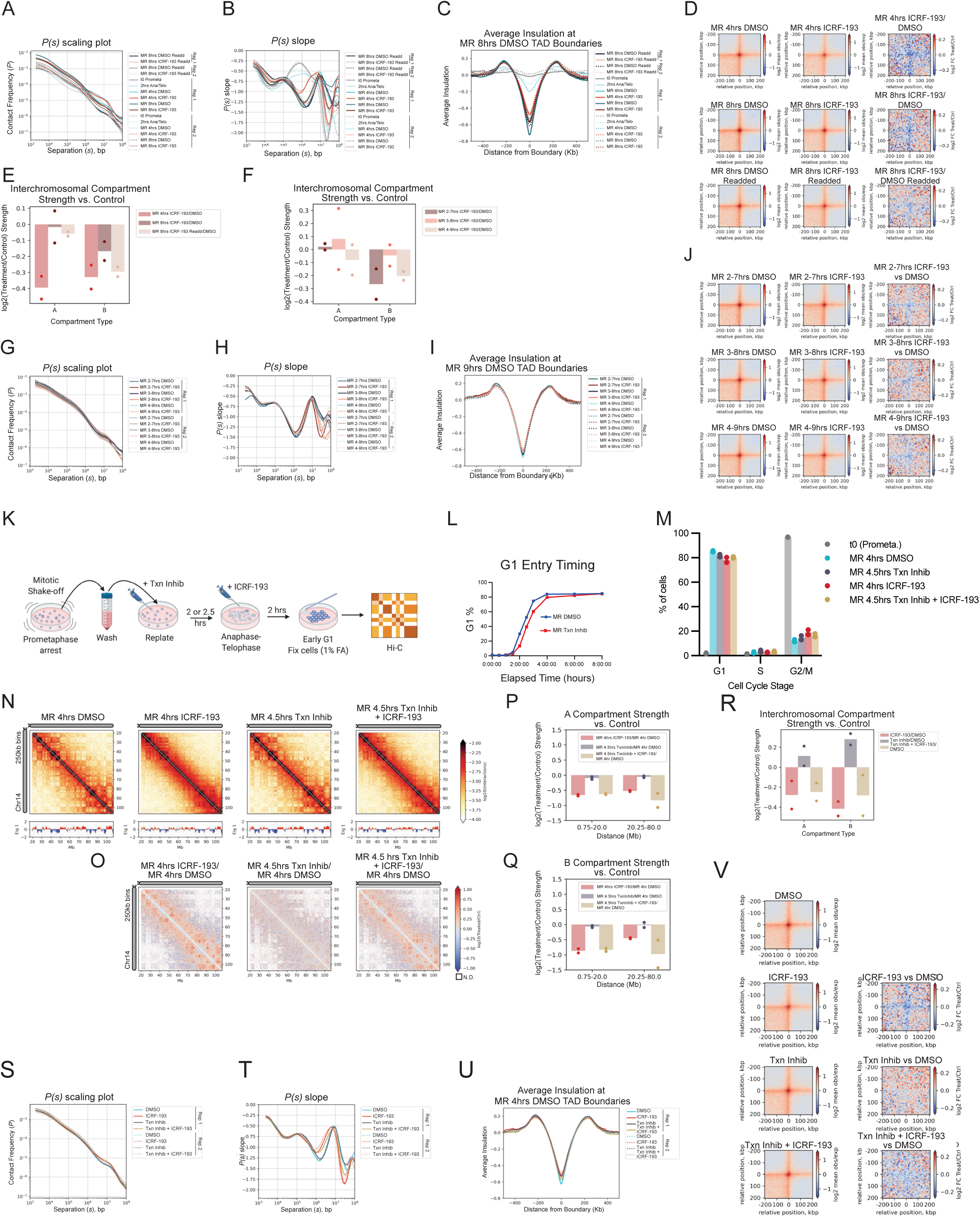
A. *P(s)* scaling plots of separate replicates of MR HeLa S3 cells + DMSO or ICRF-193 starting at *t* = 2 hrs with or without readding every two hours, collected at *t* = 4 hrs or *t* = 8 hrs. Two biological replicates. Samples: MR 8hrs DMSO Readd (black), MR 8hrs 30uM ICRF-193 Readd (pink), t0 prometaphase (grey), MR 2hrs anaphase/telophase (pale blue), MR 4hrs DMSO (bright blue), MR 4hrs 30uM ICRF-193 (bright red), MR 8hrs DMSO (dark blue), MR 8hrs 30uM ICRF-193 (dark red). Replicate 1: solid lines, Replicate 2: dotted lines. Readded and not readded samples are from separate experiments. B. Slope of *P(s)* plot of separate replicates of MR HeLa S3 cells + DMSO or ICRF-193 starting at *t* = 2 hrs with or without readding every two hours, collected at *t* = 4 hrs and *t* = 8 hrs. Two biological replicates. Samples: MR 8hrs DMSO Readd (black), MR 8hrs 30uM ICRF-193 Readd (pink), t0 prometaphase (grey), MR 2hrs anaphase/telophase (pale blue), MR 4hrs DMSO (bright blue), MR 4hrs 30uM ICRF-193 (bright red), MR 8hrs DMSO (dark blue), MR 8hrs 30uM ICRF-193 (dark red). Replicate 1: solid lines, Replicate 2: dotted lines. Readded and not readded samples are from separate experiments. C. Average insulation pileup at TAD boundaries for MR HeLa S3 cells + DMSO or ICRF-193 starting at *t* = 2 hrs with or without readding every two hours, collected at *t* = 4 hrs or *t* = 8 hrs. Two biological replicates. Samples: MR 8hrs DMSO Readd (black), MR 8hrs 30uM ICRF-193 Readd (pink), t0 prometaphase (grey), MR 2hrs anaphase/telophase (pale blue), MR 4hrs DMSO (bright blue), MR 4hrs 30uM ICRF-193 (bright red), MR 8hrs DMSO (dark blue), MR 8hrs 30uM ICRF-193 (dark red). Replicate 1: solid lines, Replicate 2: dotted lines. Readded and not readded samples are from separate experiments. D. Aggregate loop pileups at loop locations called from published high resolution HeLa S3 Hi-C data (4DNFIBM9QCFG). Hi-C data piled up at these locations is from MR with DMSO or ICRF-193 starting at *t* = 2 hrs, either collected at *t* = 4 hrs early G1 without readding or *t* = 8 hrs late G1 with or without readding every two hours. Left column and middle columns show mean log2 observed/expected Hi-C signal for DMSO or ICRF-193 treatments, left column shows log2 fold change of ICRF-193 vs DMSO. Two biological replicates combined. Readded and not readded samples are from separate experiments. E. Log2 ratio of interchromosomal AA and BB compartment strength for MR HeLa S3 cells with 30uM ICRF-193 starting at *t* = 2 hrs with or without readding every two hours, collected at *t =* 4 hrs or *t =* 8 hrs vs DMSO treatment for the same timepoints. Two biological replicates. Samples: MR 4hrs ICRF-193/MR 4hrs DMSO (red), MR 8hrs ICRF-193/MR 8hrs DMSO (dark red), MR 8hrs ICRF-193 Readd/MR 8hrs DMSO Readd (pink). Bar graph shows mean of two biological replicates, scatterplot overlay shows values of individual replicates. Readded and not readded samples are from separate experiments. F. Log2 ratio of interchromosomal AA and BB compartment strength for MR with G1 sorting HeLa S3 Hi-C for 30uM ICRF-193 vs DMSO added at *t =* 2 hrs, *t =* 3 hrs, or *t =* 4 hrs, collected at *t =* 7 hrs, *t =* 8 hrs, and *t =* 9 hrs, respectively (five hour treatment for each). Two biological replicates. MR 2-7hrs ICRF-193/MR 2-7hrs DMSO (dark red), MR 3-8hrs ICRF-193/MR 3-8hrs DMSO (red), MR 4-9hrs ICRF-193/MR 4-9hrs DMSO (pink). Bar graph shows mean of two or three biological replicates, scatterplot overlay shows values of individual replicates. G. *P(s)* scaling plots of separate replicates of MR with G1 sorting HeLa S3 Hi-C with DMSO vs ICRF-193 added at *t =* 2 hrs, *t =* 3 hrs, or *t =* 4 hrs, collected at *t =* 7 hrs, *t =* 8 hrs, and *t =* 9 hrs, respectively (five hour treatment for each). Two biological replicates. H. Slope of *P(s)* plot of separate replicates of MR with G1 sorting HeLa S3 Hi-C with DMSO vs ICRF-193 added at *t =* 2 hrs, *t =* 3 hrs, or *t =* 4 hrs, collected at *t =* 7 hrs, *t =* 8 hrs, and *t =* 9 hrs, respectively (five hour treatment for each). Two biological replicates. Samples: MR 2-7hrs DMSO (dark blue), MR 2-7hrs 30uM ICRF-193 (dark red), MR 3-8hrs DMSO (black), MR 3-8hrs 30uM ICRF-193 (salmon), MR 4-9hrs DMSO (pale blue), MR 4-9 hrs 30uM ICRF-193 (pink). Replicate 1: solid lines, Replicate 2: dotted lines. I. Average insulation pileup at TAD boundaries for MR with G1 sorting HeLa S3 Hi-C with DMSO vs ICRF-193 added at *t =* 2 hrs, *t =* 3 hrs, or *t =* 4 hrs, collected at *t =* 7 hrs, *t =* 8 hrs, and *t =* 9 hrs, respectively (five hour treatment for each). Two biological replicates. Samples: MR 2-7hrs DMSO (dark blue), MR 2-7hrs 30uM ICRF-193 (dark red), MR 3-8hrs DMSO (black), MR 3-8hrs 30uM ICRF-193 (salmon), MR 4-9hrs DMSO (pale blue), MR 4-9 hrs 30uM ICRF-193 (pink). Replicate 1: solid lines, Replicate 2: dotted lines. J. Aggregate loop pileup analysis at loop locations called in published high resolution HeLa S3 Hi-C data (4DNFIBM9QCFG). Piled-up Hi-C data is from MR with G1 sorting HeLa S3 Hi-C with DMSO vs ICRF-193 added at *t =* 2 hrs, *t =* 3 hrs, or *t =* 4 hrs, collected at *t =* 7 hrs, *t =* 8 hrs, and *t =* 9 hrs, respectively (five hour treatment for each). Left and center columns show mean log2 observed/expected for DMSO and ICRF-193 treatment, right column shows log2 fold change of ICRF-193 vs DMSO for each timepoint. Two biological replicates combined. K. Schematic of HeLa S3 mitotic synchronization and release experiment with transcription inhibition and Topo II inhibition. HeLa S3 cells were synchronized in S phase by 24hr treatment with 2mM Thymidine, released 3 hours, then arrested in prometaphase with a 12hr treatment with nocodazole. Mitotic cells were collected by mitotic shake-off, washed, and released into fresh media containing either DMSO or DRB + Triptolide (Txn Inhib). 30uM ICRF-193 was added at *t* = 2 hrs after mitotic release for the ICRF-193 only sample, and *t* = 2.5 hrs after mitotic release for the ICRF-193 + DRB + Triptolide (Txn Inhib) sample. Cells were incubated a further two hours (Early G1, *t* = 4 hrs timepoint for DMSO and ICRF-193 only, *t* = 4.5 hrs timepoint for Txn Inhib and Txn Inhib + ICRF-193 samples) before fixation of adherent cells for Hi-C 2.0 with 1% FA. Chromosome structure was analyzed by Hi-C 2.0 with DpnII digestion. L. Percentage of cells in G1 over time during mitotic exit with DMSO or DRB + TRP (Txn Inhib) treatment from nocodazole wash-out. N = 1. M. Cell cycle profiles by PI staining and flow cytometry of cells described in K. % of single-cells with G1, S, or G2/M cell content is shown (N = 2). N. Hi-C interaction heatmaps of unsorted HeLa S3 cells treated with DMSO, ICRF-193, and/or DRB + TRP (Txn Inhib) at mitotic exit and collected for Hi-C in Early G1 (*t* = 4 hrs or *t* = 4.5 hrs), for the q arm of Chr14. Binned at 250kb with iterative correction and read normalized between samples, two replicates combined. Eigenvector 1 is plotted below each heatmap. O. Hi-C interaction log10 ratio heatmap comparing ICRF-193, DRB + TRP (Txn Inhib), or the combination to DMSO control for each treatment type, 250kb bins. Two replicates combined. P. AA compartment strength log2 ratio compared to DMSO by distance, separated by compartment type, for MR HeLa S3 cells. MR 4hrs early G1 ICRF-193/MR 4hrs early G1 DMSO (red), MR 4.5hrs early G1 Txn Inhib/MR 4hrs early G1 DMSO (purple), MR 4.5hrs early G1 ICRF-193 + Txn Inhib/MR 4hrs early G1 DMSO (gold). N=2. 250kb binned data. Bar graph shows mean of two biological replicates, scatterplot overlay shows values from individual replicates. Q. BB compartment strength log2 ratio compared to DMSO by distance, separated by compartment type, for MR HeLa S3 cells. MR 4hrs early G1 ICRF-193/MR 4hrs early G1 DMSO (red), MR 4.5hrs early G1 Txn Inhib/MR 4hrs early G1 DMSO (purple), MR 4.5hrs early G1 ICRF-193 + Txn Inhib/MR 4hrs early G1 DMSO (gold). N=2. 250kb binned data. Bar graph shows mean of two biological replicates, scatterplot overlay shows values from individual replicates. R. Log2 fold change of interchromosomal AA or BB compartment strength for each treatment vs. DMSO only control. Two biological replicates are shown. Samples: MR 4hrs ICRF-193/MR 4hrs DMSO (red), MR 4.5hrs Txn Inhib/MR 4hrs DMSO (purple), MR 4.5hrs Txn Inhib + ICRF-193/MR 4hrs DMSO (gold). Mean of two biological replicates is shown in bar graph, separate replicates are shown in scatterplot overlay. S. *P(s)* scaling plots of separate replicates of MR *t* = 4 hrs or *t* = 4.5 hrs TRP + DRB or ICRF-193 or combined treated HeLa S3 cells compared to DMSO treated control. Two biological replicates are shown. Samples: MR 4hrs DMSO (bright blue), MR 4hrs 30uM ICRF-193 (bright red), MR 4.5hrs Txn Inhib (purple), MR 4.5hrs Txn Inhib + 30uM ICRF-193 (gold). Replicate 1: solid lines, Replicate 2: dotted lines. T. Slope of *P(s)* plot of separate replicates of MR *t* = 4 hrs or *t* = 4.5 hrs TRP + DRB or ICRF-193 or combined treated HeLa S3 cells compared to DMSO treated control. Two biological replicates are shown. Samples: MR 4hrs DMSO (bright blue), MR 4hrs 30uM ICRF-193 (bright red), MR 4.5hrs Txn Inhib (purple), MR 4.5hrs Txn Inhib + 30uM ICRF-193 (gold). Replicate 1: solid lines, Replicate 2: dotted lines. U. Average insulation pileup at TAD boundaries for MR TRP + DRB, ICRF-193 or combined treated HeLa S3 cells, *t* = 4 hrs DMSO R1 + R2 TAD only boundaries used for pileup locations. Two biological replicates are shown. Samples: MR 4hrs DMSO (bright blue), MR 4hrs 30uM ICRF-193 (bright red), MR 4.5hrs Txn Inhib (purple), MR 4.5hrs Txn Inhib + 30uM ICRF-193 (gold). Replicate 1: solid lines, Replicate 2: dotted lines. V. Aggregate loop analysis, MR TRP + DRB or ICRF-193 or combined treated HeLa S3 cells, AS control HeLa S3 loop locations called using published high resolution HeLa S3 Hi-C data (4DNFIBM9QCFG). Mean observed/expected for DMSO, TRP + DRB, ICRF-193, or TRP + DRB + ICRF-193 treatment shown in left column, log2 fold change of TRP + DRB and/or ICRF-193 vs DMSO is shown in right column for each cell cycle state. Two biological replicates are combined.

**Figure S4, related to Figure 4:**
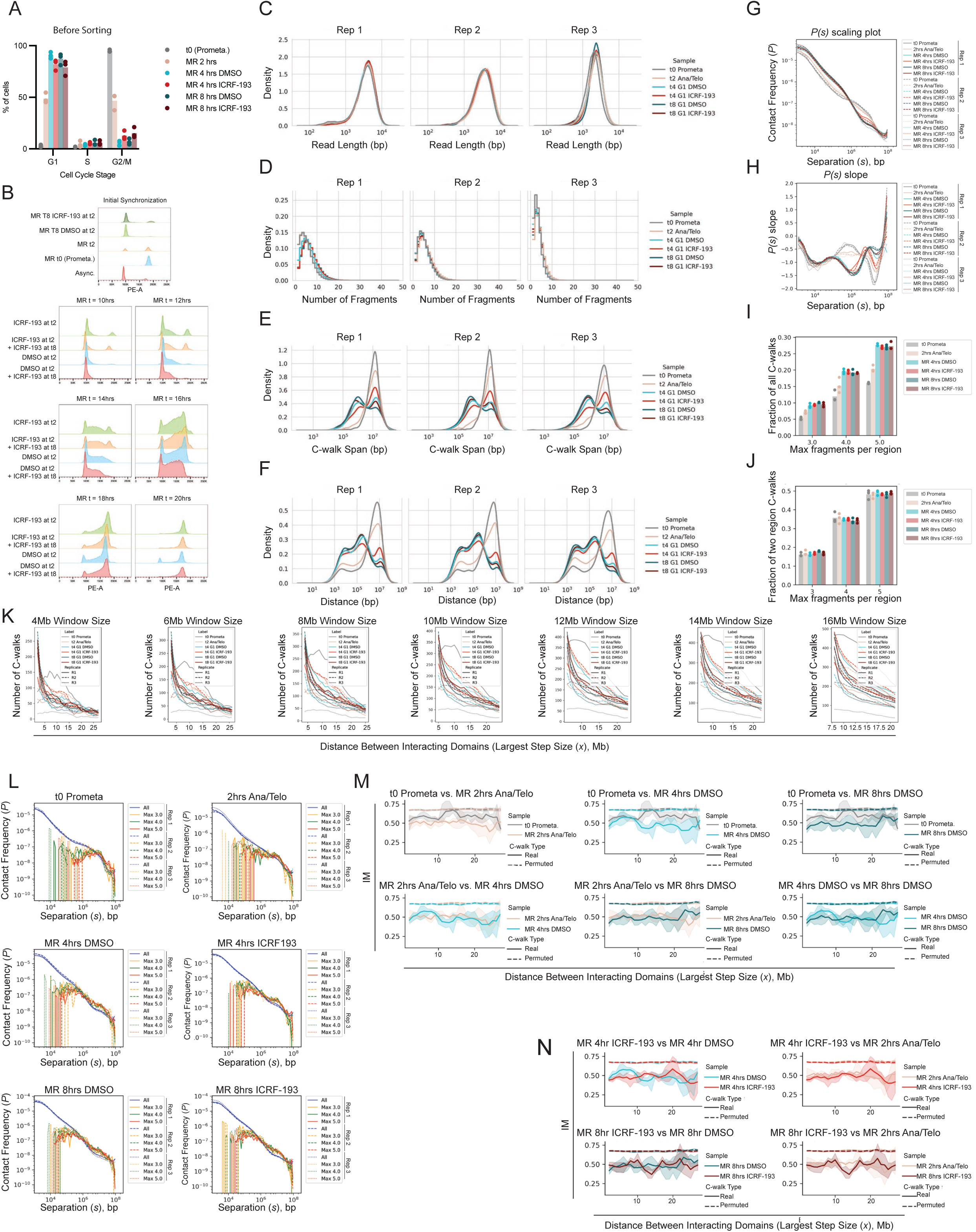
A. Cell cycle profiles by PI staining and flow cytometry of HeLa S3 cells collected for MC-3C before sorting. % of single-cells with G1, S, or G2/M cell content is shown (N = 3). B. Cell cycle profiles to measure S phase progression by PI staining and flow cytometry of HeLa S3 cells synchronized in prometaphase and released into G1, with DMSO or 30uM ICRF-193 added at t = 2hrs post mitotic release, to determine if the decrease in compartment strength observed with ICRF-193 treatment affects DNA replication progression. At 8 hrs (late G1), 30uM ICRF-193 was (re) added to both control and ICRF-193 treated samples as shown. S phase progression was monitored at 10, 12, 14, 16, 18, and 20 hours post mitotic release. C. Density plots of read-length of all aligned MC-3C C-walks for each sample, three biological replicates plotted on separate axes, as indicated. Samples: t0 prometaphase (grey), MR 2hrs Ana/Telophase (pink), MR 4hrs DMSO (bright blue), MR 4hrs 30uM ICRF-193 (bright red), MR 8hrs DMSO (dark blue), MR 8hrs 30uM ICRF-193 (dark red). D. Step style histogram density plot of the number of separate fragments per C-walk for all aligned C-walks in each sample. Three biological replicates plotted on separate axes, as indicated. Samples: t0 prometaphase (grey), MR 2hrs Ana/Telophase (pink), MR 4hrs DMSO (bright blue), MR 4hrs 30uM ICRF-193 (bright red), MR 8hrs DMSO (dark blue), MR 8hrs 30uM ICRF-193 (dark red). E. Density plot of C-walk span for the first 6 fragments of C-walks on one chromosome in A, B, or A and B compartment regions. Each replicate is plotted on a separate axis. Samples: t0 prometaphase (grey), MR 2hrs Ana/Telophase (pink), MR 4hrs DMSO (bright blue), MR 4hrs 30uM ICRF-193 (bright red), MR 8hrs DMSO (dark blue), MR 8hrs 30uM ICRF-193 (dark red). F. Density plot of the distance of direct MC-3C interactions for the first 6 fragments of C-walks on one chromosome in A, B, or A and B compartment regions. Each replicate is plotted on a separate axis. Samples: t0 prometaphase (grey), MR 2hrs Ana/Telophase (pink), MR 4hrs DMSO (bright blue), MR 4hrs 30uM ICRF-193 (bright red), MR 8hrs DMSO (dark blue), MR 8hrs 30uM ICRF-193 (dark red). G. *P(s)* scaling plot, normalized to AUC, of direct MC-3C interactions, replicates plotted separately. Three biological replicates. Samples: t0 prometaphase (grey), MR 2hrs Ana/Telophase (pink), MR 4hrs DMSO (bright blue), MR 4hrs 30uM ICRF-193 (bright red), MR 8hrs DMSO (dark blue), MR 8hrs 30uM ICRF-193 (dark red). Replicate 1: solid line, Replicate 2: dashed line, Replicate 3: dotted line. H. *P(s)* plot slope of direct MC-3C interactions, replicates plotted separately. Three biological replicates. Samples: t0 prometaphase (grey), MR 2hrs Ana/Telophase (pink), MR 4hrs DMSO (bright blue), MR 4hrs 30uM ICRF-193 (bright red), MR 8hrs DMSO (dark blue), MR 8hrs 30uM ICRF-193 (dark red). Replicate 1: solid line, Replicate 2: dashed line, Replicate 3: dotted line. I. Fraction of all first-6-fragment C-walks on one chromosome in A, B, or A and B compartment regions that have are between only two regions, as defined by each region being no more than ½ the size of the largest step in each C-walk, centered on the two sides of the largest step. These two region C-walks are separated by the number of fragments in each region, with the maximum number of fragments in one region shown on the *x* axis. Samples: t0 prometaphase (grey), MR 2hrs Ana/Telophase (pink), MR 4hrs DMSO (bright blue), MR 4hrs 30uM ICRF-193 (bright red), MR 8hrs DMSO (dark blue), MR 8hrs 30uM ICRF-193 (dark red). Average of three replicates is shown in bar graph, separate replicate values are shown in scatterplot overlay. J. Fraction of one chromosome two region C-walks, as defined in I, that have a maximum of 3, 4, or 5 fragments in one region. Samples: t0 prometaphase (grey), MR 2hrs Ana/Telophase (pink), MR 4hrs DMSO (bright blue), MR 4hrs 30uM ICRF-193 (bright red), MR 8hrs DMSO (dark blue), MR 8hrs 30uM ICRF-193 (dark red). Average of three replicates is shown in bar graph, separate replicate values are shown in scatterplot overlay. K. Number of C-walks in each sample at each distance of separation for C-walks used in intermingling analysis, separated by the size of the sliding window. Samples: t0 prometaphase (grey), MR 2hrs Ana/Telophase (pink), MR 4hrs DMSO (bright blue), MR 4hrs 30uM ICRF-193 (bright red), MR 8hrs DMSO (dark blue), MR 8hrs 30uM ICRF-193 (dark red). Replicate 1: solid line, Replicate 2: dashed line, Replicate 3: dotted line. L. *P(s)* scaling plot, normalized to AUC, of either all direct interactions from first 6 fragments of all C-walks (blue line), or only the largest step in 6 fragment C-walks that visit two regions on one chromosome, for walks with a maximum of 3 (orange), 4 (green), or 5 (red) fragments on one side of the largest step. Replicate 1: solid line, Replicate 2: dashed line, Replicate 3: dotted line. M. IM for 4Mb sliding window for cell cycle progression comparisons indicated. Solid line and surrounding shading show the mean of three biological replicates +/- 95% CI. Dashed line and surrounding shading shows mean of 100 sets of permuted/shuffled walks per biological replicate (300 total for each treatment) +/- 95% CI in the surrounding shaded regions. Samples: t0 prometaphase (grey), MR 2hrs Ana/Telophase (pink), MR 4hrs DMSO (bright blue), MR 4hrs 30uM ICRF-193 (bright red), MR 8hrs DMSO (dark blue), MR 8hrs 30uM ICRF-193 (dark red). N. IM for 4Mb sliding window for ICRF-193 vs DMSO and ICRF-193 vs *t* = 2 hrs comparisons, as indicated. Solid line and surrounding shading show the mean of three biological replicates +/- 95% CI. Dashed line and surrounding shading shows mean of 100 sets of permuted/shuffled walks per biological replicate (300 total for each treatment) +/- 95% CI in the surrounding shaded regions. Samples: t0 prometaphase (grey), MR 2hrs Ana/Telophase (pink), MR 4hrs DMSO (bright blue), MR 4hrs 30uM ICRF-193 (bright red), MR 8hrs DMSO (dark blue), MR 8hrs 30uM ICRF-193 (dark red).

**Figure S5, related to Figures 5 and 6.**
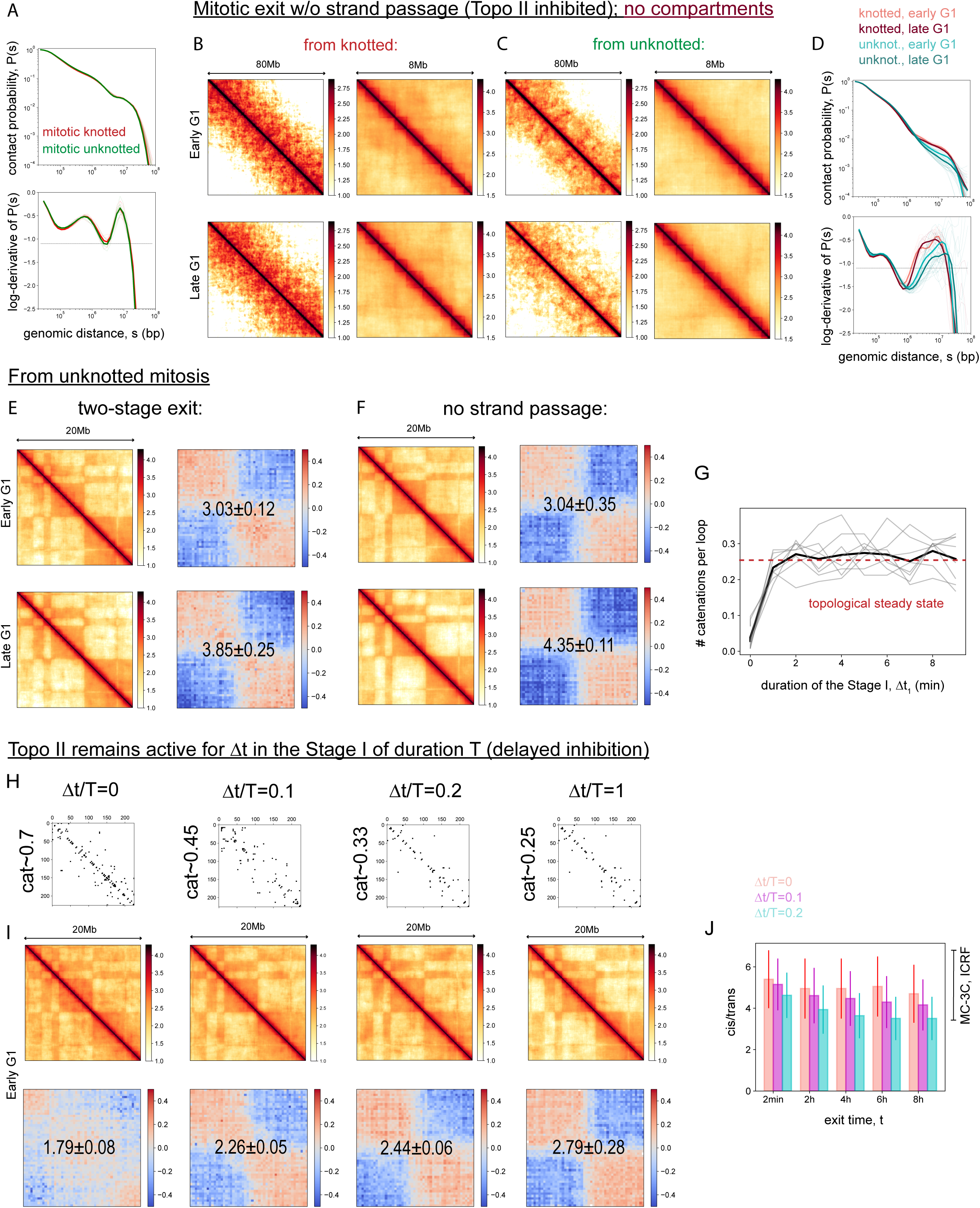
A. Mitotic contact probabilities for the two states: knotted (red) and unknotted (green), as well as the corresponding log-derivatives. Despite different topologies, the two mitotic states have identical contact probabilities. B. Simulated contact maps for the expansion from the knotted mitotic state without compartmental interactions and with inhibited strand passage. Two time points (early and late G1) and two resolutions (80Mb and 8Mb) are shown. C. The same as in A, but the expansion is performed from the unknotted mitotic state. D. Contact probability curves and the corresponding log-derivatives for the expansion out of the two mitotic states without compartmental interaction and with inhibited strand passage. E. Simulated contact maps and the respective compartmentalization saddle plots for the two-stage exit from the unknotted mitotic state. Two time points (early and late G1) are shown. The compartmentalization score in the close band (0.75-20Mb) is indicated on the saddle plots. F. The same as in D, but for the mitotic exit with inhibited strand passage. The data corresponds to Figure 5B,C. G. The number of catenations per loop as a function of elapsed time during Stage I of the two-stage exit out of the unknotted mitotic state. Eight replicates are shown in gray and the averaged curve is shown in bold black. The steady-state value of catenations for the two-stage exit out of the knotted mitosis (Figure 6E) is shown by the dashed red line. H. Simulations of the delayed strand passage inhibition: Topo II activity is inhibited in time Δ*t* after the beginning of Stage I of the total duration T=10 min. Expansion is performed from the knotted initial state. The matrices of pairwise catenations between the mitotic loops at the end of Stage I are shown in the first row for each value of 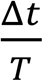; the resulting mean number of catenations per loop is indicated for each of the matrices. I. The contact maps at the early G1 time point are shown for the delayed strand passage inhibition; the corresponding saddle plots with indicated compartmental scores (in the close band, 0.75-20Mb) are shown for each 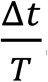. J. Cis/trans ratio for the chains expanded in the delayed strand passage simulation as a function of the elapsed time. The results for three values of the parameter 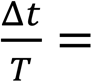 0, 0.1, 0.2 are demonstrated by different colors as indicated. The case of 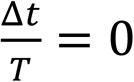 corresponds to the “ideal” Topo II inhibition, shown in Figure 5F in red.

**Figure S6, related to Figure 6.**
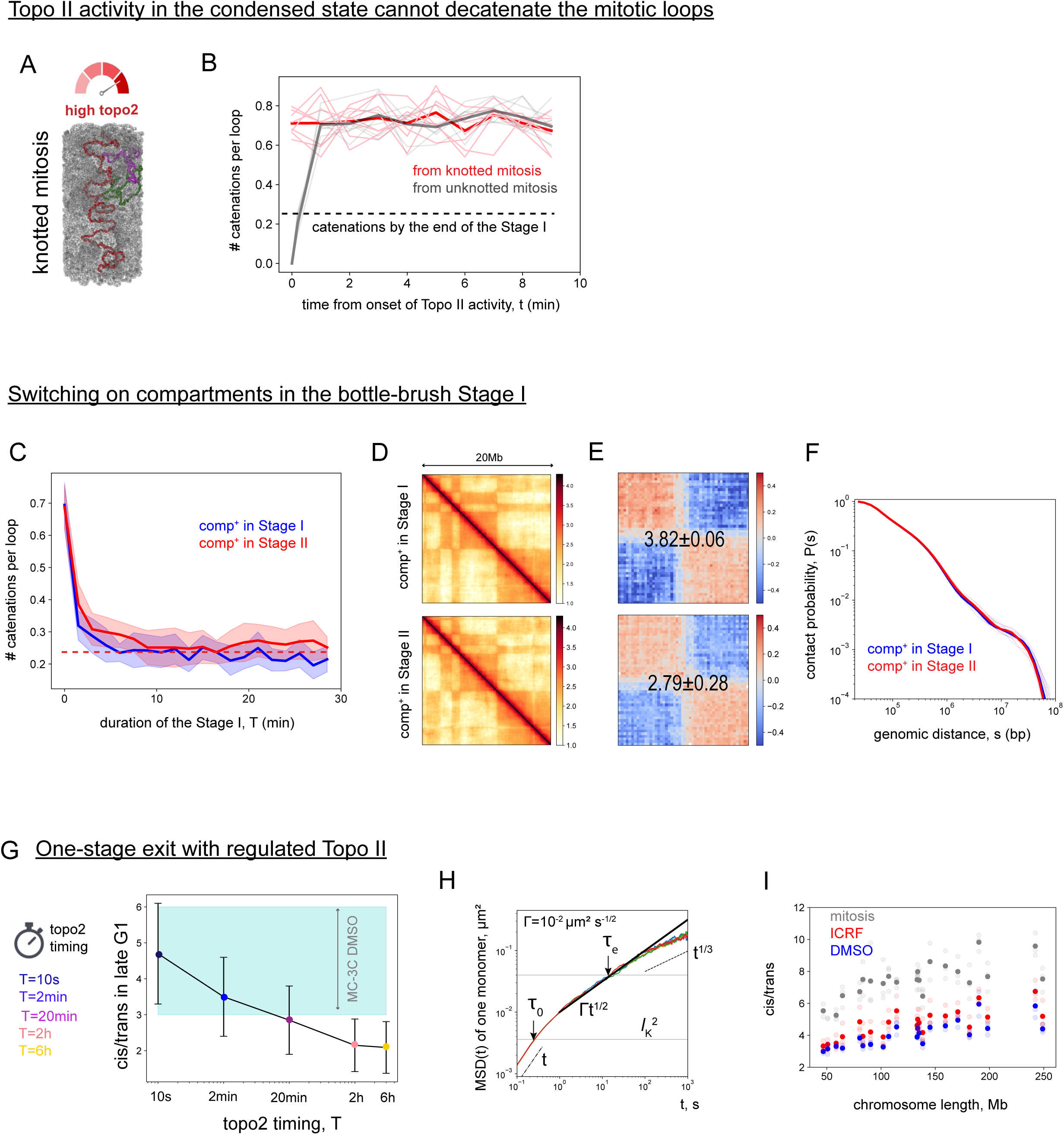
A. Induction of Topo II activity in the condensed knotted mitotic state. Topo II activity (allowed strand passage) is activated in the initial mitotic configuration with catenated loops within the confining cylindrical volume. B. The plot shows the dependence of the number of catenations per loop as a function of duration of Topo II activity for eight replicates of a knotted mitotic chromosome (thin red lines); the averaged curve is shown in bold red. For the comparison a similar set of curves is shown for the unknotted mitotic state (gray lines). Notably, the steady-state level of catenations is independent from the topology of the initial state. The dashed black line indicates the residual level of catenations obtained by the end of Stage I of the two-stage model. C. Simulations of the two-stage exit with the compartments switched on in the beginning of Stage I, i.e. in the bottle-brush stage. The graph compares the evolution of the number of catenations per loop with time in Stage I for the cases with compartmental interactions (blue) and without (red). The latter case is presented in Figure 6E. The curves reflect the averages and the strips correspond to the standard deviation in the sample of 16 replicates. The red dashed line shows the steady-state amount of catenations in the model, when compartmental interactions are added in Stage II. D. Same simulations as in panel C. The respective contact maps for the cases of compartments added in Stage I and Stage II are shown for the early G1 time point. E. The saddle plots for two contact maps from panel D quantify the compartmental strength for the two cases (in the close band, 0.75-20Mb). F. Contact probability curves for the cases of compartments added in Stage I and Stage II as indicated. The contact statistics is not sensitive to the eventual strength of compartments, which agree with the results showin in Figure S5D. G. Simulations of the one-stage exit from the knotted mitotic state with regulated activity of Topo II. A chromosome is expanded through a simultaneous removal of the mitotic loops and cylindrical confinement; compartments and cohesin loops are added from the very beginning; Topo II is active during some time T and then is switched off. The graph shows the cis/trans ratio at the late G1 time point as a function of the Topo II timing, T. While short activity of Topo II is not sufficient to reduce the catenations (see Figure 6E), the graph demonstrates that long timing of Topo II (>20 minutes) results in improbably low levels of territoriality as compared to the experimental range of cis/trans values for the DMSO MC-3C data (Figure S6I). H. Monomer mean-squared displacements (MSD) as a function of time as computed in simulations at the late G1 time point. Four replicates are shown by different colors (red, blue, orange, green). The bold black line corresponds to the Rouse behavior 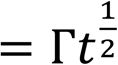 with the diffusion coefficient 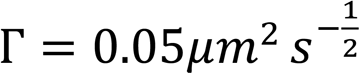. Dashed lines correspond to the short (∼ *t*, diffusive) and long (∼ *t*^0.3^, crumpled) time scaling of the MSD. The upper horizontal line corresponds to the squared spatial size of the entanglement blob, roughly determined as the size of crossover from Rouse to crumpled dynamics. The gray horizontal line corresponds to the squared spatial size of the Kuhn segment and is used to determine the microscopic Rouse time *τ*_0_ ≈ 0.3*s* I. Computed cis/trans ratios on experimental maps, generated from MC-3C experiment, for mitosis (gray), ICRF-193 (red) and DMSO (blue). Three replicates for each experimental point are shown by transparent dots, and their averages are shown by solid dots.

**Figure S7, related to Figure 7:**
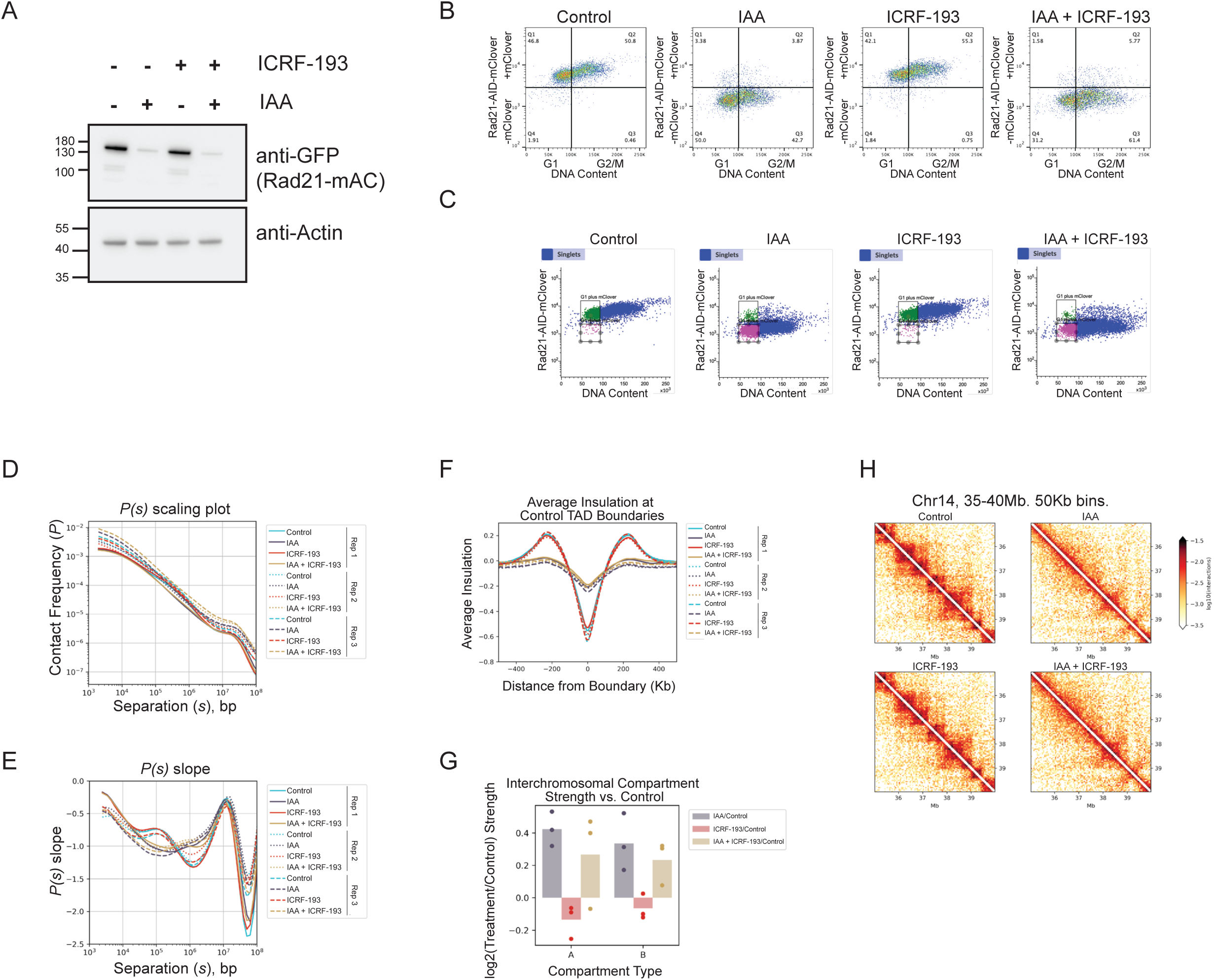
A. Representative western blot of RAD21-mAID-mClover degradation with two-hour 500uM IAA treatment and/or 30uM ICRF-193 treatment in the HCT116 + RAD21-mAC cell line. Anti-GFP was used to detect mClover tagged Rad21, and anti-actin was used as a loading control. B. Representative flow cytometry plots of PI vs mClover (GFP channel), showing shifts from plus to minus mClover signal with two-hour IAA treatment. C. Example of sorting strategy for G1 cells (PI stain, DNA content) plus or minus GFP, (to detect RAD21-AID-mClover). The following populations were collected for Hi-C: Control: G1 + mClover, IAA: G1 - mClover, ICRF-193: G1 + mClover, IAA + ICRF-193: G1 - mClover. D. *P(s)* scaling plot of separate replicates of HCT116 + RAD21-mAC Hi-C. Three biological replicates. Samples: Control (blue), 500uM IAA (purple), 30uM ICRF-193 (red), 500uM IAA + 30uM ICRF-193 (gold). Replicate 1: solid line, Replicate 2: dotted line, Replicate 3: dashed line. E. *P(s)* plot slope of separate replicates of HCT116 + RAD21-mAC Hi-C. Samples: Control (blue), 500uM IAA (purple), 30uM ICRF-193 (red), 500uM IAA + 30uM ICRF-193 (gold). Replicate 1: solid line, Replicate 2: dotted line, Replicate 3: dashed line. F. Average insulation pileup at Control TAD boundaries for HCT116 + RAD21-mAC Hi-C. Samples: Control (blue), 500uM IAA (purple), 30uM ICRF-193 (red), 500uM IAA + 30uM ICRF-193 (gold). Replicate 1: solid line, Replicate 2: dotted line, Replicate 3: dashed line. G. Log2 fold change of interchromosomal AA and BB compartment strength for each treatment vs control from HCT116 + RAD21-mAC Hi-C. Three biological replicates. Samples: IAA/Control (purple), ICRF-193/Control (red), IAA + ICRF-193/Control (gold). Bar graph shows mean of three biological replicates, scatterplot overlay shows individual replicate values. H. TAD level heatmaps for HCT116 + RAD21-mAC Hi-C showing loss of TADs with cohesin degradation. Chr14, 35-40Mb. 50Kb bins.

## Key resources table

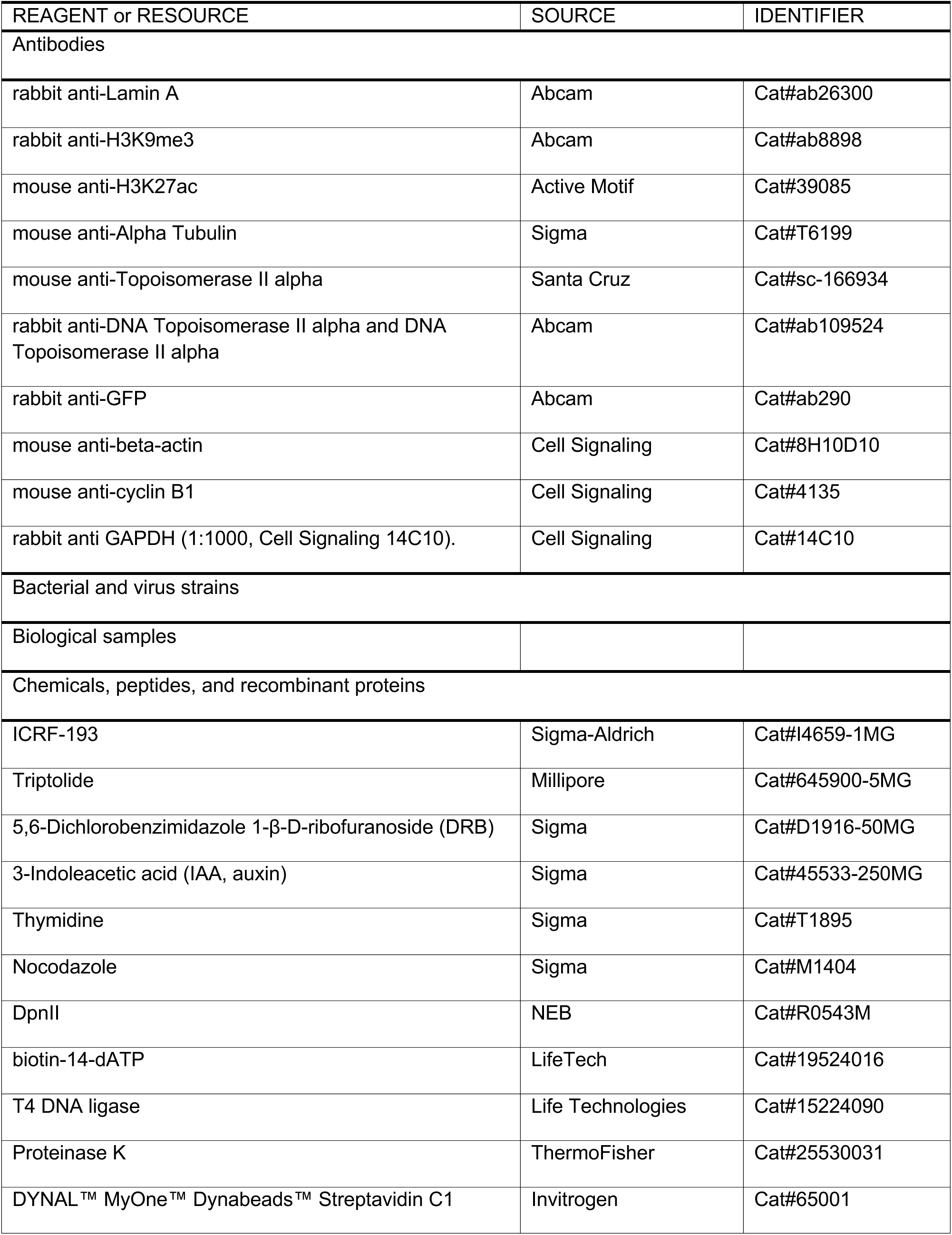
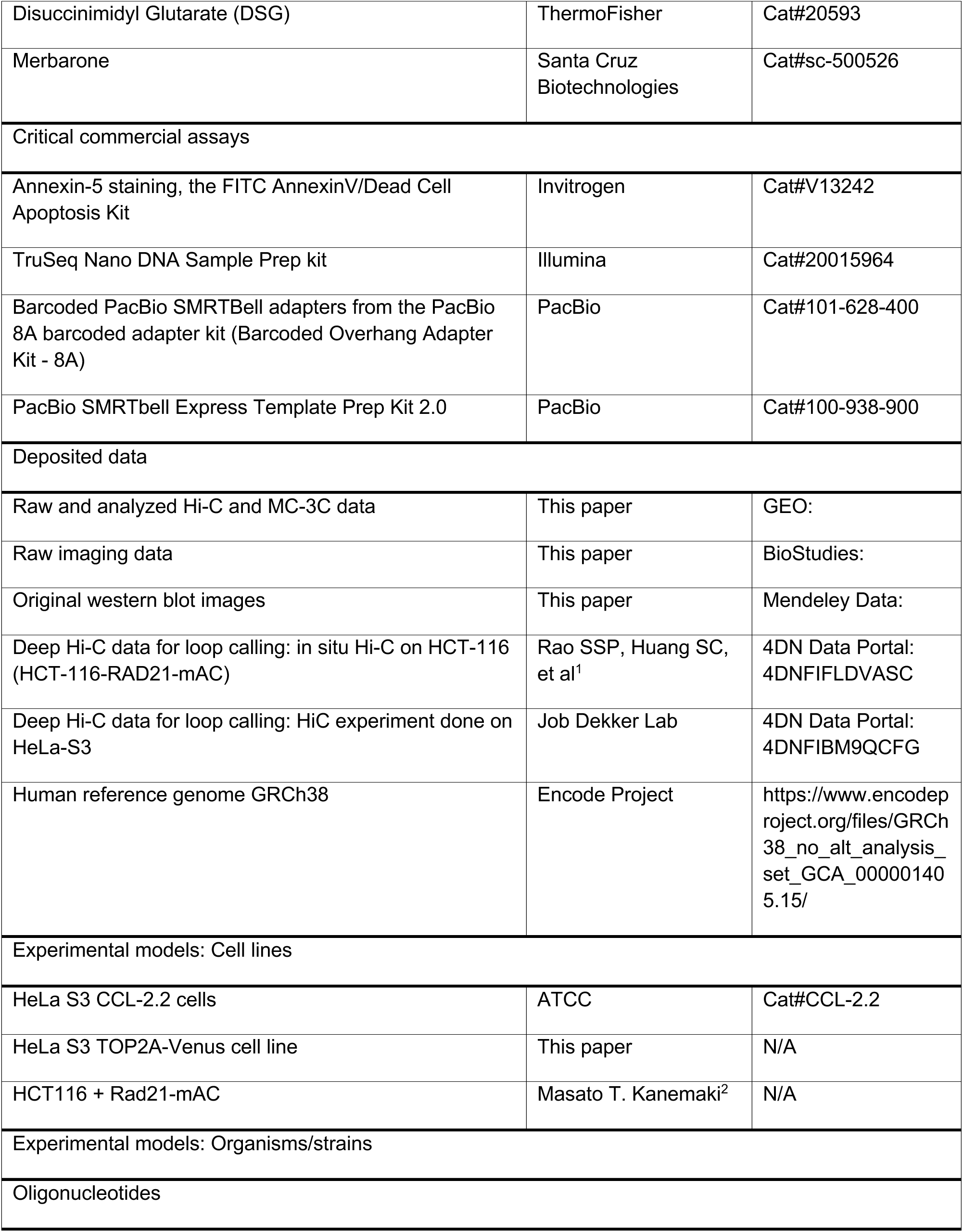
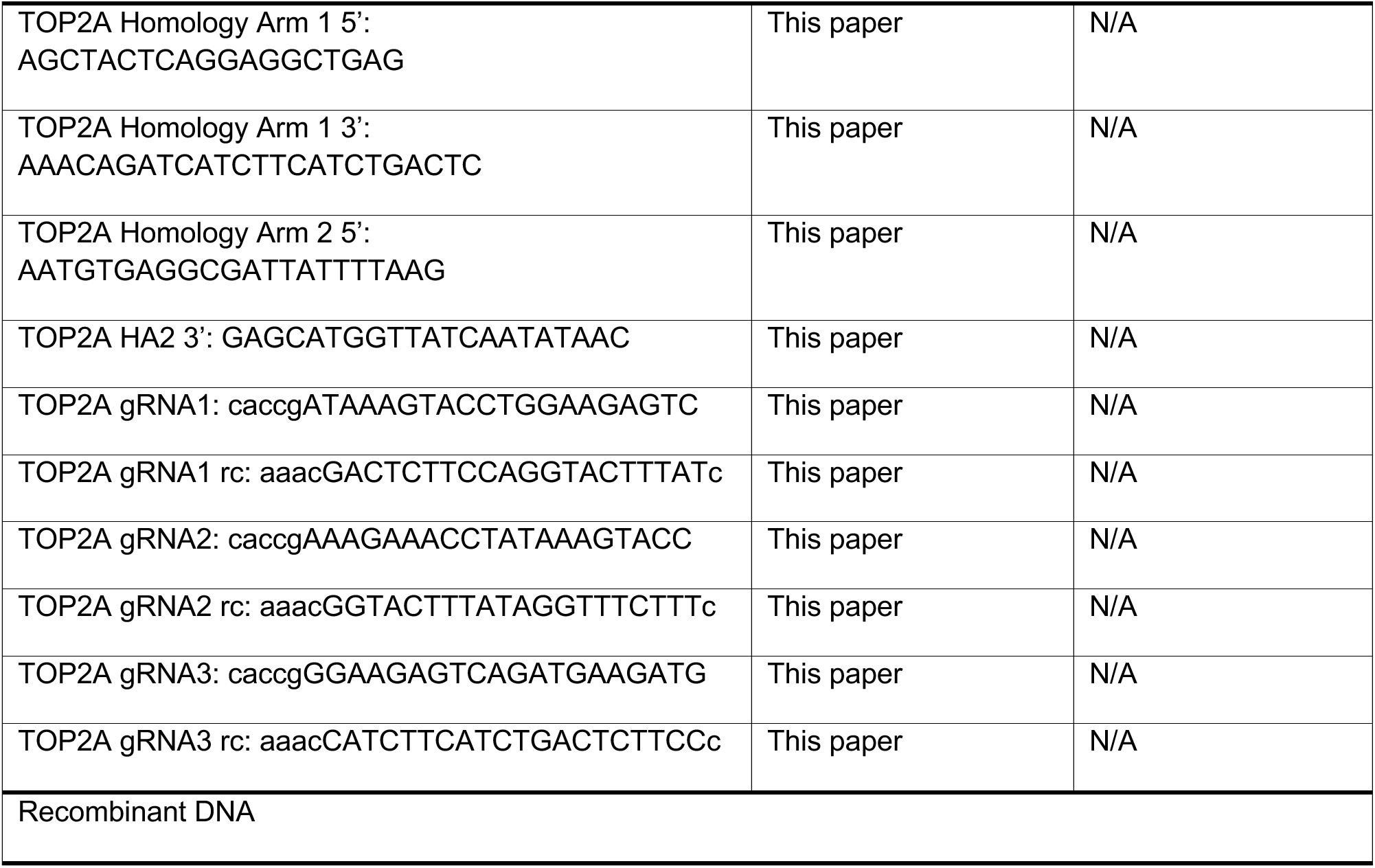
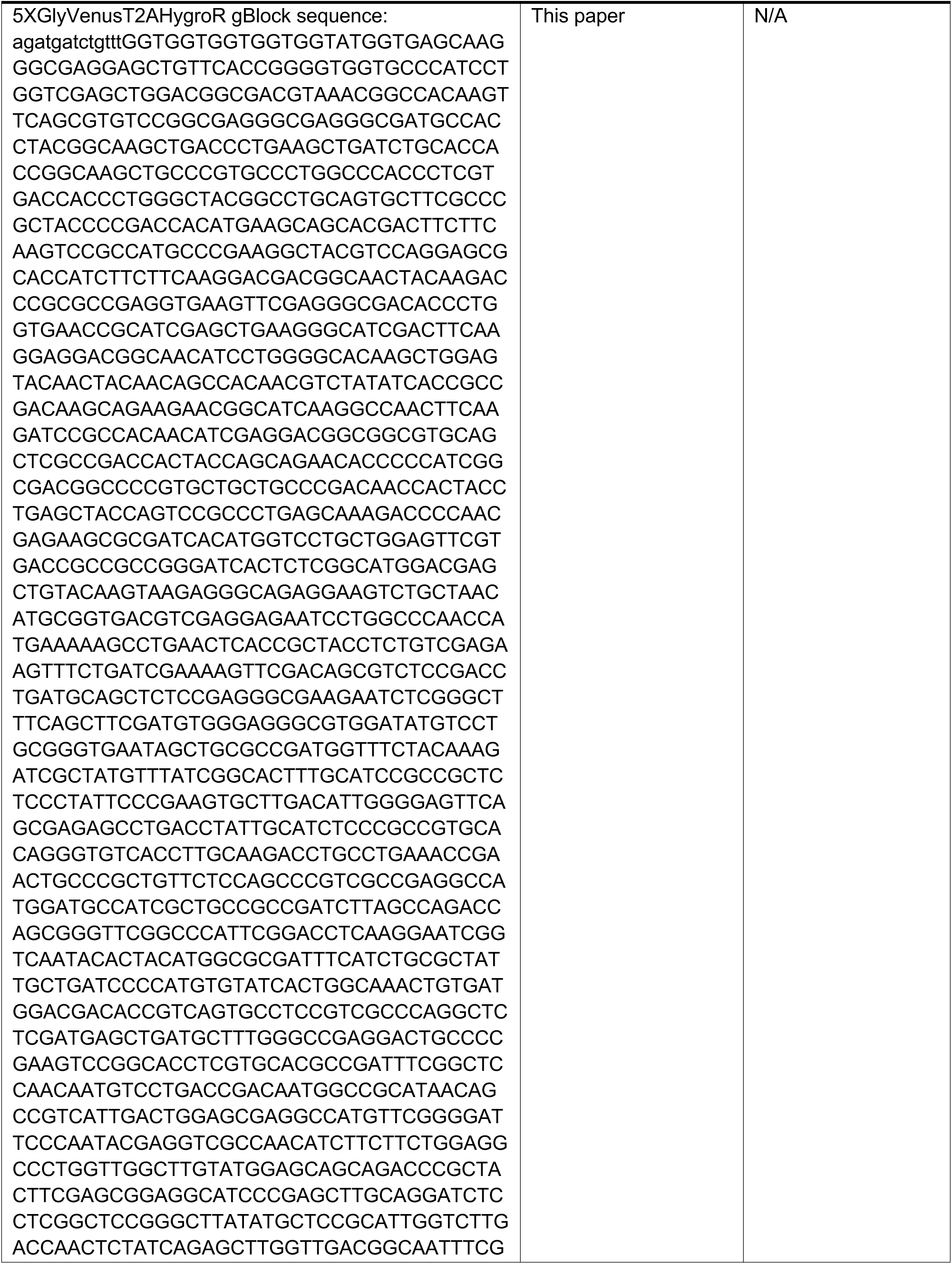
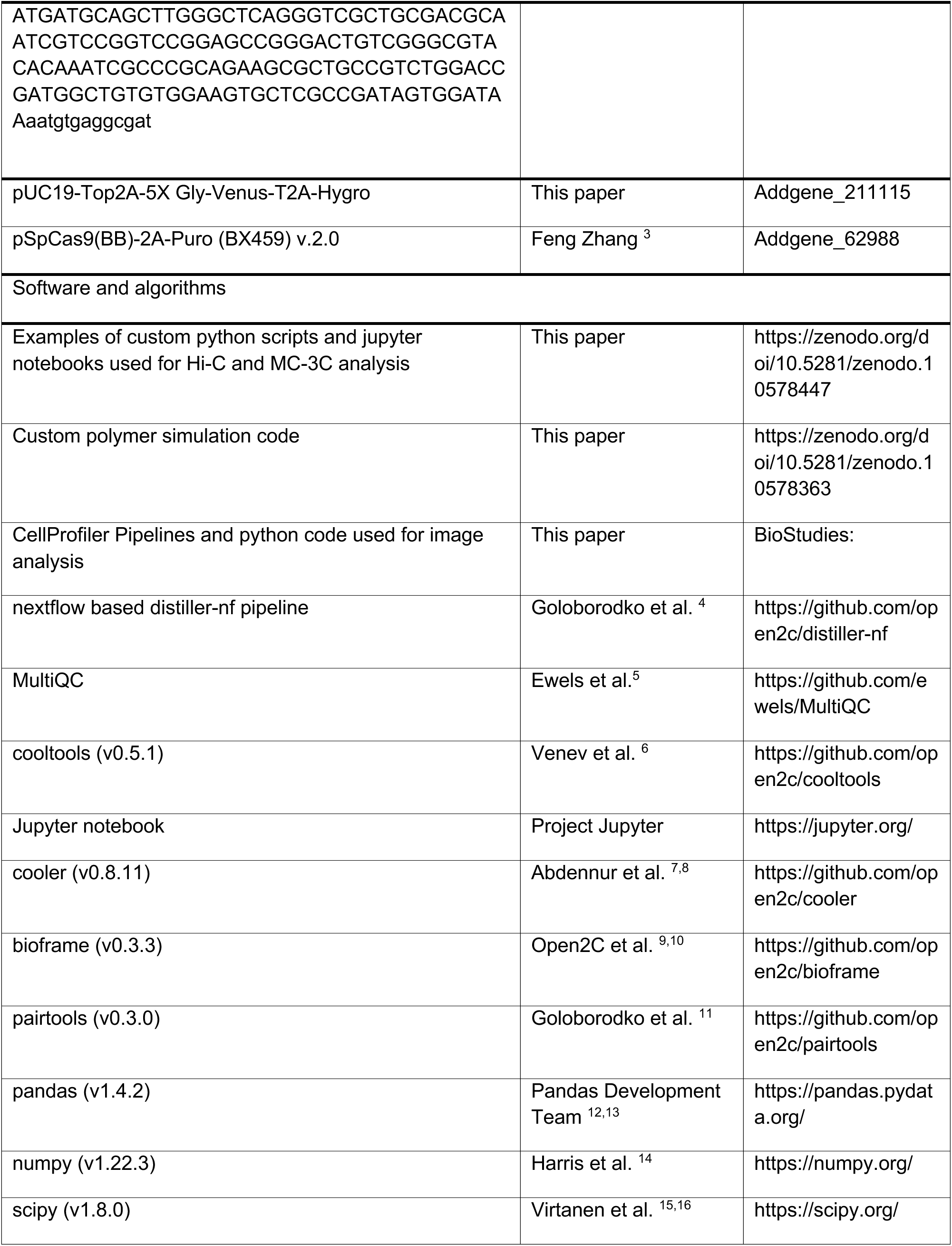
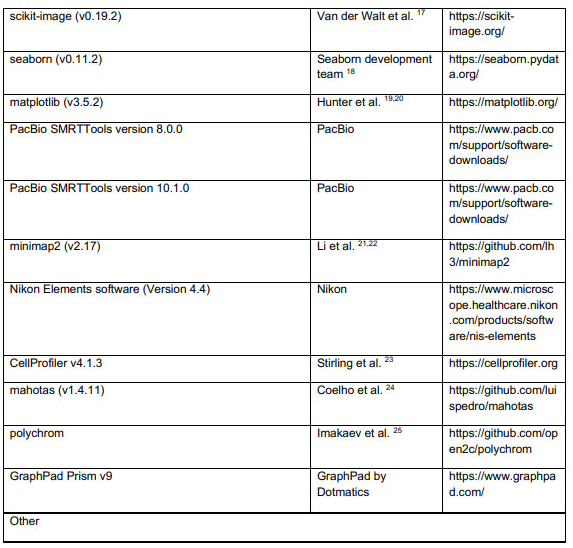

## REFERENCES

1. Pommier, Y., Sun, Y., Huang, S.N., and Nitiss, J.L. (2016). Roles of eukaryotic topoisomerases in transcription, replication and genomic stability. Nat Rev Mol Cell Biol 17, 703–721. 10.1038/nrm.2016.111.

2. Austin, C.A., and Fisher, L.M. (1990). Isolation and characterization of a human cDNA clone encoding a novel DNA topoisomerase II homologue from HeLa cells. FEBS Lett 266, 115–117. 10.1016/0014-5793(90)81520-x.

3. Lewis, C.D., and Laemmli, U.K. (1982). Higher order metaphase chromosome structure: evidence for metalloprotein interactions. Cell 29, 171–181. 10.1016/0092-8674(82)90101-5.

4. Earnshaw, W.C., and Heck, M.M. (1985). Localization of topoisomerase II in mitotic chromosomes. J Cell Biol 100, 1716–1725. 10.1083/jcb.100.5.1716.

5. Earnshaw, W.C., Halligan, B., Cooke, C.A., Heck, M.M., and Liu, L.F. (1985). Topoisomerase II is a structural component of mitotic chromosome scaffolds. J Cell Biol 100, 1706–1715. 10.1083/jcb.100.5.1706.

6. Drake, F.H., Zimmerman, J.P., McCabe, F.L., Bartus, H.F., Per, S.R., Sullivan, D.M., Ross, W.E., Mattern, M.R., Johnson, R.K., Crooke, S.T., and, et al. (1987). Purification of topoisomerase II from amsacrine-resistant P388 leukemia cells. Evidence for two forms of the enzyme. J Biol Chem 262, 16739–16747.

7. Wang, J.C. (2002). Cellular roles of DNA topoisomerases: a molecular perspective. Nat Rev Mol Cell Biol 3, 430–440. 10.1038/nrm831.

8. Austin, C.A., Barot, H., Margerrison, E.E.C., Hayes, M.V., and Fisher, L.M. (1989). Biochemical and immunological characterization of mammalian DNA topoisomerase II. Biochemical Society Transactions 17, 528–529. 10.1042/bst0170528.

9. Tsai-Pflugfelder, M., Liu, L.F., Liu, A.A., Tewey, K.M., Whang-Peng, J., Knutsen, T., Huebner, K., Croce, C.M., and Wang, J.C. (1988). Cloning and sequencing of cDNA encoding human DNA topoisomerase II and localization of the gene to chromosome region 17q21-22. Proc Natl Acad Sci U S A 85, 7177–7181. 10.1073/pnas.85.19.7177.

10. Broderick, R., and Niedzwiedz, W. (2015). Sister chromatid decatenation: bridging the gaps in our knowledge. Cell Cycle 14, 3040–3044. 10.1080/15384101.2015.1078039.

11. Bower, J.J., Karaca, G.F., Zhou, Y., Simpson, D.A., Cordeiro-Stone, M., and Kaufmann, W.K. (2010). Topoisomerase IIalpha maintains genomic stability through decatenation G(2) checkpoint signaling. Oncogene 29, 4787–4799. 10.1038/onc.2010.232.

12. Luo, K., Yuan, J., Chen, J., and Lou, Z. (2009). Topoisomerase IIalpha controls the decatenation checkpoint. Nat Cell Biol 11, 204–210. 10.1038/ncb1828.

13. Samejima, K., Samejima, I., Vagnarelli, P., Ogawa, H., Vargiu, G., Kelly, D.A., de Lima Alves, F., Kerr, A., Green, L.C., Hudson, D.F., et al. (2012). Mitotic chromosomes are compacted laterally by KIF4 and condensin and axially by topoisomerase IIalpha. J Cell Biol 199, 755–770. 10.1083/jcb.201202155.

14. Warburton, P.E., and Earnshaw, W.C. (1997). Untangling the role of DNA topoisomerase II in mitotic chromosome structure and function. Bioessays 19, 97–99. 10.1002/bies.950190203.

15. Moens, P.B., and Earnshaw, W.C. (1989). Anti-topoisomerase II recognizes meiotic chromosome cores. Chromosoma 98, 317–322. 10.1007/BF00292383.

16. Heck, M.M., and Earnshaw, W.C. (1986). Topoisomerase II: A specific marker for cell proliferation. J Cell Biol 103, 2569–2581. 10.1083/jcb.103.6.2569.

17. Heck, M.M., Hittelman, W.N., and Earnshaw, W.C. (1988). Differential expression of DNA topoisomerases I and II during the eukaryotic cell cycle. Proc Natl Acad Sci U S A 85, 1086–1090. 10.1073/pnas.85.4.1086.

18. Gasser, S.M., Laroche, T., Falquet, J., Boy de la Tour, E., and Laemmli, U.K. (1986). Metaphase chromosome structure. Involvement of topoisomerase II. J Mol Biol 188, 613–629. 10.1016/s0022-2836(86)80010-9.

19. Gasser, S.M., and Laemmli, U.K. (1986). The organisation of chromatin loops: characterization of a scaffold attachment site. EMBO J 5, 511–518. 10.1002/j.1460-2075.1986.tb04240.x.

20. Mirkovitch, J., Gasser, S.M., and Laemmli, U.K. (1988). Scaffold attachment of DNA loops in metaphase chromosomes. J Mol Biol 200, 101–109. 10.1016/0022-2836(88)90336-1.

21. Canela, A., Maman, Y., Huang, S.N., Wutz, G., Tang, W., Zagnoli-Vieira, G., Callen, E., Wong, N., Day, A., Peters, J.M., et al. (2019). Topoisomerase II-Induced Chromosome Breakage and Translocation Is Determined by Chromosome Architecture and Transcriptional Activity. Mol Cell 75, 252–266 e258. 10.1016/j.molcel.2019.04.030.

22. Canela, A., Maman, Y., Jung, S., Wong, N., Callen, E., Day, A., Kieffer-Kwon, K.R., Pekowska, A., Zhang, H., Rao, S.S.P., et al. (2017). Genome Organization Drives Chromosome Fragility. Cell 170, 507–521 e518. 10.1016/j.cell.2017.06.034.

23. Canela, A., Sridharan, S., Sciascia, N., Tubbs, A., Meltzer, P., Sleckman, B.P., and Nussenzweig, A. (2016). DNA Breaks and End Resection Measured Genome-wide by End Sequencing. Mol Cell 63, 898–911. 10.1016/j.molcel.2016.06.034.

24. Uuskula-Reimand, L., Hou, H., Samavarchi-Tehrani, P., Rudan, M.V., Liang, M., Medina-Rivera, A., Mohammed, H., Schmidt, D., Schwalie, P., Young, E.J., et al. (2016). Topoisomerase II beta interacts with cohesin and CTCF at topological domain borders. Genome Biol 17, 182. 10.1186/s13059-016-1043-8.

25. Manville, C.M., Smith, K., Sondka, Z., Rance, H., Cockell, S., Cowell, I.G., Lee, K.C., Morris, N.J., Padget, K., Jackson, G.H., and Austin, C.A. (2015). Genome-wide ChIP-seq analysis of human TOP2B occupancy in MCF7 breast cancer epithelial cells. Biol Open 4, 1436–1447. 10.1242/bio.014308.

26. Lieberman-Aiden, E., van Berkum, N.L., Williams, L., Imakaev, M., Ragoczy, T., Telling, A., Amit, I., Lajoie, B.R., Sabo, P.J., Dorschner, M.O., et al. (2009). Comprehensive mapping of long-range interactions reveals folding principles of the human genome. Science 326, 289–293. 10.1126/science.1181369.

27. Tavares-Cadete, F., Norouzi, D., Dekker, B., Liu, Y., and Dekker, J. (2020). Multi-contact 3C reveals that the human genome during interphase is largely not entangled. Nat Struct Mol Biol 27, 1105–1114. 10.1038/s41594-020-0506-5.

28. Goundaroulis, D., Lieberman Aiden, E., and Stasiak, A. (2020). Chromatin Is Frequently Unknotted at the Megabase Scale. Biophys J 118, 2268–2279. 10.1016/j.bpj.2019.11.002.

29. Grosberg, A., Rabin, Y., Havlin, S., and Neer, A. (1993). Crumpled Globule Model of the Three-Dimensional Structure of DNA. Europhysics Letters 23, 373. 10.1209/0295-5075/23/5/012.

30. Grosberg, A.Y., Nechaev, S.K., and Shakhnovich, E.I. (1988). The role of topological constraints in the kinetics of collapse of macromolecules. Journal de Physique 49 (*12*). 10.1051/jphys:0198800490120209500.

31. Polovnikov, K., and Slavov, B. (2023). Topological and nontopological mechanisms of loop formation in chromosomes: Effects on the contact probability. Physical Review E 107. 10.1103/PhysRevE.107.054135.

32. Polovnikov, K.E., Belan, S., Imakaev, M., Brandao, H., and Mirny, L. (2023). Crumpled polymer with loops recapitulates key features of chromosome organization. Physical Review X (Accepted).

33. Kawamura, R., Pope, L.H., Christensen, M.O., Sun, M., Terekhova, K., Boege, F., Mielke, C., Andersen, A.H., and Marko, J.F. (2010). Mitotic chromosomes are constrained by topoisomerase II-sensitive DNA entanglements. J Cell Biol 188, 653–663. 10.1083/jcb.200910085.

34. Rosa, A., Di Stefano, M., and Micheletti, C. (2019). Topological Constraints in Eukaryotic Genomes and How They Can Be Exploited to Improve Spatial Models of Chromosomes. Front Mol Biosci 6, 127. 10.3389/fmolb.2019.00127.

35. Rosa, A., and Everaers, R. (2008). Structure and dynamics of interphase chromosomes. PLoS Comput Biol 4, e1000153. 10.1371/journal.pcbi.1000153.

36. Sikorav, J.L., and Jannink, G. (1994). Kinetics of chromosome condensation in the presence of topoisomerases: a phantom chain model. Biophys J 66, 827–837. 10.1016/s0006-3495(94)80859-8.

37. Nielsen, C.F., Zhang, T., Barisic, M., Kalitsis, P., and Hudson, D.F. (2020). Topoisomerase IIalpha is essential for maintenance of mitotic chromosome structure. Proc Natl Acad Sci U S A 117, 12131–12142. 10.1073/pnas.2001760117.

38. Antonin, W., and Neumann, H. (2016). Chromosome condensation and decondensation during mitosis. Curr Opin Cell Biol 40, 15–22. 10.1016/j.ceb.2016.01.013.

39. Belaghzal, H., Dekker, J., and Gibcus, J.H. (2017). Hi-C 2.0: An optimized Hi-C procedure for high-resolution genome-wide mapping of chromosome conformation. Methods 123, 56–65. 10.1016/j.ymeth.2017.04.004.

40. Abramo, K., Valton, A.L., Venev, S.V., Ozadam, H., Fox, A.N., and Dekker, J. (2019). A chromosome folding intermediate at the condensin-to-cohesin transition during telophase. Nat Cell Biol 21, 1393–1402. 10.1038/s41556-019-0406-2.

41. Naumova, N., Imakaev, M., Fudenberg, G., Zhan, Y., Lajoie, B.R., Mirny, L.A., and Dekker, J. (2013). Organization of the mitotic chromosome. Science 342, 948–953. 10.1126/science.1236083.

42. Iwai, M., Hara, A., Andoh, T., and Ishida, R. (1997). ICRF-193, a catalytic inhibitor of DNA topoisomerase II, delays the cell cycle progression from metaphase, but not from anaphase to the G1 phase in mammalian cells. FEBS Lett 406, 267–270. 10.1016/s0014-5793(97)00282-2.

43. Tanabe, K., Ikegami, Y., Ishida, R., and Andoh, T. (1991). Inhibition of topoisomerase II by antitumor agents bis(2,6-dioxopiperazine) derivatives. Cancer Res 51, 4903–4908.

44. Naughton, C., Avlonitis, N., Corless, S., Prendergast, J.G., Mati, I.K., Eijk, P.P., Cockroft, S.L., Bradley, M., Ylstra, B., and Gilbert, N. (2013). Transcription forms and remodels supercoiling domains unfolding large-scale chromatin structures. Nat Struct Mol Biol 20, 387–395. 10.1038/nsmb.2509.

45. Zhang, H., Emerson, D.J., Gilgenast, T.G., Titus, K.R., Lan, Y., Huang, P., Zhang, D., Wang, H., Keller, C.A., Giardine, B., et al. (2019). Chromatin structure dynamics during the mitosis-to-G1 phase transition. Nature 576, 158–162. 10.1038/s41586-019-1778-y.

46. Gibcus, J.H., Samejima, K., Goloborodko, A., Samejima, I., Naumova, N., Nuebler, J., Kanemaki, M.T., Xie, L., Paulson, J.R., Earnshaw, W.C., et al. (2018). A pathway for mitotic chromosome formation. Science 359. 10.1126/science.aao6135.

47. Rao, S.S., Huntley, M.H., Durand, N.C., Stamenova, E.K., Bochkov, I.D., Robinson, J.T., Sanborn, A.L., Machol, I., Omer, A.D., Lander, E.S., and Aiden, E.L. (2014). A 3D map of the human genome at kilobase resolution reveals principles of chromatin looping. Cell 159, 1665–1680. 10.1016/j.cell.2014.11.021.

48. Hildebrand, E.M., and Dekker, J. (2020). Mechanisms and Functions of Chromosome Compartmentalization. Trends Biochem Sci 45, 385–396. 10.1016/j.tibs.2020.01.002.

49. Solovei, I., Kreysing, M., Lanctot, C., Kosem, S., Peichl, L., Cremer, T., Guck, J., and Joffe, B. (2009). Nuclear architecture of rod photoreceptor cells adapts to vision in mammalian evolution. Cell 137, 356–368. 10.1016/j.cell.2009.01.052.

50. Poleshko, A., Smith, C.L., Nguyen, S.C., Sivaramakrishnan, P., Wong, K.G., Murray, J.I., Lakadamyali, M., Joyce, E.F., Jain, R., and Epstein, J.A. (2019). H3K9me2 orchestrates inheritance of spatial positioning of peripheral heterochromatin through mitosis. Elife 8. 10.7554/eLife.49278.

51. Falk, M., Feodorova, Y., Naumova, N., Imakaev, M., Lajoie, B.R., Leonhardt, H., Joffe, B., Dekker, J., Fudenberg, G., Solovei, I., and Mirny, L.A. (2019). Heterochromatin drives compartmentalization of inverted and conventional nuclei. Nature 570, 395–399. 10.1038/s41586-019-1275-3.

52. Smith, C.L., Lan, Y., Jain, R., Epstein, J.A., and Poleshko, A. (2021). Global chromatin relabeling accompanies spatial inversion of chromatin in rod photoreceptors. Sci Adv 7, eabj3035. 10.1126/sciadv.abj3035.

53. Xu, J., Ma, H., Jin, J., Uttam, S., Fu, R., Huang, Y., and Liu, Y. (2018). Super-Resolution Imaging of Higher-Order Chromatin Structures at Different Epigenomic States in Single Mammalian Cells. Cell Rep 24, 873–882. 10.1016/j.celrep.2018.06.085.

54. Haralick, R.M., Shanmugam, K., and Dinstein, I.H. (1973). Textural Features for Image Classification. IEEE Transactions on Systems, Man, and Cybernetics SMC*-*3, 610–621. 10.1109/tsmc.1973.4309314.

55. Bensaude, O. (2011). Inhibiting eukaryotic transcription: Which compound to choose? How to evaluate its activity? Transcription 2, 103–108. 10.4161/trns.2.3.16172.

56. Polovnikov, K., Nechaev, S., and Tamm, M.V. (2018). Effective Hamiltonian of topologically stabilized polymer states. Soft Matter 14, 6561–6570. 10.1039/c8sm00785c.

57. Polovnikov, K.E., Nechaev, S., and Tamm, M.V. (2019). Many-body contacts in fractal polymer chains and fractional Brownian trajectories. Phys Rev E 99, 032501. 10.1103/PhysRevE.99.032501.

58. Halverson, J.D., Lee, W.B., Grest, G.S., Grosberg, A.Y., and Kremer, K. (2011). Molecular dynamics simulation study of nonconcatenated ring polymers in a melt. I. Statics. J Chem Phys 134, 204904. 10.1063/1.3587137.

59. Halverson, J.D., Smrek, J., Kremer, K., and Grosberg, A.Y. (2014). From a melt of rings to chromosome territories: the role of topological constraints in genome folding. Rep Prog Phys 77, 022601. 10.1088/0034-4885/77/2/022601.

60. Rosa, A., Becker, N.B., and Everaers, R. (2010). Looping probabilities in model interphase chromosomes. Biophys J 98, 2410–2419. 10.1016/j.bpj.2010.01.054.

61. Rao, S.S.P., Huang, S.C., Glenn St Hilaire, B., Engreitz, J.M., Perez, E.M., Kieffer-Kwon, K.R., Sanborn, A.L., Johnstone, S.E., Bascom, G.D., Bochkov, I.D., et al. (2017). Cohesin Loss Eliminates All Loop Domains. Cell 171, 305–320 e324. 10.1016/j.cell.2017.09.026.

62. Hsieh, T.-H.S., Cattoglio, C., Slobodyanyuk, E., Hansen, A.S., Darzacq, X., and Tjian, R. (2021). Enhancer-promoter interactions and transcription are maintained upon acute loss of CTCF, cohesin, WAPL, and YY1. bioRxiv 2021.07.*14.452365*. doi: 10.1101/2021.07.14.452365.

63. Mora-Bermudez, F., Gerlich, D., and Ellenberg, J. (2007). Maximal chromosome compaction occurs by axial shortening in anaphase and depends on Aurora kinase. Nat Cell Biol 9, 822–831. 10.1038/ncb1606.

64. Grosberg, A.Y., and Khokhlov, A.R. (1994). Stastical Physics of Macromolecules (AIP Press).

65. Halverson, J.D., Lee, W.B., Grest, G.S., Grosberg, A.Y., and Kremer, K. (2011). Molecular dynamics simulation study of nonconcatenated ring polymers in a melt. II. Dynamics. J Chem Phys 134, 204905. 10.1063/1.3587138.

66. Nishimura, K., Fukagawa, T., Takisawa, H., Kakimoto, T., and Kanemaki, M. (2009). An auxin-based degron system for the rapid depletion of proteins in nonplant cells. Nat Methods 6, 917–922. 10.1038/nmeth.1401.

67. Natsume, T., Kiyomitsu, T., Saga, Y., and Kanemaki, M.T. (2016). Rapid Protein Depletion in Human Cells by Auxin-Inducible Degron Tagging with Short Homology Donors. Cell Rep 15, 210–218. 10.1016/j.celrep.2016.03.001.

68. Liu, Y., and Dekker, J. (2021). Biochemically distinct cohesin complexes mediate positioned loops between CTCF sites and dynamic loops within chromatin domains. bioRxiv 2021*.08.24.457555*. doi: 10.1101/2021.08.24.457555.

69. Haarhuis, J.H.I., van der Weide, R.H., Blomen, V.A., Yanez-Cuna, J.O., Amendola, M., van Ruiten, M.S., Krijger, P.H.L., Teunissen, H., Medema, R.H., van Steensel, B., et al. (2017). The Cohesin Release Factor WAPL Restricts Chromatin Loop Extension. Cell 169, 693–707 e614. 10.1016/j.cell.2017.04.013.

70. Schwarzer, W., Abdennur, N., Goloborodko, A., Pekowska, A., Fudenberg, G., Loe-Mie, Y., Fonseca, N.A., Huber, W., Haering, C.H., Mirny, L., and Spitz, F. (2017). Two independent modes of chromatin organization revealed by cohesin removal. Nature 551, 51–56. 10.1038/nature24281.

71. Shintomi, K., and Hirano, T. (2021). Guiding functions of the C-terminal domain of topoisomerase IIalpha advance mitotic chromosome assembly. Nat Commun 12, 2917. 10.1038/s41467-021-23205-w.

72. Walther, N., Hossain, M.J., Politi, A.Z., Koch, B., Kueblbeck, M., Odegard-Fougner, O., Lampe, M., and Ellenberg, J. (2018). A quantitative map of human Condensins provides new insights into mitotic chromosome architecture. J Cell Biol 217, 2309–2328. 10.1083/jcb.201801048.

73. Gerlich, D., Hirota, T., Koch, B., Peters, J.M., and Ellenberg, J. (2006). Condensin I stabilizes chromosomes mechanically through a dynamic interaction in live cells. Curr Biol 16, 333–344. 10.1016/j.cub.2005.12.040.

74. Bintu, B., Mateo, L.J., Su, J.H., Sinnott-Armstrong, N.A., Parker, M., Kinrot, S., Yamaya, K., Boettiger, A.N., and Zhuang, X. (2018). Super-resolution chromatin tracing reveals domains and cooperative interactions in single cells. Science 362. 10.1126/science.aau1783.

75. Ran, F.A., Hsu, P.D., Wright, J., Agarwala, V., Scott, D.A., and Zhang, F. (2013). Genome engineering using the CRISPR-Cas9 system. Nat Protoc 8, 2281–2308. 10.1038/nprot.2013.143.

76. Hsieh, T.S., Fudenberg, G., Goloborodko, A., and Rando, O.J. (2016). Micro-C XL: assaying chromosome conformation from the nucleosome to the entire genome. Nat Methods 13, 1009–1011. 10.1038/nmeth.4025.

77. Krietenstein, N., Abraham, S., Venev, S.V., Abdennur, N., Gibcus, J., Hsieh, T.S., Parsi, K.M., Yang, L., Maehr, R., Mirny, L.A., et al. (2020). Ultrastructural Details of Mammalian Chromosome Architecture. Mol Cell 78, 554–565 e557. 10.1016/j.molcel.2020.03.003.

78. Krietenstein, N., and Rando, O.J. (2022). Mammalian Micro-C-XL. Methods Mol Biol 2458, 321–332. 10.1007/978-1-0716-2140-0_17.

79. Sarkans, U., Gostev, M., Athar, A., Behrangi, E., Melnichuk, O., Ali, A., Minguet, J., Rada, J.C., Snow, C., Tikhonov, A., et al. (2018). The BioStudies database-one stop shop for all data supporting a life sciences study. Nucleic Acids Res 46, D1266–D1270. 10.1093/nar/gkx965.

80. Eastman, P., and Pande, V.S. (2015). OpenMM: A Hardware Independent Framework for Molecular Simulations. Comput Sci Eng 12, 34–39. 10.1109/MCSE.2010.27.

81. Arbona, J.M., Herbert, S., Fabre, E., and Zimmer, C. (2017). Inferring the physical properties of yeast chromatin through Bayesian analysis of whole nucleus simulations. Genome Biol 18, 81. 10.1186/s13059-017-1199-x.

82. Doi, M., and Edwards, S.F. (1988). The Theory of Polymer Dynamics (Oxford University Press).

83. Kremer, K., and Grest, G. (1990). Dynamics of entangled linear polymer melts: A molecular-dynamics simulation. The Journal of Chemical Physics 92.

84. Nuebler, J., Fudenberg, G., Imakaev, M., Abdennur, N., and Mirny, L.A. (2018). Chromatin organization by an interplay of loop extrusion and compartmental segregation. Proc Natl Acad Sci U S A 115, E6697–E6706. 10.1073/pnas.1717730115.

85. Davidson, I.F., Bauer, B., Goetz, D., Tang, W., Wutz, G., and Peters, J.M. (2019). DNA loop extrusion by human cohesin. Science 366, 1338–1345. 10.1126/science.aaz3418.

86. Golfier, S., Quail, T., Kimura, H., and Brugues, J. (2020). Cohesin and condensin extrude DNA loops in a cell cycle-dependent manner. Elife 9. 10.7554/eLife.53885.

87. Kim, Y., Shi, Z., Zhang, H., Finkelstein, I.J., and Yu, H. (2019). Human cohesin compacts DNA by loop extrusion. Science 366, 1345–1349. 10.1126/science.aaz4475.

88. Polovnikov, K.E., Gherardi, M., Cosentino-Lagomarsino, M., and Tamm, M.V. (2018). Fractal Folding and Medium Viscoelasticity Contribute Jointly to Chromosome Dynamics. Phys Rev Lett 120, 088101. 10.1103/PhysRevLett.120.088101.

89. Tamm, M.V., Nazarov, L.I., Gavrilov, A.A., and Chertovich, A.V. (2015). Anomalous diffusion in fractal globules. Physical review letters, 114, 178102.

90. Ge, T., Panyukov, S., and Rubinstein, M. (2016). Self-similar conformations and dynamics in entangled melts and solutions of nonconcatenated ring polymers. Macromolecules 49, 708–722.

91. Smrek, J., and Grosberg, A.Y. (2015). Understanding the dynamics of rings in the melt in terms of the annealed tree model. Journal of Physics: Condensed Matter 27.

92. Walther, N., and Ellenberg, J. (2018). Quantitative live and super-resolution microscopy of mitotic chromosomes. Methods Cell Biol 145, 65–90. 10.1016/bs.mcb.2018.03.014.

93. Goloborodko, A., Venev, S., Abdennur, N., azkalot1, and Tommaso, P.D. (2019). mirnylab/distiller-nf: v0.3.3 (v0.3.3). Zenodo. 10.5281/zenodo.3350937.

94. Venev, S., Abdennur, N., Goloborodko, A., Flyamer, I., Fudenberg, G., Nuebler, J., Galitsyna, A., Akgol, B., Abraham, S., Kerpedjiev, P., and Imakaev, M. (2022). open2c/cooltools: v0.5.1. Zenodo. 10.5281/zenodo.6324229.

95. Ewels, P., Magnusson, M., Lundin, S., and Kaller, M. (2016). MultiQC: summarize analysis results for multiple tools and samples in a single report. Bioinformatics 32, 3047–3048. 10.1093/bioinformatics/btw354.

96. Abdennur, N., Goloborodko, A., Imakaev, M., Kerpedjiev, P., Fudenberg, G., Oullette, S., Lee, S., Strobelt, H., Gehlenborg, N., and Mirny, L. (2021). open2c/cooler: v0.8.11 (v0.8.11). Zenodo. 10.5281/zenodo.4655850.

97. Abdennur, N., and Mirny, L.A. (2020). Cooler: scalable storage for Hi-C data and other genomically labeled arrays. Bioinformatics 36, 311–316. 10.1093/bioinformatics/btz540.

98. Abdennur, N., Goloborodko, A., gfudenberg, Imakaev, M., agalitsyna, Venev, S., Abraham, S., Flyamer, I., Spracklin, G., Chumpitaz, L., and Aafke (2022). open2c/bioframe: v0.3.3. Zenodo. 10.5281/zenodo.6317259.

99. Abdennur, N., Goloborodko, A., gfudenberg, Imakaev, M., Galitsyna, A., Abraham, S., Spracklin, G., Venev, S., Chumpitaz, L., Flyamer, I., and Aafke (2021). open2c/bioframe: v0.3.0 (v0.3.0). Zenodo. 10.5281/zenodo.5348312.

100. Goloborodko, A., Abdennur, N., Venev, S., hbbrandao, and gfudenberg (2019). mirnylab/pairtools v0.3.0 (v0.3.0). Zenodo. 10.5281/zenodo.2649383.

101. Reback, J., jbrockmendel, McKinney, W., Bossche, J.V.d., Augspurger, T., Roeschke, M., Hawkins, S., Cloud, P., gfyoung, Sinhrks, et al. (2022). pandas-dev/pandas: Pandas 1.4.2 (v1.4.2). Zenodo. 10.5281/zenodo.6408044.

102. Harris, C.R., Millman, K.J., van der Walt, S.J., Gommers, R., Virtanen, P., Cournapeau, D., Wieser, E., Taylor, J., Berg, S., Smith, N.J., et al. (2020). Array programming with NumPy. Nature 585, 357–362. 10.1038/s41586-020-2649-2.

103. team, T.s.d. (2021). scipy/scipy: SciPy 1.7.1 (v1.7.1). Zenodo. 10.5281/zenodo.5152559.

104. Virtanen, P., Gommers, R., Oliphant, T.E., Haberland, M., Reddy, T., Cournapeau, D., Burovski, E., Peterson, P., Weckesser, W., Bright, J., et al. (2020). SciPy 1.0: fundamental algorithms for scientific computing in Python. Nat Methods 17, 261–272. 10.1038/s41592-019-0686-2.

105. Hunter, J.D. (2007). Matplotlib: A 2D Graphics Environment. Computing in Science & Engineering 9, 90–95. 10.1109/mcse.2007.55.

106. team, T.m.d. (2021). matplotlib/matplotlib: REL: v3.4.3 (v3.4.3). Zenodo. 10.5281/zenodo.5194481.

107. van der Walt, S., Schönberger, J., Nunez-Iglesias, J., Boulogne, F., Warner, J., Yager, N., Gouillart, E., Yu, T., and contributors., t.s.-i. (2014). scikit-image: image processing in Python. PeerJ 2. 10.7717/peerj.453.

108. team, T.s.d. (2021). mwaskom/seaborn: v0.11.2 (August 2021) (v0.11.2). Zenodo. 10.5281/zenodo.5205191.

109. Imakaev, M., Fudenberg, G., McCord, R.P., Naumova, N., Goloborodko, A., Lajoie, B.R., Dekker, J., and Mirny, L.A. (2012). Iterative correction of Hi-C data reveals hallmarks of chromosome organization. Nat Methods 9, 999–1003. 10.1038/nmeth.2148.

110. Venev, S., Abdennur, N., Goloborodko, A., Flyamer, I., Fudenberg, G., Nuebler, J., Galitsyna, A., Akgol, B., Abraham, S., Kerpedjiev, P., and Imakaev, M. (2021). open2c/cooltools: v0.4.0 (v0.4.0). Zenodo. 10.5281/zenodo.4667696.

111. Li, H. (2021). New strategies to improve minimap2 alignment accuracy. Bioinformatics. 10.1093/bioinformatics/btab705.

112. Li, H. (2018). Minimap2: pairwise alignment for nucleotide sequences. Bioinformatics 34, 3094–3100. 10.1093/bioinformatics/bty191.

113. Stirling, D.R., Swain-Bowden, M.J., Lucas, A.M., Carpenter, A.E., Cimini, B.A., and Goodman, A. (2021). CellProfiler 4: improvements in speed, utility and usability. BMC Bioinformatics 22, 433. 10.1186/s12859-021-04344-9.

## References

1. Rao, S.S.P., Huang, S.C., Glenn St Hilaire, B., Engreitz, J.M., Perez, E.M., Kieffer-Kwon, K.R., Sanborn, A.L., Johnstone, S.E., Bascom, G.D., Bochkov, I.D., et al. (2017). Cohesin Loss Eliminates All Loop Domains. Cell 171, 305–320 e324. 10.1016/j.cell.2017.09.026.

2. Natsume, T., Kiyomitsu, T., Saga, Y., and Kanemaki, M.T. (2016). Rapid Protein Depletion in Human Cells by Auxin-Inducible Degron Tagging with Short Homology Donors. Cell Rep 15, 210–218. 10.1016/j.celrep.2016.03.001.

3. Ran, F.A., Hsu, P.D., Wright, J., Agarwala, V., Scott, D.A., and Zhang, F. (2013). Genome engineering using the CRISPR-Cas9 system. Nat Protoc 8, 2281–2308. 10.1038/nprot.2013.143.

4. Goloborodko, A., Venev, S., Abdennur, N., azkalot1, and Tommaso, P.D. (2019). mirnylab/distiller-nf: v0.3.3 (v0.3.3). Zenodo. 10.5281/zenodo.3350937.

5. Ewels, P., Magnusson, M., Lundin, S., and Kaller, M. (2016). MultiQC: summarize analysis results for multiple tools and samples in a single report. Bioinformatics 32, 3047–3048. 10.1093/bioinformatics/btw354.

6. Venev, S., Abdennur, N., Goloborodko, A., Flyamer, I., Fudenberg, G., Nuebler, J., Galitsyna, A., Akgol, B., Abraham, S., Kerpedjiev, P., and Imakaev, M. (2022). open2c/cooltools: v0.5.1. Zenodo. 10.5281/zenodo.6324229.

7. Abdennur, N., Goloborodko, A., Imakaev, M., Kerpedjiev, P., Fudenberg, G., Oullette, S., Lee, S., Strobelt, H., Gehlenborg, N., and Mirny, L. (2021). open2c/cooler: v0.8.11 (v0.8.11). Zenodo. 10.5281/zenodo.4655850.

8. Abdennur, N., and Mirny, L.A. (2020). Cooler: scalable storage for Hi-C data and other genomically labeled arrays. Bioinformatics 36, 311–316. 10.1093/bioinformatics/btz540.

9. Abdennur, N., Goloborodko, A., gfudenberg, Imakaev, M., agalitsyna, Venev, S., Abraham, S., Flyamer, I., Spracklin, G., Chumpitaz, L., and Aafke (2022). open2c/bioframe: v0.3.3. Zenodo. 10.5281/zenodo.6317259.

10. Open2C, Abdennur, N., Fudenberg, G., Flyamer, I., Galitsyna, A.A., Goloborodko, A., Imakaev, M., and Venev, S.V. (2022). Bioframe: Operations on Genomic Intervals in Pandas Dataframes. bioRxiv 2022.02.16.480748. 10.1101/2022.02.16.480748.

11. Goloborodko, A., Abdennur, N., Venev, S., hbbrandao, and gfudenberg (2019). mirnylab/pairtools v0.3.0 (v0.3.0). Zenodo. 10.5281/zenodo.2649383.

12. Reback, J., jbrockmendel, McKinney, W., Bossche, J.V.d., Augspurger, T., Roeschke, M., Hawkins, S., Cloud, P., gfyoung, Sinhrks, et al. (2022). pandas-dev/pandas: Pandas 1.4.2 (v1.4.2). Zenodo. 10.5281/zenodo.6408044.

13. team, T.p.d. (2021). pandas-dev/pandas: Pandas 1.3.2 (v1.3.2). Zenodo. 10.5281/zenodo.5203279.

14. Harris, C.R., Millman, K.J., van der Walt, S.J., Gommers, R., Virtanen, P., Cournapeau, D., Wieser, E., Taylor, J., Berg, S., Smith, N.J., et al. (2020). Array programming with NumPy. Nature 585, 357–362. 10.1038/s41586-020-2649-2.

15. team, T.s.d. (2021). scipy/scipy: SciPy 1.7.1 (v1.7.1). Zenodo. 10.5281/zenodo.5152559.

16. Virtanen, P., Gommers, R., Oliphant, T.E., Haberland, M., Reddy, T., Cournapeau, D., Burovski, E., Peterson, P., Weckesser, W., Bright, J., et al. (2020). SciPy 1.0: fundamental algorithms for scientific computing in Python. Nat Methods 17, 261–272. 10.1038/s41592-019-0686-2.

17. van der Walt, S., Schönberger, J., Nunez-Iglesias, J., Boulogne, F., Warner, J., Yager, N., Gouillart, E., Yu, T., and contributors., t.s.-i. (2014). scikit-image: image processing in Python. PeerJ 2. 10.7717/peerj.453.

18. team, T.s.d. (2021). mwaskom/seaborn: v0.11.2 (August 2021) (v0.11.2). Zenodo. 10.5281/zenodo.5205191.

19. Hunter, J.D. (2007). Matplotlib: A 2D Graphics Environment. Computing in Science & Engineering 9, 90–95. 10.1109/mcse.2007.55.

20. team, T.m.d. (2021). matplotlib/matplotlib: REL: v3.4.3 (v3.4.3). Zenodo. 10.5281/zenodo.5194481.

21. Li, H. (2021). New strategies to improve minimap2 alignment accuracy. Bioinformatics. 10.1093/bioinformatics/btab705.

22. Li, H. (2018). Minimap2: pairwise alignment for nucleotide sequences. Bioinformatics 34, 3094–3100. 10.1093/bioinformatics/bty191.

23. Stirling, D.R., Swain-Bowden, M.J., Lucas, A.M., Carpenter, A.E., Cimini, B.A., and Goodman, A. (2021). CellProfiler 4: improvements in speed, utility and usability. BMC Bioinformatics 22, 433. 10.1186/s12859-021-04344-9.

24. Coelho, L.P. (2013). Mahotas: Open source software for scriptable computer vision. Journal of Open Research Software 1, e3. 10.5334/jors.ac.

25. Imakaev, M., Goloborodko, A., and Brandao, H. (2019). mirnylab/polychrom: v0.1.0 (v0.1.0). Zenodo. 10.5281/zenodo.3579473.

